# OmicsView: omics data analysis through interactive visual analytics

**DOI:** 10.1101/2021.09.15.460469

**Authors:** Fergal Casey, Soumya Negi, Jing Zhu, Maria Zavodszky, Derrick Cheng, Dongdong Lin, Sally John, Michelle A. Penny, David Sexton, Baohong Zhang

**Author notes:** Contributed to the work equally (Fergal Casey, Soumya Negi).

## Abstract

With advances in NGS technologies, transcriptional profiling of human tissue across many diseases is becoming more routine, leading to the generation of petabytes of data deposited in public repositories. There is a need for bench scientists with little computational expertise to be able to access and mine this data to understand disease pathology, identify robust biomarkers of disease and the effect of interventions (in vivo or in vitro). To this end we release an open source analytics and visualization platform for expression data called OmicsView, http://omicsview.org.

This platform comes preloaded with 1000s of samples across many disease areas and normal tissue, including the GTEx database, all processed with a harmonized pipeline. We demonstrate the power and ease-of-use of the platform by means of a Crohn’s disease data mining exercise where we can quickly uncover disease pathology and identify strong biomarkers of disease and response to treatment.

## INTRODUCTION

Comprehensive omics profiling of patients is fundamental for understanding the molecular mechanism of diseases as well as drug discovery in the era of precision medicine. Modern technologies have enabled the generation of terabytes of RNA-seq and microarray data across all disease areas, and typically they are deposited into public repositories from both academia and industry. However, the deposited data are heterogeneous, and analysed using highly customized pipelines against a variety of annotations, making it a formidable task to make comparisons across projects and diseases. Interpretation and visualization of high-dimensional omics data sets is another daunting task that requires considerable computational training and practice. Although web portals have been developed to analyze publicly available and proprietary datasets^1–4^, the analytical capacity is either limited or specialized to the disease of interest, e.g. oncology. Thanks to major breakthroughs in data generation, especially Next Generation Sequencing (NGS), there is a booming demand for advanced tools providing comprehensive visualizations as well as analytical capabilities for biological interpretation of data. Here, we make the public release of OmicsView (http://omicsview.org), an open source solution that is designed to be data type agnostic with all visualization modules easily adaptable to new data types, and has a straightforward user data upload process. Furthermore, we provide sample level gene expression data sets across ten disease areas from a curation effort in collaboration with Qiagen to demonstrate the versatile functionalities of the system with an emphasis on accessibility of advanced visual and analytical capabilities and cross study meta-analyses. With OmicsView, every bench scientist, regardless of their computational skill level, can carry out in-depth data analysis and interpretation.

## MATERIAL AND METHODS

OmicsView was developed as an independent interactive web application with backend database support that uses multiple underlying libraries and tools for visualizations. The interactive plots have been generated with a diverse array of JavaScript libraries like CanvasXpress, D3, Plotly and Highcharts running within the web browser. Additionally, R packages, specifically from Bioconductor have been used to create some specific plots like KEGG pathways with a gene highlighting feature. JavaScript and PHP were the underlying programming languages for the web framework and to tie the visualizations together. Finally, MySQL was used for data storage and management.

### Data processing

To demonstrate the utility of the web-based application, a subset of curated data from DiseaseLand (https://www.qiagenbioinformatics.com/diseaseland/) is accessible through OmicsView. The DiseaseLand data service uses common analysis pipelines to quantify and normalize publicly available microarray and RNA-seq expression data from raw files. For each project, and each sample, metadata are curated to apply controlled vocabularies and ensure consistent formatting of metadata fields. Information on the data processing pipeline and the DiseaseLand product is available at the following links: http://www.arrayserver.com/wiki/index.php?title=DiseaseLand_Curation_Pipeline http://www.arrayserver.com/wiki/index.php?title=Omicsoft_Affymetrix_Microarray_Preprocessing http://www.arrayserver.com/wiki/index.php?title=RNA-Seq_Normalized_FPKM_Values_in_Land. The data in OmicsView was exported from DiseaseLand prior to 8/27/2019. None of the preloaded curated data can be exported from the portal, distributed with a software package, or otherwise redistributed without express permission from QIAGEN.

### Plotting and Visualization

OmicsView offers two main ways to visualize datasets: through a gene-centric or a dataset-centric way. The gene-centric way shows expression of a single or multiple candidate genes across the datasets available in OmicsView (see Supplemental text: Section 2. Visualize Gene expression). On the other hand, the dataset-centric way is based on one or more candidate datasets, and plots expression of all genes within them (see Supplemental text: Section 3. Visualize Comparison data). OmicsView offers a variety of plotting types like Bubble Plots, PCA Plots, Volcano Plots, Heatmaps, Pathway plots etc, that are highly customizable.. These visualization options are data-type agnostic enabling plotting of micro-array, RNA-seq and proteomic datasets together for comparative analysis (Supplemental text, section 3.12). This is possible because of the uniform way the datasets are analysed, normalized and stored in OmicsView.

### Functional and Pathway Enrichment

The functional enrichment algorithms implemented in OmicsView come in two flavors. The first is based on hypergeometric enrichment of differentially expressed genes (DEGs) against a number of knowledge databases, based on the HOMER package (http://homer.salk.edu/homer/microarray/go.html). This requires a significance cutoff to be applied to the two group comparison expression data in order to select a set of genes that is upregulated and downregulated, i.e. differentially expressed genes. The HOMER package comes with pre-compiled collections of gene knowledge databases that we use for enrichment. The second approach is based on a variant of the Gene Set Enrichment Analysis (GSEA) approach called PAGE (Parametric Analysis of Gene Set Enrichment) which is faster than the original GSEA approach and more sensitive^5^. We use an implementation of PAGE included in the R piano package (https://www.bioconductor.org/packages/release/bioc/html/piano.html). GSEA based approaches just require a ranked list of genes (genes with a numeric quantity for ranking them that shows how different they are between groups in the comparison, usually logFC values) and are typically more sensitive in detecting pathway enrichment. Both, HOMER and PAGE packages come pre-built with a set of known functions/pathways in the form of gene sets, against which we look for enrichment in the comparison of interest. The enrichment tools are run against differential expression data as they are being imported into OmicsView, so the end user is presented with pre-computed results readily. We describe both enrichment approaches in the subsequent sections.

### HOMER workflow

The HOMER workflow is our first enrichment tool that enables pathway analysis on differentially expressed genes (DEGs) in two ways. The first step is selection of differentially expressed genes (DEGs) for enrichment analysis. For each comparison, a list of the up- and down-regulated genes are selected as input for functional enrichment analysis. We use a dynamic cutoff of logFC and AdjustedPValue / PValue to aim for 200-2000 genes in each list. To achieve this, we start with a stringent cutoff of adjusted p-value of 0.05, and 2-fold change up or down. Getting a list of greater than 200 genes is ideal for the downstream analysis. If there are fewer, we drop the nominal p-value to 0.01 (with 2-fold up or down), and subsequently if 200 is not reached we drop the fold change to 1.2 in either direction or increase the adjusted p-value to 0.1. If the most lenient cutoff does not generate a list of 50 up or down genes, we take the top 50 logFC (negative and positive) to generate the lists for enrichment analysis. The second step is functional enrichment of the DEG list against the genome. In this workflow the findGo.pl function from the Homer package implements the hypergeometric test and is used to analyse the functional enrichment for each DEG list. There are several different “ontologies”, or libraries of gene groupings that come with the Homer package. We used Homer version v4.8.3, human-o v5.8 and mouse-o v5.8 libraries (current as of Dec 6, 2016). The HTML output from HOMER is further enhanced by a custom php script for a cleaner display of browsable results.

### Gene Set Enrichment Analysis (GSEA) and PAGE workflow

For our second enrichment tool, we used a variation of the GSEA method, PAGE (Parametric Analysis of Gene Set Enrichment)^5^ to process the DEG data, as PAGE is much faster than GSEA and more sensitive. For each comparison, we produce a rank file with gene symbol and logFC values. If a gene symbol appears multiple times in the same comparison, the average logFC is used. Human gene sets were downloaded from MSigDB, version 5.2 (msigdb.v5.2.symbols.gmt). Mouse gene sets were downloaded from Bader Lab from Univ. of Toronto (http://baderlab.org/GeneSets), version from December 2016 (Mouse_GO_AllPathways_with_GO_iea_December_01_2016_symbol.gmt). This gmt file contains special characters that cannot be used by the R piano package, therefore we manually replace special characters to / or _. In addition, some mouse gene sets have the same name, so we added suffix _altSetX to make all the names unique. For the PAGE workflow, the R piano package is used for all the rank files. To simplify the piano output, we combined down-regulated and up-regulated gene sets into a single table for each gene set. From the piano results we extract out the p-value, FDR and Z-score.

### Meta-Analysis

The meta-analysis functions allow a user to combine expression data across multiple studies to find changes that are robust and much less likely to be due to batch differences. OmicsView offers two ways to perform meta-analysis. The first method works on comparison data. The system will use the comparison data (logFC, p-value) to compute a combined p-value and rank product (product of ranks for each gene across the comparisons of interest). This method is fast and can be applied to any type of comparison data. However, it does not use the individual sample data, nor does it consider the number of samples in each comparison. The second method uses per sample gene expression data. The user inputs a list of factors that indicate comparisons across or within studies and then gene level significant changes are recomputed by extracting expression data from all samples for each comparison, and then applying the RankProd (https://bioconductor.org/packages/release/bioc/html/RankProd.html)^6^ and/or metaDE (https://github.com/metaOmics/MetaDE)^7^ packages to perform meta-analysis. Limma is also applied to get statistics for each individual comparison. This analysis takes much longer (10 minutes to an hour for a typical analysis, even longer if number of samples are very large), and it has more strict sample requirements (no samples can occur in two different comparisons). Statistically, the second method is more robust but both methods should detect consistently changed genes.

#### Meta-analysis statistics

For comparison data the combined p-value is computed using Fisher’s method, that is −2*(sum of ln(p-value)) is compared against a Chi-squared distribution with N degrees of freedom, where N is the number of p-values being combined. This is carried out for every gene and yields a combined p-value that is reported. Another simpler approach is to report the maximum p-value for the gene across all the comparisons – a much more stringent measure of overall significance. This approach to combining p-values is implemented in the MetaDE R package 7. Note MetaDE will not produce results if > 30% of the comparisons for a gene have missing values for the p-value. In addition, with this approach, the p-value combination does *not* account for the direction of the fold change (up or down) so the up regulated & down regulated percentage summaries need to be referred to for interpretation.

The RankProd^6^ method converts log2 fold changes across all genes in a comparison to ranks and then computes a meta-statistic per gene which is the geometric mean of the ranks across comparisons. It is a non-parametric approach, and computes statistical significance based on permutations. Permutations also account for the multiple testing aspect of looking for significance within the set of all genes in the transcriptome.

## RESULTS AND DISCUSSION

Currently, OmicsView is published with the entire database from the Genotype-Tissue Expression project (GTEx)^8^ and a disease database containing samples from highly studied disease areas, both processed with the same computational pipeline and extracted from QIAGEN DiseaseLand. The web-based portal provides an easy-to-use framework for wet-lab biologists to interactively explore, compare and analyze data and produce publication-quality figures in SVG format, Figure 1. In addition, the user has the option of uploading their own datasets (see Supplementary Material for instructions), and by following the installation guide they can create their own private instance of the OmicsView application to store proprietary data.

**Figure 1.**
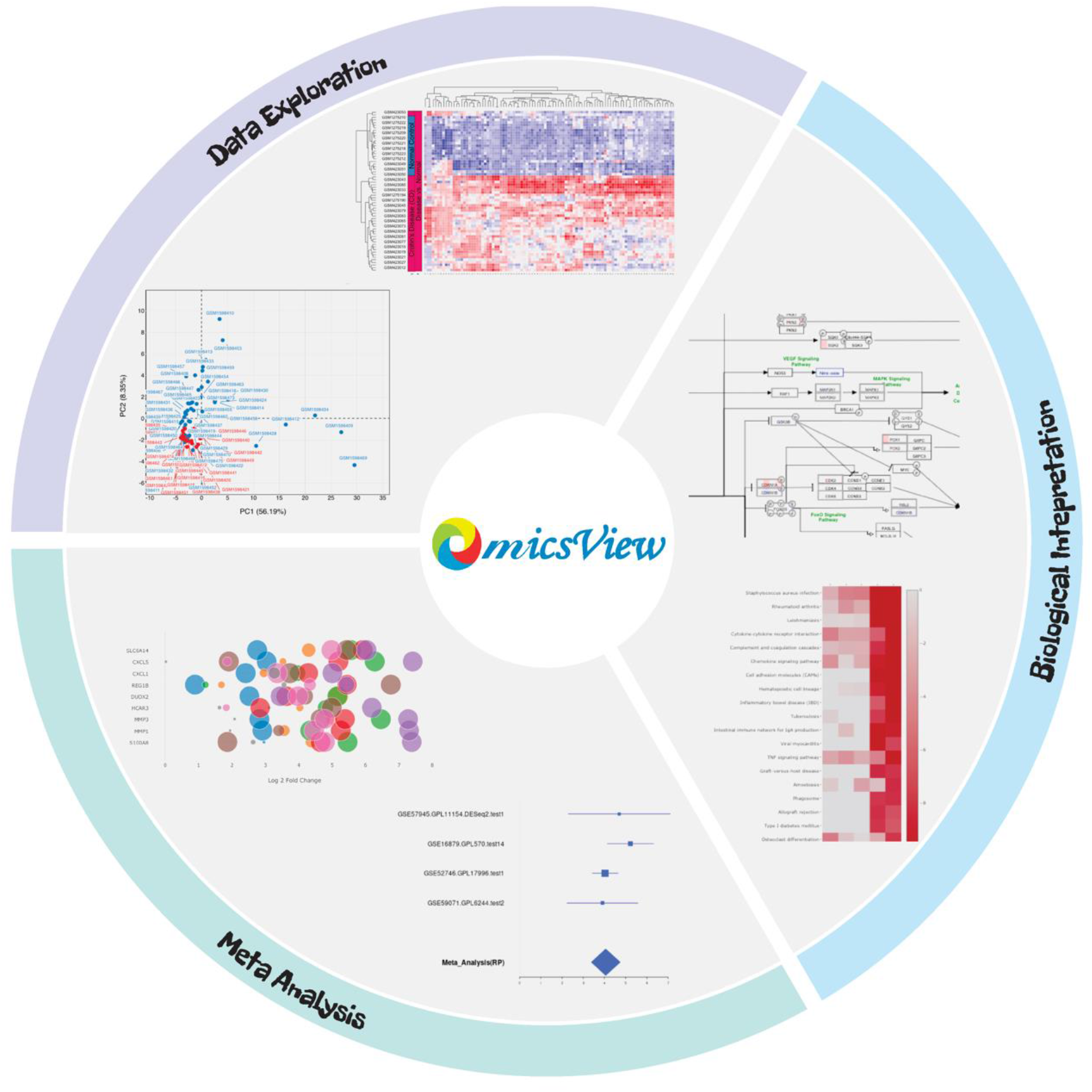
Schematic view of main analytical functionalities in OmicsView on Data Exploration by PCA, heatmap analysis, Biological Interpretation by pathway mapping and comparison, and Meta-Analysis by bubble and forest plots.

Users can easily search genes (using gene symbols, descriptions, alias or database IDs) and samples for detailed annotations in the database. By providing a list of gene names, users can compare expression of multiple genes between multiple samples within and across tissue types, disease states and disease categories. Differential gene expression between relevant groups is pre-computed along with pathway enrichment results. Gene, sample and pathway level plots are readily visualized in interactive JavaScript based views including PCA, Volcano, Multiple Gene Boxplot, Pathway Heatmap and Pathway Overlay and many more options (see Supplementary Material for user guide with all functionality).

In addition to gene expression and differential gene expression data, OmicsView’s powerful pathway analysis module pre-computes and stores pathway enrichment results for popular public databases from the Gene Ontology Consortium^9^, WikiPathways^10^, Gene Set Enrichment Analysis (GSEA - MSigDB)^11^, Kyoto Encyclopedia of Genes and Genomes (KEGG)^12^, and Reactome^13^.

### Exploration of Crohn’s disease pathology and identification of TNFalpha signaling pathway dysregulation

#### Crohn’s disease datasets

To demonstrate the power of OmicsView in discovering underlying disease pathology with minimal effort, we mine Crohn’s disease datasets that are available from QIAGEN’s DiseaseLand. The dashboard interface allows you to quickly see the distribution of disease types and determine that the current database has a good representation of Crohn’s disease datasets where disease versus normal comparisons have been carried out and are available, see Figure 2. We then select out the comparisons directly from the dashboard by selecting relevant categories. We retrieve 8 comparisons in ileum or colon tissue across several GEO studies which enables a robust meta-analysis. This can be saved to a comparison list for easy access.

**Figure 2.**
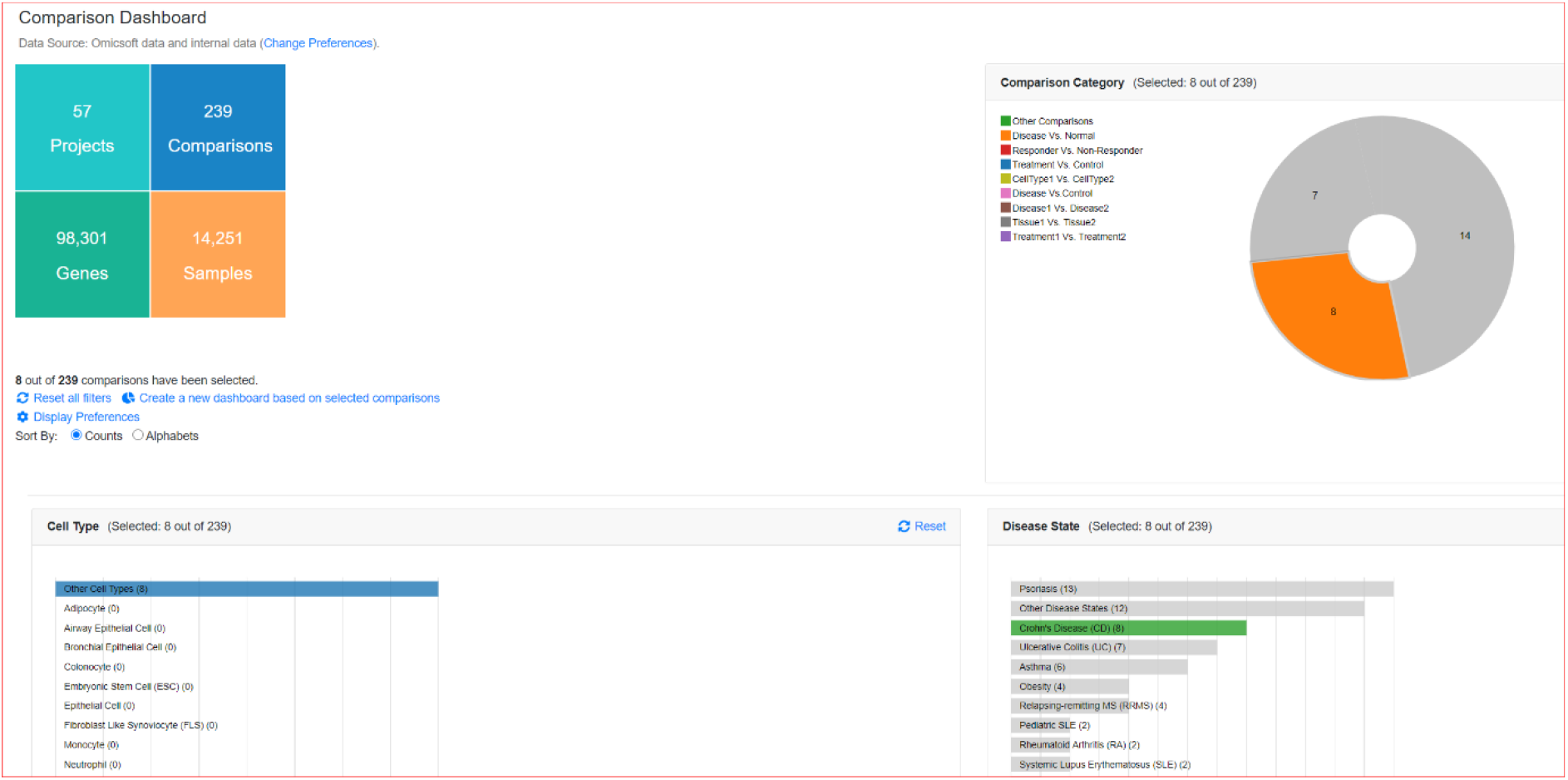
Part of “Comparison Dashboard” view in OmicsView showing breakdown of comparisons for Crohn’s disease (highlighted interactively in green barplot), 8 disease versus normal expression comparisons are available and selected (orange selection in pie chart, top right).

We next pass that through a pathway meta-analysis by using the “Pathway Heatmap” tool against all PAGE genesets to get a sense of what the general disease pathology is in Crohn’s disease. We find very strong upregulation in a number of inflammatory genesets – many of these are oncology related or computationally predicted but one canonical pathway that appears in the top 5 is TNFalpha signalling which is strongly enriched (z-score > 10 in all comparisons tested), Figure 3A. To further probe this, by clicking directly on the enrichment heatmap, we can overlay the gene-expression for each comparison onto the TNF signalling pathway from KEGG where we can identify genes that are consistently upregulated in comparisons, including TNF, CEBP and many downstream cytokines and chemokines, Figure 3B. We believe this is a novel approach to pathway enrichment visualization as it points to key genes in the pathway that are consistently dysregulated in a meta-analysis, and the overlayed network structure can suggest potential targets for modulation.

**Figure 3.**
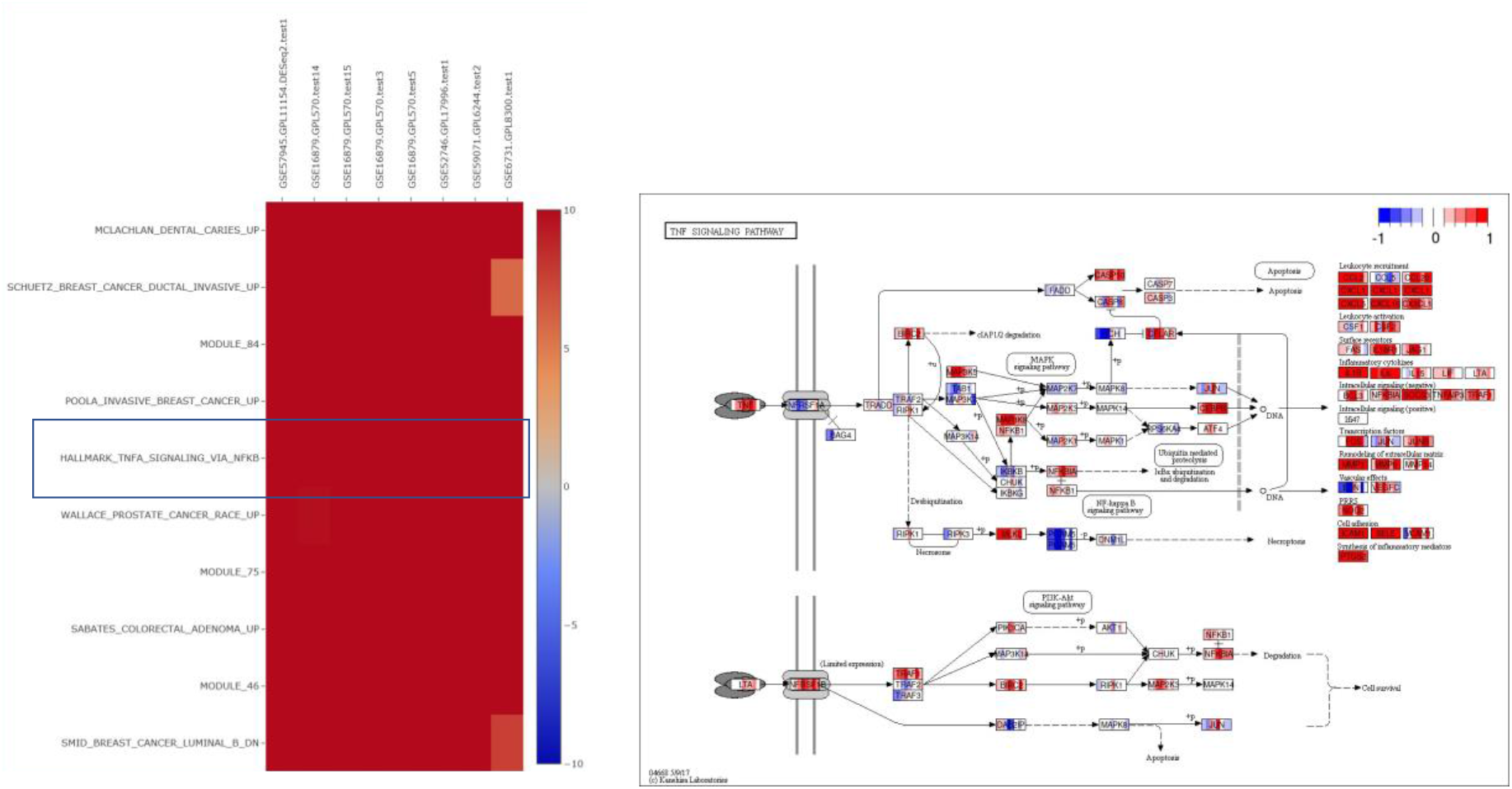
(A) PAGE results: TNFalpha signalling is a top enriched pathway across 8 Crohn’s disease dataset comparisons of disease versus normal. (B) KEGG pathway highlight of the upregulated TNF pathway showing central role for TNF and downstream consequences on chemokines and cytokines.

Then, by browsing and searching comparisons in the “Review Comparisons” interface we find that there is a GEO study, GSE52746, that includes colon gene expression after anti-TNF treatment. This comparison can be added to the disease versus control list, and on a selected set of immunologic pathways (including TNFalpha signalling) we see a striking reversal of the disease signature when comparing treatment at 12 weeks versus baseline groups, Figure 4A. A sample level representation of the full gene expression profiles can be generated from the PCA tool for this study, Figure 4B, which shows very clearly disease samples returning to normal in gene expression space, and interestingly two annotated non-responders in the study are outliers at 12 weeks.

**Figure 4.**
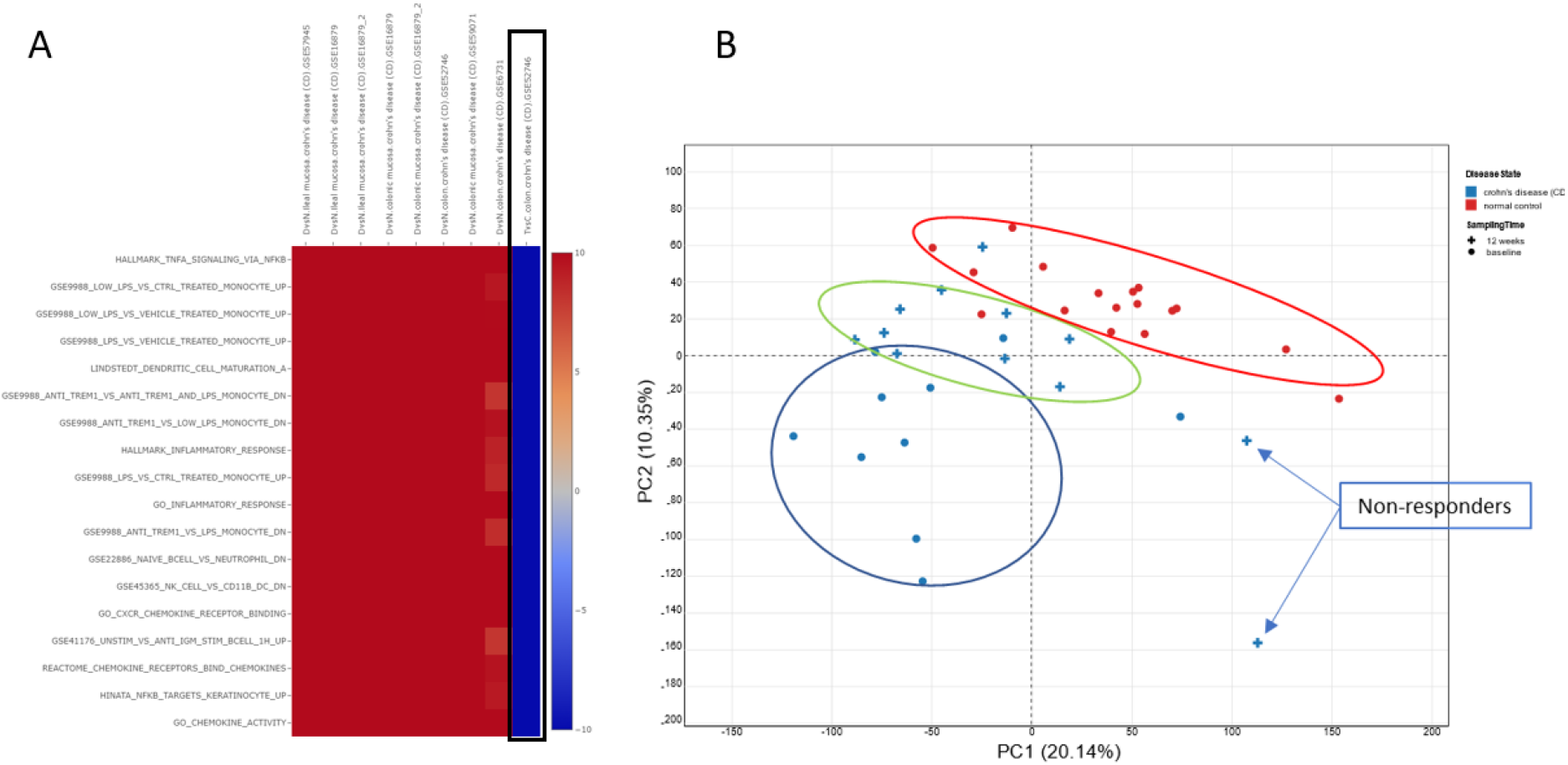
(A) GSE52746 study: Pathway scores for an anti-TNF expression profile compared to baseline untreated. On a selected set of immunologic pathways, including TNFalpha signalling (top row in the heatmap) we see a very strong reversal of the disease signature (red – upregulated) compared to TNF treated (blue – downregulated). (B) The PCA tool in OmicsView allows us to examine GSE52746 samples individually where we can see baseline disease (blue circles), normal control (red circles) and the 12 week treated samples which show an intermediate profile and are returning to control. Interestingly the two outlier treated samples are non-responders.

To query the magnitude of fold changes of the anti-TNF response, we can apply annotated volcano plots, Figure 5A where we have highlighted the genes of interest from the GSE52746 study showing the relative strength of the effect compared to the full transcriptome., Alternatively we can readily generate multi-gene boxplots where we can group and colour by relevant patient groups, Figure 5B.

**Figure 5.**
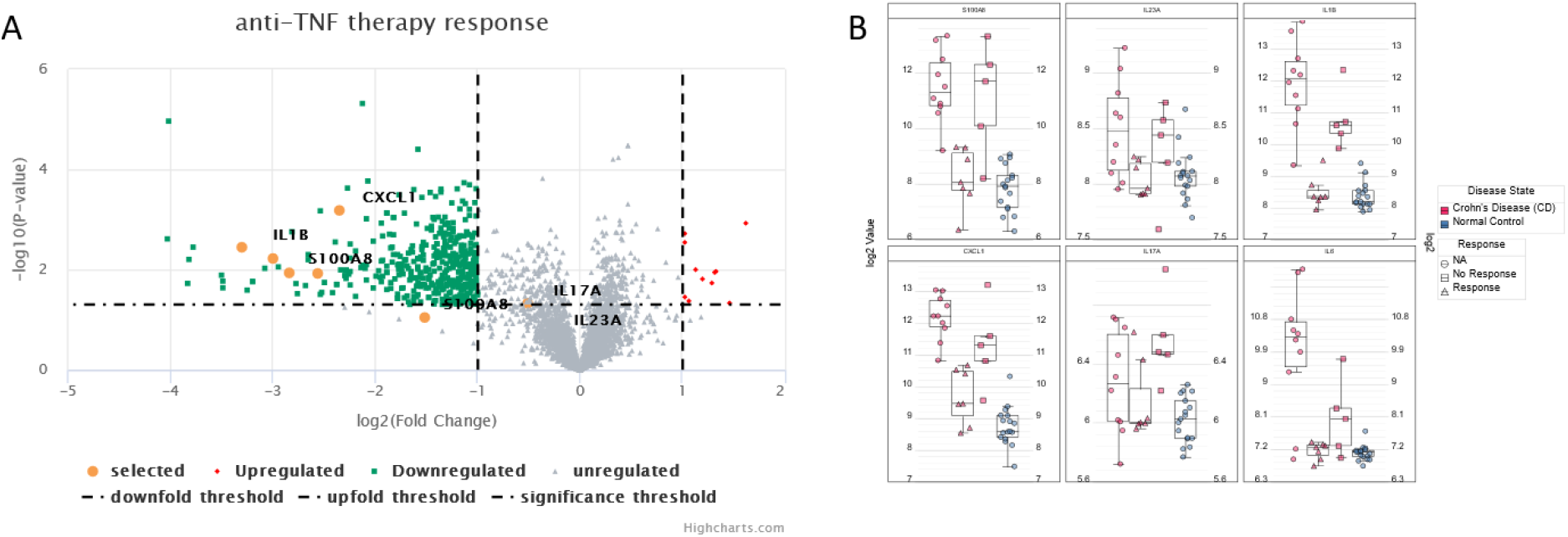
Gene level drill down into selected genes for inflammatory mediators in Crohn’s disease from GSE52746. A select set of genes from the original publication are viewed in a volcano plot for anti-TNFa response (A) and as boxplots partitioned by gene and grouped by response category (NA response means baseline samples, prior to treatment). The responders (red triangles) clearly show much more pronounced cytokine suppression towards normal levels compared to non-responders (red squares).

We may also want to look for novel biomarkers of disease independent of prior biological knowledge or pathway enrichment, so we can ask “What is the most significantly dysregulated genes across the whole transcriptome in Crohn’s disease?”. The Meta Analysis (Comparisons) tools will apply several statistics to rank genes (see Methods) by level of dysregulation across studies. To carry out the analysis, we need merely re-load the previously saved comparison list, and in Fig 6A and 6B we see the results of such an analysis on Crohn’s disease comparisons. Most of these genes can be related back to IBD, colitis or colonic epithelial cell homeostasis^15–17^. MLKL itself is in the TNF signaling pathway as can be seen in Figure 3B, and has been noted as a biomarker for intestinal inflammation in IBD^16^.

**Figure 6.**
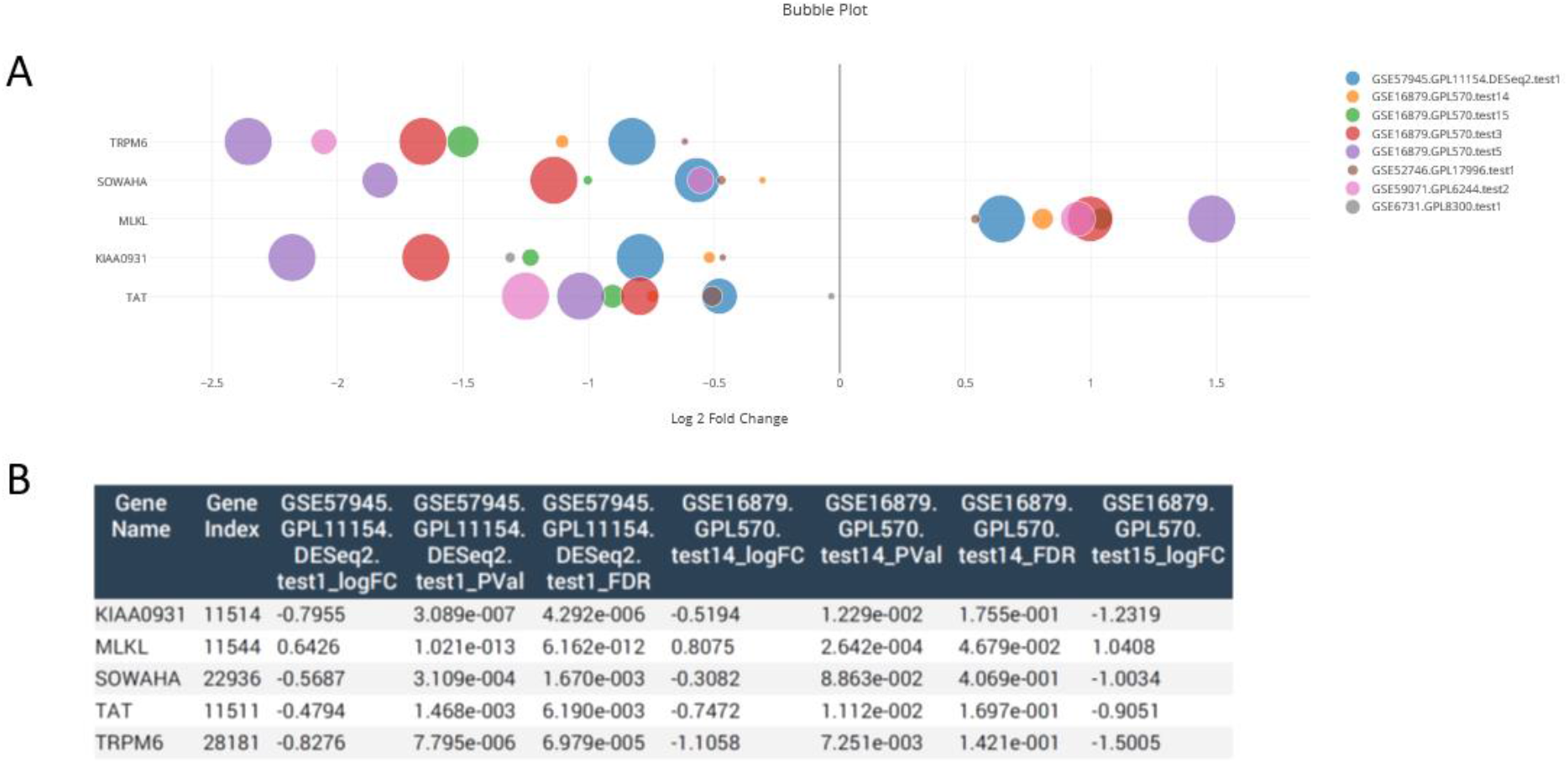
Unbiased (independent of pathway knowledge) meta-analysis of most dysregulated genes in Crohn’s disease using the Meta-analysis (Comparisons) feature of OmicsView. Panel (A) provides a bubble plot view where the consistency of changes in disease versus control comparisons across studies is apparent. Panel (B) is part of a numeric table exported showing the individual comparison data. Four out of five of these genes have literature evidence for involvement in inflammatory bowel disease, colitis and epithelial cell homeostasis.

The power of OmicsView is in these meta-analyses across studies and diseases. Another cross-disease use case is a new indication search for a given target gene. For example, motivated by the Crohn’s disease analysis, we may want to ask “In what other indications may TNF be a good target?”. This is readily answered by creating a Bubble Plot for TNF where its fold changes across many diseases and comparisons can be shown, and the top 10 can be highlighted, Figure 7. In this analysis we see, along with the consistent TNF upregulation in Crohn’s disease versus normal, there also appears to be significant TNF upregulation in psoriasis and ulcerative colitis. These validate the approach, as anti-TNF treatments (e.g. infliximab, adulumamab) are approved for both of these conditions^18,19^.

**Figure 7.**
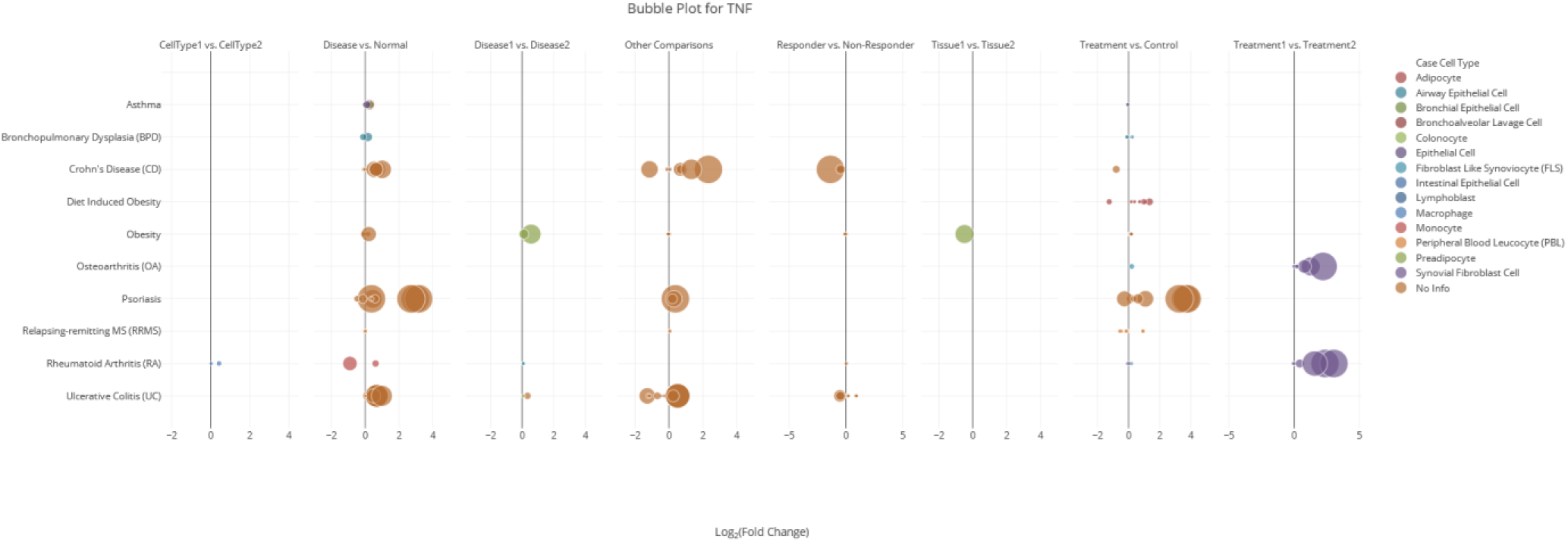
Looking at TNF fold changes across disease types in OmicsView to identify other potential indications for anti-TNF treatment. The bubble plot shows the top ten diseases (y-axis) where TNF is significantly dysregulated. The x axis is log2 fold change, and the bubble size is proportional to significance. Each panel is a different comparison type, the most relevant one for new disease discovery is “Disease vs. Normal”. We see both psoriasis and ulcerative colitis are both consistently upregulated in TNF. Note that by hovering over the “Treament1 vs. Treatment2” bubbles for osteoarthritis and rheumatoid arthritis we see that in fact this is an *in vitro* experiment with administration of TNF (versus vehicle), hence the high observed TNF levels.

## Discussion

The combined user-friendly experience, accessibility and functionality of the OmicsView pathway analysis module improves upon popular web portals like GSEA and DAVID^14^ in terms of comprehensiveness and interactivity.

Other omics results can also be overlaid along with transcriptomics for integrated data analysis with appropriate name mapping in the input dataset, and by means of the metanalysis features. By combining the latest advancement of visualization, harmonized state-of-the-art bioinformatics analysis and a large publicly available expression dataset, OmicsView is an easy-to-use powerful platform that can benefit all scientists, especially ones with limited programming skills in the biomedical research field. The entire system is released as open source for wide adoption and further enhancement. The Supplementary Material outlines the detailed steps for interacting with the system and generating plots and analyses of interest.

## DATA AVAILABILITY

OmicsView source code and installation guide is freely available at https://github.com/interactivereport/OmicsView

A publicly hosted instance of OmicsView, preloaded with GTEx and selected DiseaseLand studies is available at http://omicsview.org

## SUPPLEMENTARY DATA

Supplementary Data are available at NAR online.

## ACKNOWLEDGEMENT

We thank QIAGEN for allowing use of partial data from their curated DiseaseLand data sets (https://www.qiagenbioinformatics.com/diseaseland/), and the institutes and authors who made their research data available.

## FUNDING

No public funding source was utilized for this work.

## CONFLICT OF INTEREST

The authors have no conflict of interest to declare.

## Notes

http://omicsview.org

### Competing Interest Statement

The authors have declared no competing interest.

