## Supplementary Information for "OmicsView: omics data analysis through interactive visual analytics"

### OmicsView Supplementary

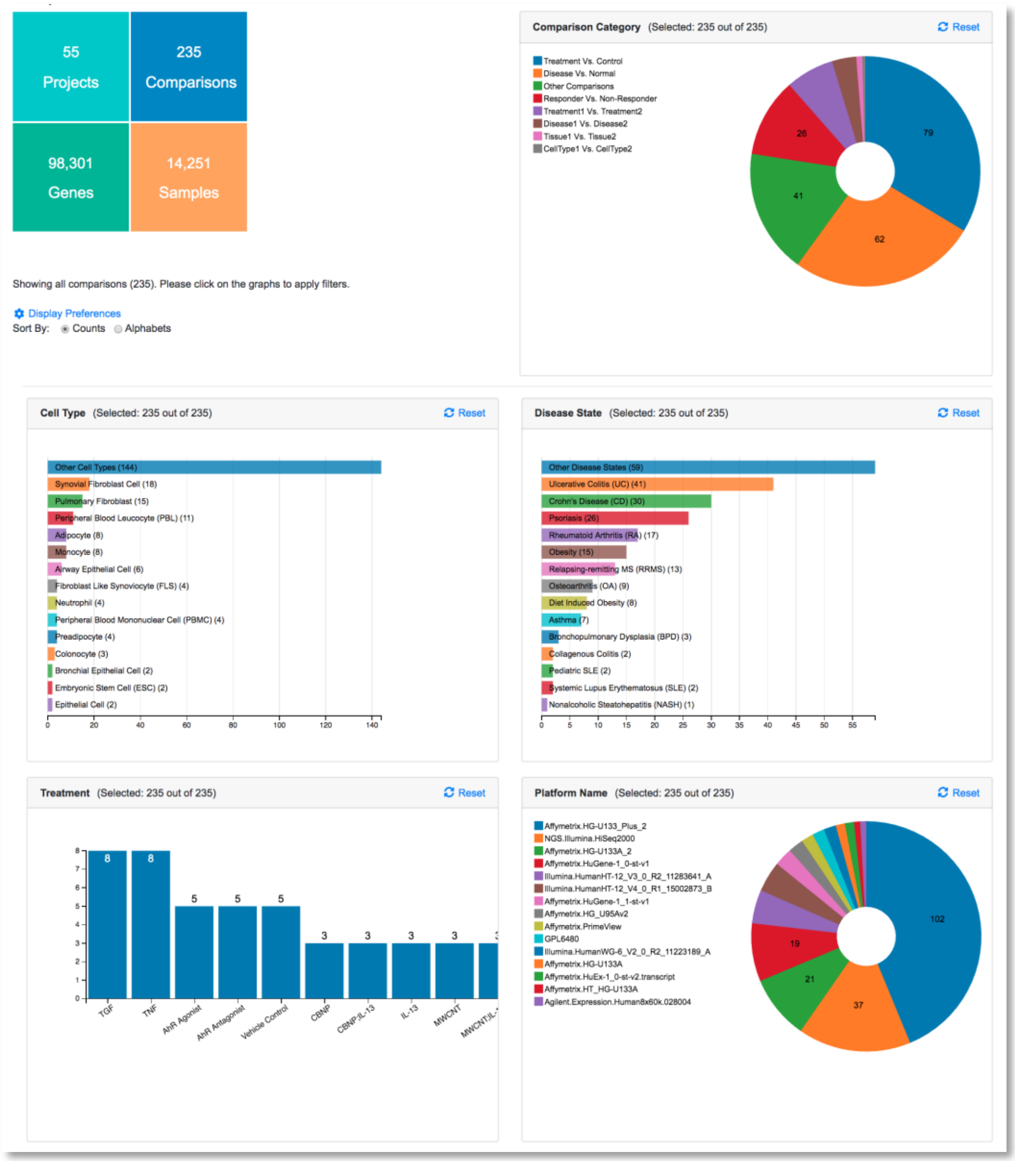

**[http://omicsview.org/](http://omicsview.org/)**

From the login page, users can use their email to register an account (recommended), as this enables users to save results and upload their own data. Otherwise, a guest account can be used to view public data.

### 1. Overview of Data in OmicsView

The homepage of OmicsView gives an overview of the data curated. The Pie-charts and Bar-graphs are reactive and change depending on the selection. Users can access the different data tables from the left menu under search, or from the shortcuts at the top menu bar. Left menu can be hidden to provide more space for tables and graphs.

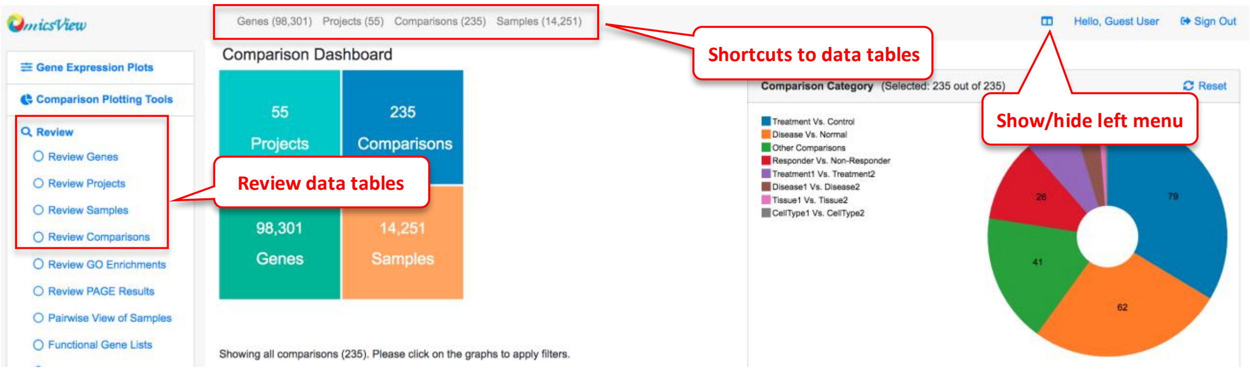

#### 1.1 Projects

Individual research projects are typically associated with a published study and has NCBI GEO accession number. Users can search projects from the home page. In the search result page, click each Project ID to view the full description of a project.

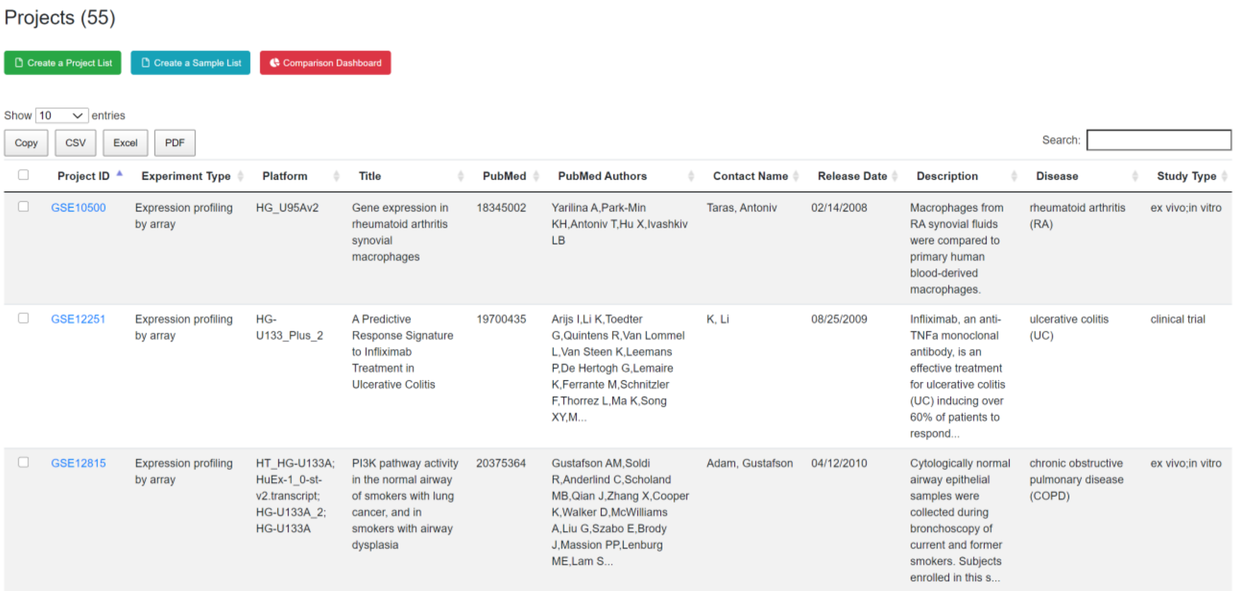

Projects (55)

Create a Project List | Create a Sample List | Comparison Dashboard

Show 10 entries

Copy | CSV | Excel | PDF

Search:

| Project ID | Experiment Type | Platform | Title | PubMed | PubMed Authors | Contact Name | Release Date | Description | Disease | Study Type |
|---|---|---|---|---|---|---|---|---|---|---|
| GSE10500 | Expression profiling by array | HG_U95Av2 | Gene expression in rheumatoid arthritis synovial macrophages | 18345002 | Yarilina A,Park-Min KH,Antoniv T,Hu X,Ivashkiv LB | Taras, Antoniv | 02/14/2008 | Macrophages from RA synovial fluids were compared to primary human blood-derived macrophages. | rheumatoid arthritis (RA) | ex vivo;in vitro |
| GSE12251 | Expression profiling by array | HG-U133_Plus_2 | A Predictive Response Signature to Infliximab Treatment in Ulcerative Colitis | 19700435 | Arijs I,Li K,Toedter G,Quintens R,Van Lommel L,Van Steen K,Leemans P,De Hertogh G,Lemaire K,Ferrante M,Schnitzler F,Thorrez L,Ma K,Song XY,M... | K, Li | 08/25/2009 | Infliximab, an anti-TNFa monoclonal antibody, is an effective treatment for ulcerative colitis (UC) inducing over 60% of patients to respond... | ulcerative colitis (UC) | clinical trial |
| GSE12815 | Expression profiling by array | HT_HG-U133A; HuEx-1_0-st-v2.transcript; HG-U133A_2; HG-U133A | PI3K pathway activity in the normal airway of smokers with lung cancer, and in smokers with airway dysplasia | 20375364 | Gustafson AM,Soldi R,Anderlind C,Scholand MB,Qian J,Zhang X,Cooper K,Walker D,McWilliams A,Liu G,Szabo E,Brody J,Massion PP,Lenburg ME,Lam S... | Adam, Gustafson | 04/12/2010 | Cytologically normal airway epithelial samples were collected during bronchoscopy of current and former smokers. Subjects enrolled in this s... | chronic obstructive pulmonary disease (COPD) | ex vivo;in vitro |

#### 1.2 Samples

A project may include many samples. Each sample has its own description (full details available by clicking each Sample ID). Each sample has a gene expression profile.

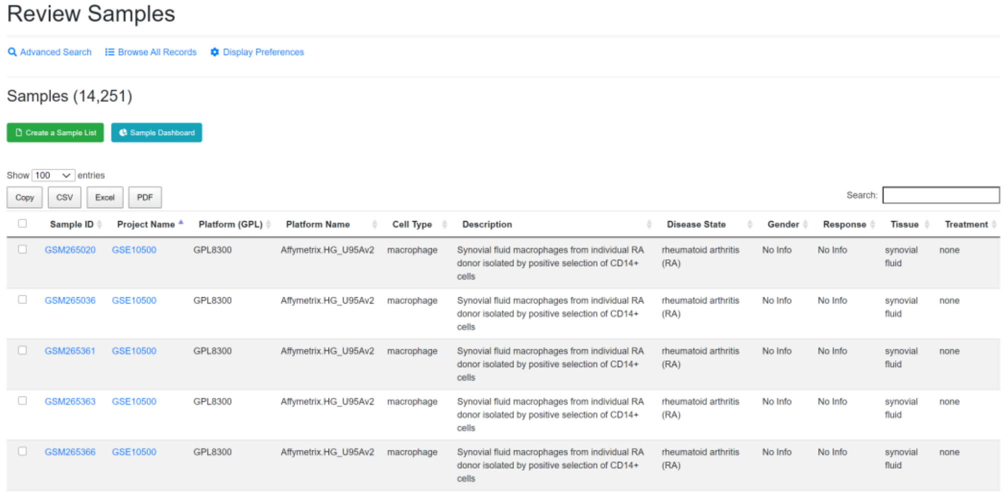

Review Samples

Advanced Search | Browse All Records | Display Preferences

Samples (14,251)

Create a Sample List | Sample Dashboard

Show 100 entries

Copy | CSV | Excel | PDF

Search:

| | Sample ID | Project Name | Platform (GPL) | Platform Name | Cell Type | Description | Disease State | Gender | Response | Tissue | Treatment |
|---|---|---|---|---|---|---|---|---|---|---|---|
| | GSM265020 | GSE10500 | GPL8300 | Affymetrix.HG_U95Av2 | macrophage | Synovial fluid macrophages from individual RA donor isolated by positive selection of CD14+ cells | rheumatoid arthritis (RA) | No Info | No Info | synovial fluid | none |
| | GSM265036 | GSE10500 | GPL8300 | Affymetrix.HG_U95Av2 | macrophage | Synovial fluid macrophages from individual RA donor isolated by positive selection of CD14+ cells | rheumatoid arthritis (RA) | No Info | No Info | synovial fluid | none |
| | GSM265361 | GSE10500 | GPL8300 | Affymetrix.HG_U95Av2 | macrophage | Synovial fluid macrophages from individual RA donor isolated by positive selection of CD14+ cells | rheumatoid arthritis (RA) | No Info | No Info | synovial fluid | none |
| | GSM265363 | GSE10500 | GPL8300 | Affymetrix.HG_U95Av2 | macrophage | Synovial fluid macrophages from individual RA donor isolated by positive selection of CD14+ cells | rheumatoid arthritis (RA) | No Info | No Info | synovial fluid | none |
| | GSM265366 | GSE10500 | GPL8300 | Affymetrix.HG_U95Av2 | macrophage | Synovial fluid macrophages from individual RA donor isolated by positive selection of CD14+ cells | rheumatoid arthritis (RA) | No Info | No Info | synovial fluid | none |

Users can search for specific samples using the search box.

To change columns displayed in the table, go to the Display Preferences.

Users can create lists by selecting specific samples using the “Create a Sample List” button. Users can then specifically download this list as a CSV.Excel/PDF. Samples from your collection can be loaded to other tools like heatmap.

#### 1.3 Comparisons

Two groups of samples are compared, and the statistic values (fold change, p-value, adjusted p-values) are computed.

There are a lot of meta data available for each comparison. See the dashboard for an overview of key categories, and the detailed description of each comparison has the full information.

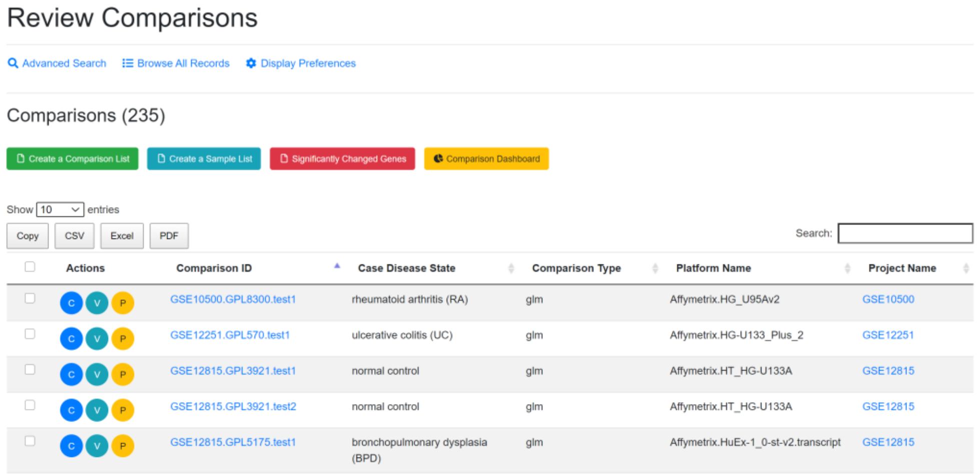

Review Comparisons

Advanced Search | Browse All Records | Display Preferences

Comparisons (235)

Create a Comparison List | Create a Sample List | Significantly Changed Genes | Comparison Dashboard

Show 10 entries

Copy | CSV | Excel | PDF

Search:

| | Actions | Comparison ID | Case Disease State | Comparison Type | Platform Name | Project Name |
|---|---|---|---|---|---|---|
| | C V P | GSE10500.GPL8300.test1 | rheumatoid arthritis (RA) | glm | Affymetrix.HG_U95Av2 | GSE10500 |
| | C V P | GSE12251.GPL570.test1 | ulcerative colitis (UC) | glm | Affymetrix.HG-U133_Plus_2 | GSE12251 |
| | C V P | GSE12815.GPL3921.test1 | normal control | glm | Affymetrix.HT_HG-U133A | GSE12815 |
| | C V P | GSE12815.GPL3921.test2 | normal control | glm | Affymetrix.HT_HG-U133A | GSE12815 |
| | C V P | GSE12815.GPL5175.test1 | bronchopulmonary dysplasia (BPD) | glm | Affymetrix.HuEx-1_0-st-v2.transcript | GSE12815 |

View full details of a comparison by clicking the (Review Comparison) button.

View volcano plot by clicking the (View Volcano Chart) button.

Users can also map the comparison data onto wikipathways by clicking the (View Pathway) button.

The selected comparisons can be saved to your collection by creating a list using the “Create a Comparison List” button for easy loading into the plotting tools.

#### 1.4 Genes

The genome-wide gene expression values were detected in each sample using RNA-Seq or microarrays. All the human genes that have expression values are listed in gene table. The gene annotation from different platforms were all mapped to NCBI gene ID (EntrezID) for consistency across platforms.

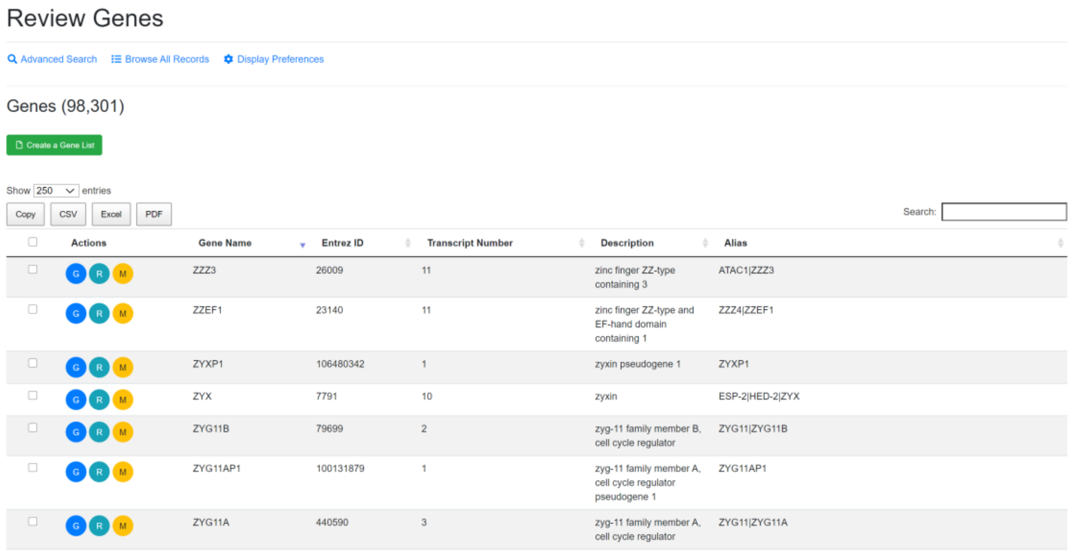
Review Genes

Advanced Search  Browse All Records  Display Preferences

Genes (98,301)

Create a Gene List

Show 250 entries

Copy CSV Excel PDF  Search:

| | Actions | Gene Name | Entrez ID | Transcript Number | Description | Alias |
|---|---|---|---|---|---|---|
| | G R M | ZZZ3 | 26009 | 11 | zinc finger ZZ-type containing 3 | ATAC1\|ZZZ3 |
| | G R M | ZZEF1 | 23140 | 11 | zinc finger ZZ-type and EF-hand domain containing 1 | ZZZ4\|ZZEF1 |
| | G R M | ZYXP1 | 106480342 | 1 | zyxin pseudogene 1 | ZYXP1 |
| | G R M | ZYX | 7791 | 10 | zyxin | ESP-2\|HED-2\|ZYX |
| | G R M | ZYG11B | 79699 | 2 | zyg-11 family member B, cell cycle regulator | ZYG11\|ZYG11B |
| | G R M | ZYG11AP1 | 100131879 | 1 | zyg-11 family member A, cell cycle regulator pseudogene 1 | ZYG11AP1 |
| | G R M | ZYG11A | 440590 | 3 | zyg-11 family member A, cell cycle regulator | ZYG11\|ZYG11A |

To find a gene, Users can specifically search in each field like gene symbol, gene description, gene alias, NCBI gene ID, Ensembl gene ID or Uniprot ID in the Advanced Search option.

For some common genes, the symbols used in publications are often not the official symbol, and users can try search alias field in Advanced Search. For example, TP53 is often referred to as P53 in publication. Users need to search P53 in alias or tumor protein p53 in description to find it if you don't know its official symbol.

The NCBI Gene search https://www.ncbi.nlm.nih.gov/gene is a good source to get official gene symbols and IDs.

Users can view full details of a gene by clicking the (Review Gene) button.

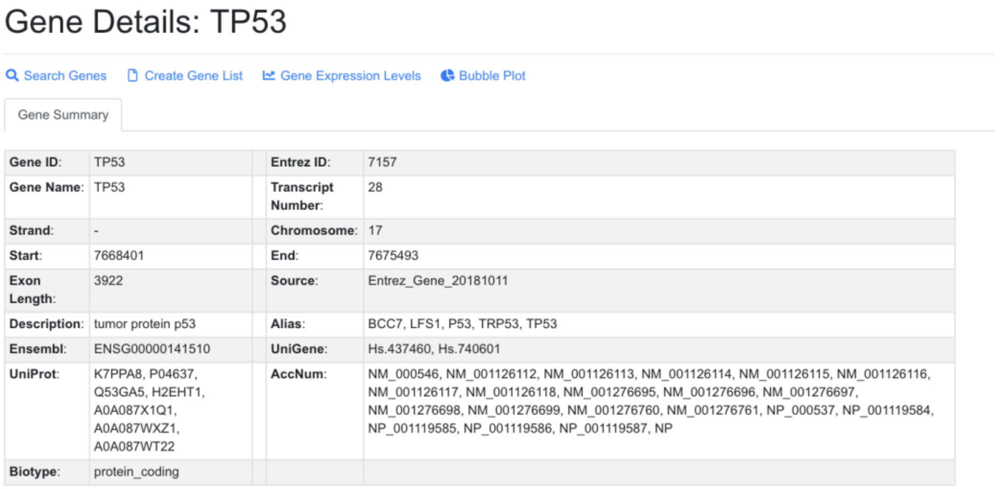

| | | | |
|---|---|---|---|
| **Gene ID**: | TP53 | **Entrez ID**: | 7157 |
| **Gene Name**: | TP53 | **Transcript Number**: | 28 |
| **Strand**: | - | **Chromosome**: | 17 |
| **Start**: | 7668401 | **End**: | 7675493 |
| **Exon Length**: | 3922 | **Source**: | Entrez_Gene_20181011 |
| **Description**: | tumor protein p53 | **Alias**: | BCC7, LFS1, P53, TRP53, TP53 |
| **Ensembl**: | ENSG00000141510 | **UniGene**: | Hs.437460, Hs.740601 |
| **UniProt**: | K7PPA8, P04637, Q53GA5, H2EHT1, A0A087X1Q1, A0A087WXZ1, A0A087WT22 | **AccNum**: | NM_000546, NM_001126112, NM_001126113, NM_001126114, NM_001126115, NM_001126116, NM_001126117, NM_001126118, NM_001276695, NM_001276696, NM_001276697, NM_001276698, NM_001276699, NM_001276760, NM_001276761, NP_000537, NP_001119584, NP_001119585, NP_001119586, NP_001119587, NP |
| **Biotype**: | protein_coding | | |

From gene details, Users can access RNA-Seq data in a box plot, or view all comparisons including this gene in a bubble plot.

#### 1.5 Saved Genes and Comparisons

Users can save selected genes or comparison for future use (e.g. multiple gene and multiple comparisons bubble plot). From gene search, check the genes you want to save, and click the yellow button "save selected genes

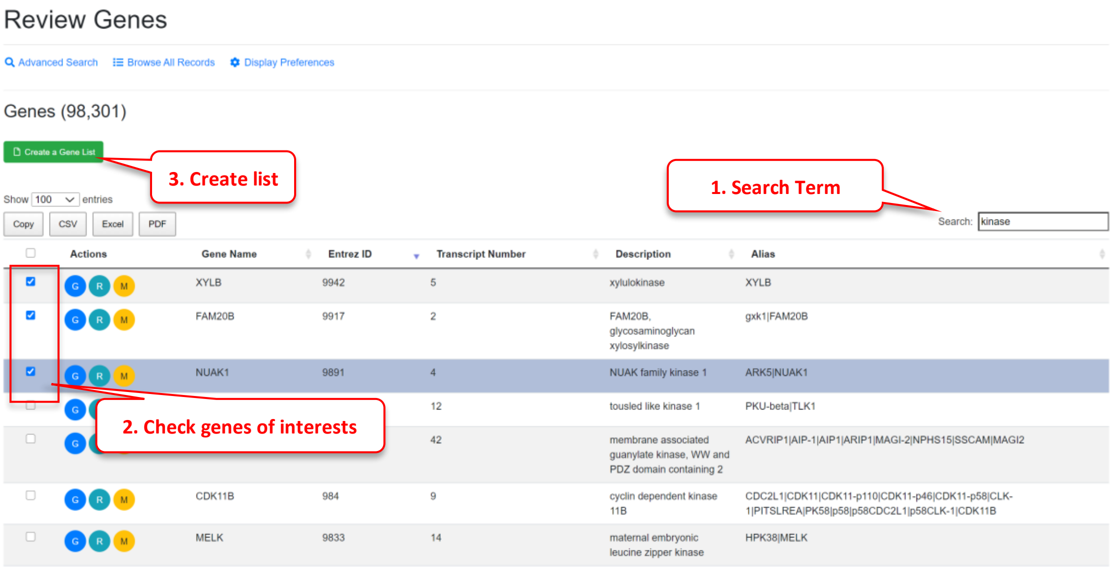

| Actions | Gene Name | Entrez ID | Transcript Number | Description | Alias |
|---|---|---|---|---|---|
| G R M | XYLB | 9942 | 5 | xylulokinase | XYLB |
| G R M | FAM20B | 9917 | 2 | FAM20B, glycosaminoglycan xylosylkinase | gxk1\|FAM20B |
| G R M | NUAK1 | 9891 | 4 | NUAK family kinase 1 | ARK5\|NUAK1 |
| G | | | 12 | tousled like kinase 1 | PKU-beta\|TLK1 |
| G | | | 42 | membrane associated guanylate kinase, WW and PDZ domain containing 2 | ACVRIP1\|AIP-1\|AIP1\|ARIP1\|MAGI-2\|NPHS15\|SSCAM\|MAGI2 |
| G R M | CDK11B | 984 | 9 | cyclin dependent kinase 11B | CDC2L1\|CDK11\|CDK11-p110\|CDK11-p46\|CDK11-p58\|CLK-1\|PITSLREA\|PK58\|p58\|p58CDC2L1\|p58CLK-1\|CDK11B |
| G R M | MELK | 9833 | 14 | maternal embryonic leucine zipper kinase | HPK38\|MELK |

Similarly, you can save comparisons.

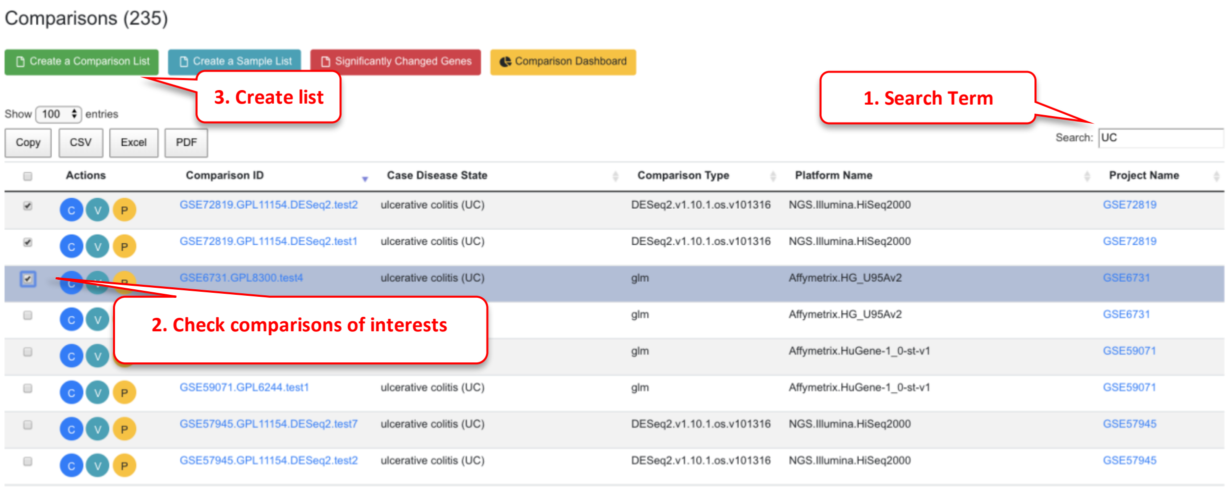

To view saved genes, click "My Results" link on the left menu and then "Gene Lists".

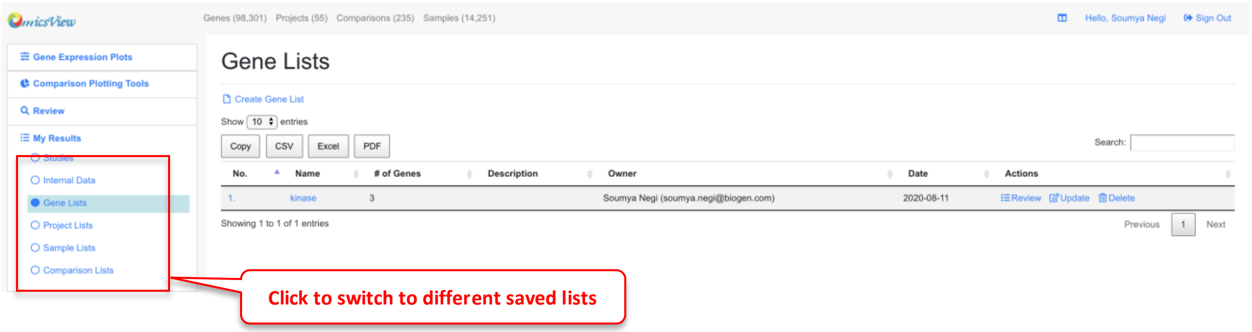

Similarly, to view the saved comparisons, select "Comparison Lists" under "My Results":

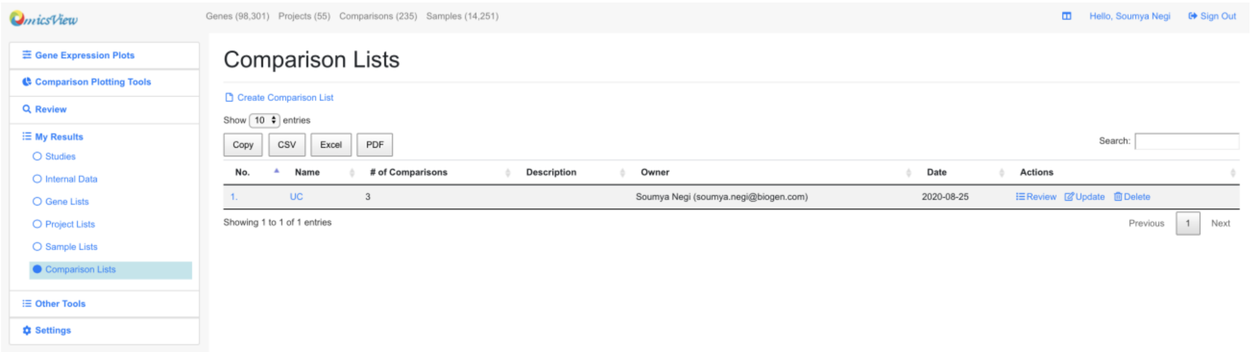

### 2 Visualize Gene Expression

For each gene, Users can view its expression levels across multiple samples. Most data in OmicsView are from microarray data, consisting of more than 50K samples, and nearly 3000 samples are from RNA-Seq.

#### 2.1 View Gene Expression from RNA-Seq

Choose the Gene Expression from RNA-Seq -> Single Gene from left menu, and enter the official symbol of gene. Alternatively, in the gene details page, click View Gene Expression link.

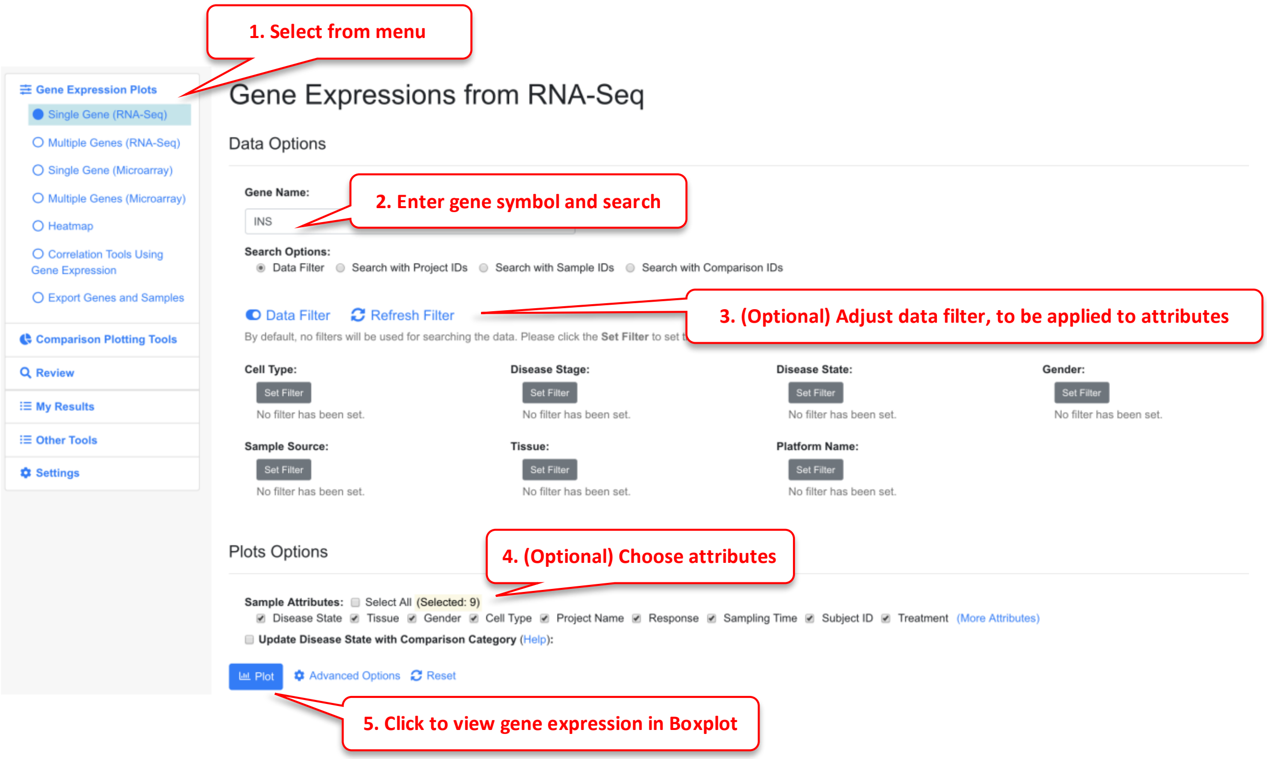

As an optional step, Users can choose what sample attributes to pass to the plot and use data filter to choose only a subset of data points.

The Data Filter can be very useful if there are too many data points, and you can focus on a few diseases or tissue types.

The screenshots below show default boxplot showing all diseases, and by after filtering for few diseases.

Plot showing all disease states (scroll to view other diseases). 5000 randomly selected data points out of 9538 are shown.

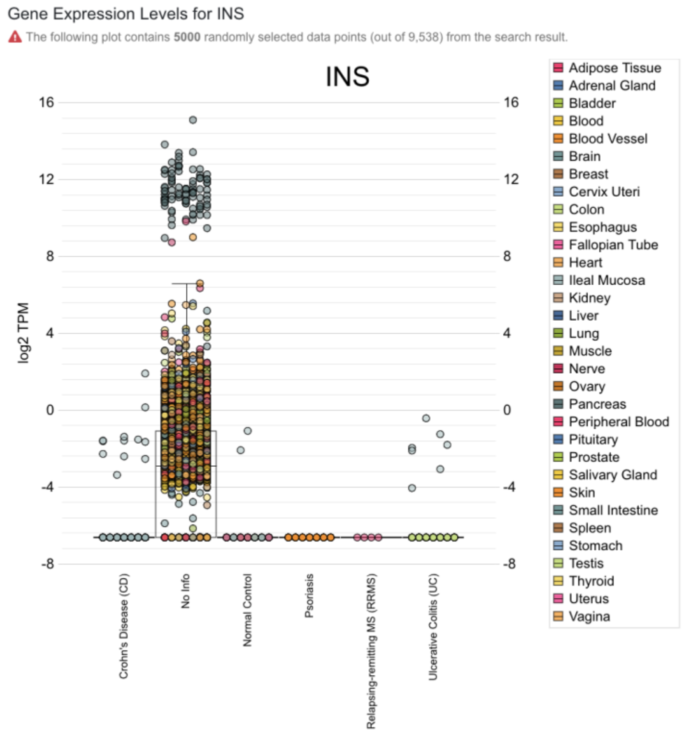

Plot after data filtering was applied to select a few diseases. Now 410 out of 9538 data points are shown. The data filter pop-up window was shown to the left of the box plot in the screen shot below.

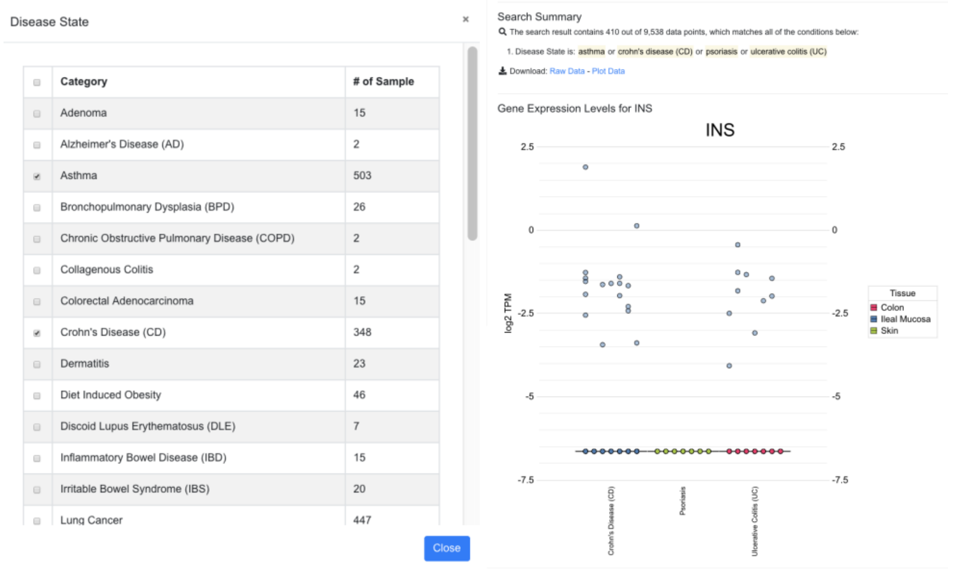

#### 2.2 Change Sample Grouping in Gene Expression Plot

The boxplot is created using CanvasXpress ( https://canvasxpress.org )plug-in, and sample grouping and coloring can be customized by the user. In the example below, we show how sample grouping can be changed.

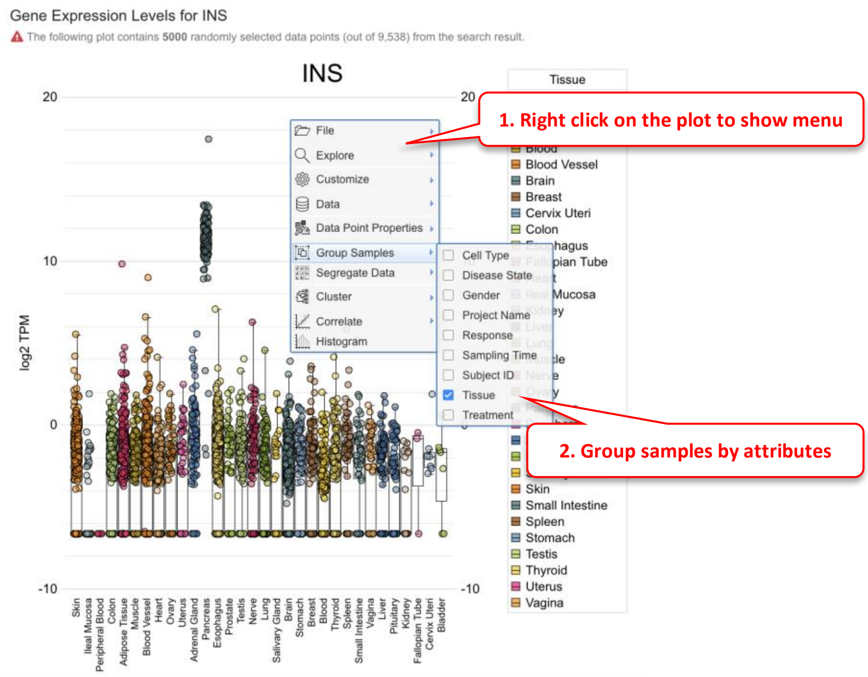

Once the sample grouping was changed to tissue, the box plot shows that insulin is only expressed in pancreatic islets. Users can also change how the data points are colored (default is by disease).

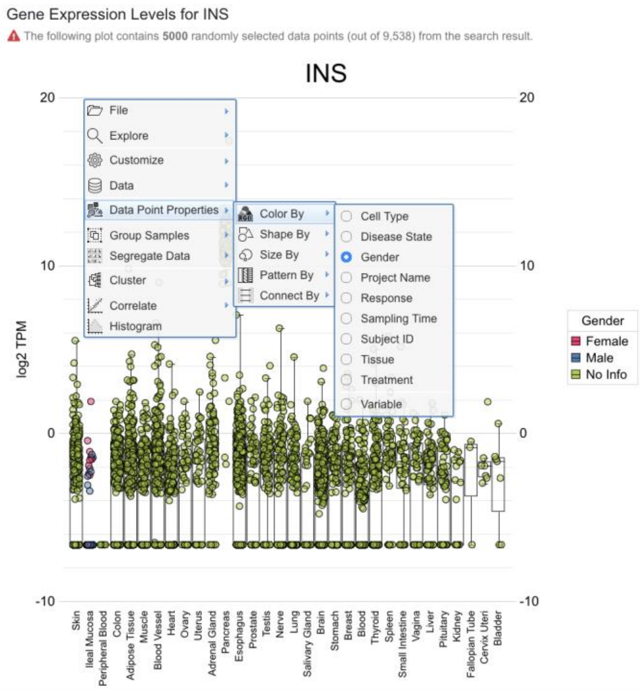

#### 2.3 View Gene Expression from Microarray Data

The way to view microarray data is very similar to RNA-Seq data. However, since the expression values from different array platforms are typically not directly comparable, the system by default will choose the platform with the largest number of data points. The user can override this filter if needed.

In addition, we recommend the user to add data filter because typically there are too many data points from microarray data. Data filters will help the user to focus on the most important tissues or diseases. If there are still more than 5000 data points after data filer, only the first 5000 are shown in the boxplot.

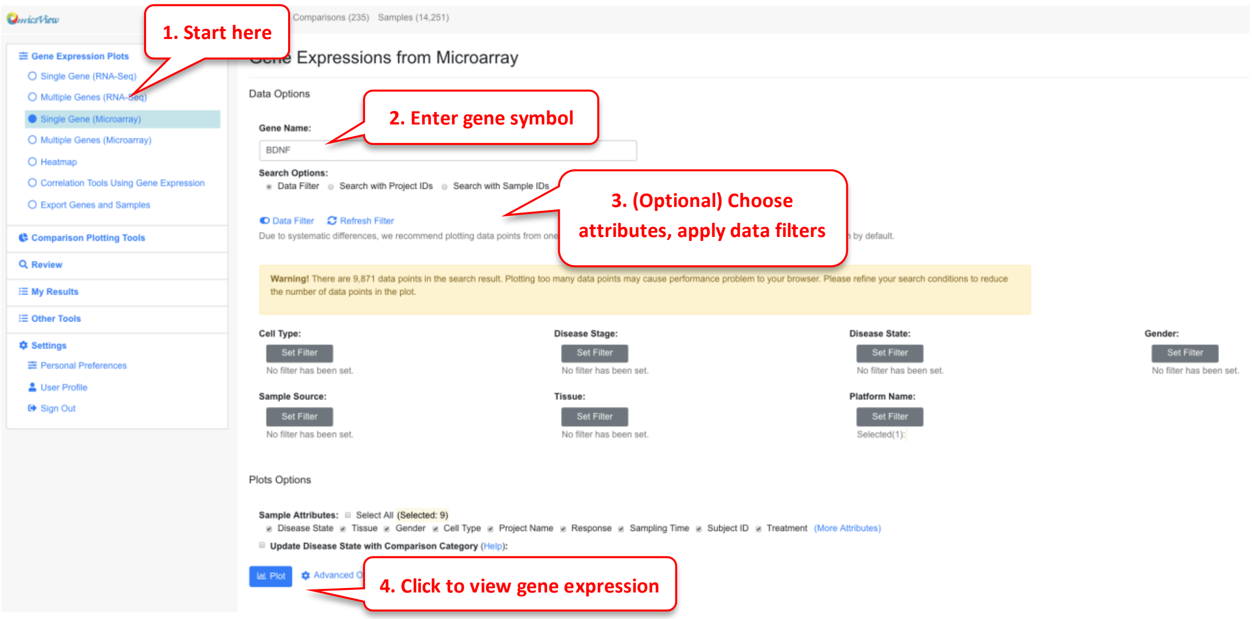

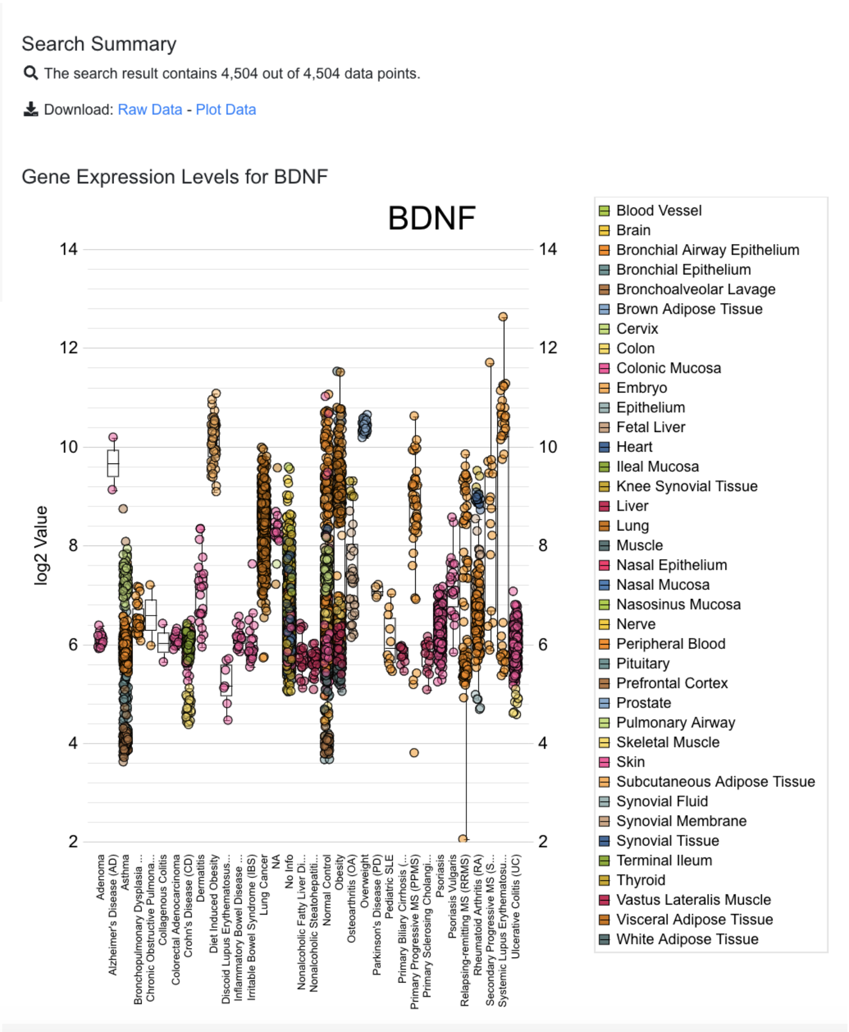

The sample grouping and coloring can be changed by the user. See the section before for details.

#### 2.4 View Gene Expression from Multiple Genes

Users can view multiple genes on the same boxplot. The interface is similar to single gene, here you need to enter a list of gene symbols. Users can also load the saved gene lists from your collection. In multiple gene plot, you want to filter the data to come up with a reasonable number of data points.

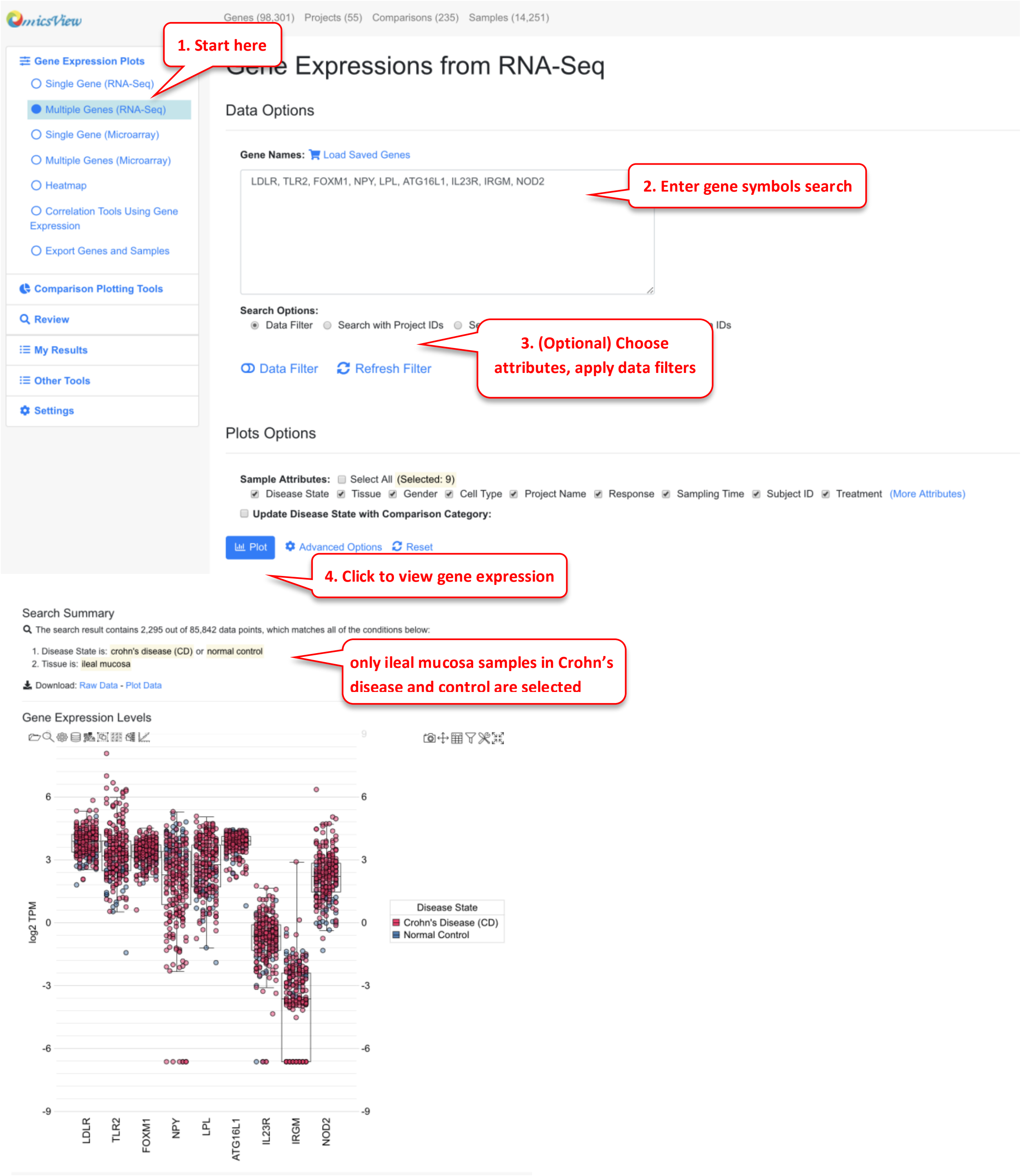

From multiple gene plot, you can use the built-in function of CanvassXpress to change grouping and coloring.

In the example plot above, we can change the setting to use group by to distinguish Crohn's disease samples and control samples.

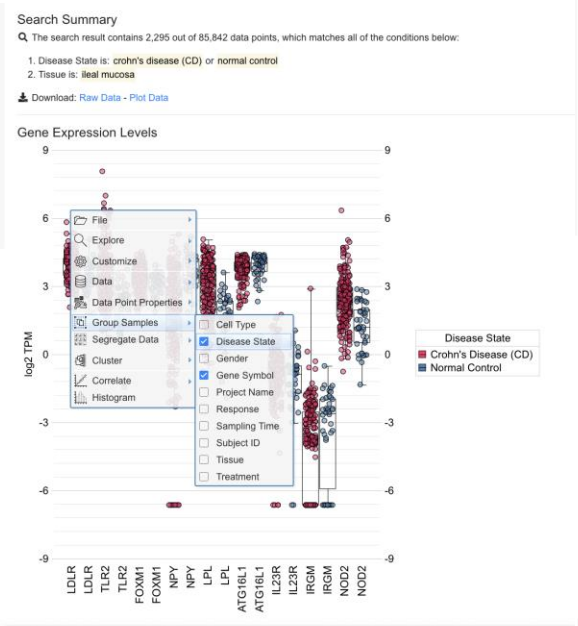

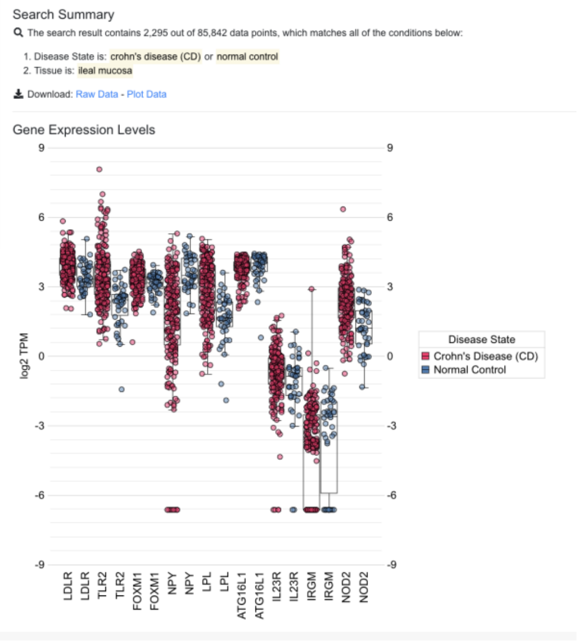

In the updated plot above, it's easy to see that many of these genes are expressed slightly higher in disease (red) vs. control (blue).

#### 2.5 View Gene Expression in Heatmap

Heatmap can be useful to visualize gene profiles from multiple samples. It can also provide information about how genes and samples cluster. This data is from PMID: 25003194

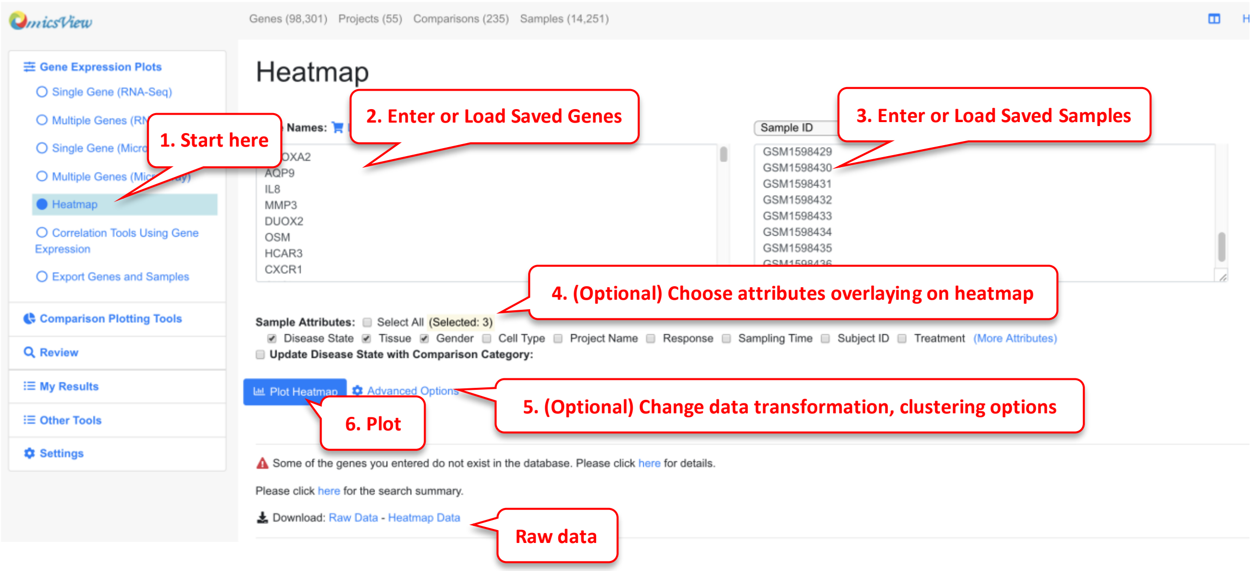

Users can enter genes and samples in the box, or load pre-saved genes and samples quickly from your collection. Be default, we will log2 transform the gene expression data, perform scaling of the data across samples for each gene, and limit the scaled value to -3 to 3 before displaying the data in heatmap. This works well in most situations. However, advanced users can change the options. For example, if you want to keep the order of samples as you entered, just uncheck “Enable Clustering Samples”.

The heatmap is rendered by CanvasXpress. Users can change the plot size if needed.

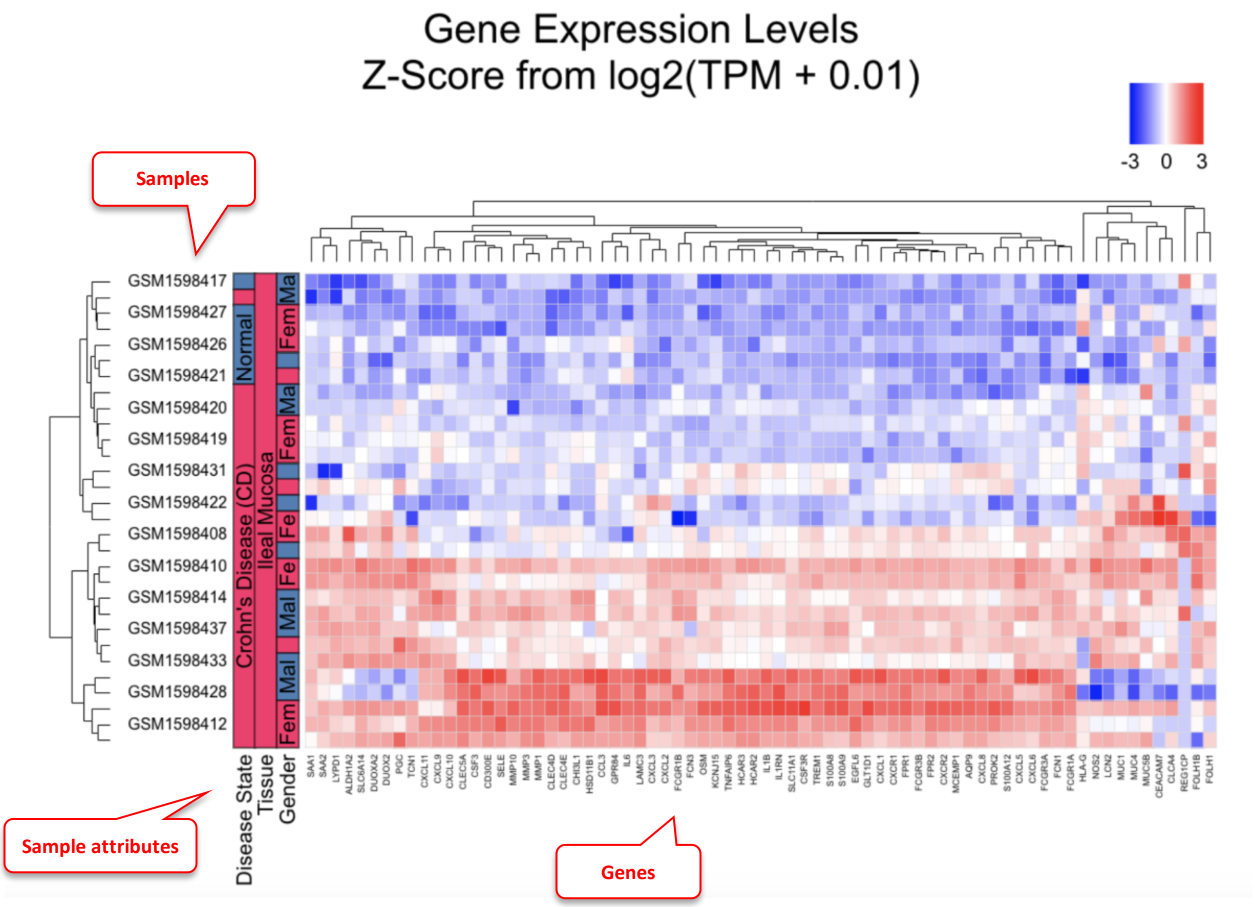

In the example heatmap, we entered a few significantly changed genes in Crohn's disease, and choose a few disease and control samples. As shown in the heatmap, most of these genes show distinct patterns between disease and control.

One of the samples GSM1598422 is labeled as disease, but its gene expression signature match those with normal control very well. Therefore from heatmap clustering, we can decide that this sample is likely an outlier, the sample may be mislabeled, or this patient had very different gene expression patterns form all other patients.

#### 2.6 Export Genes and Samples

This is a useful feature to download raw (counts) and processed (TPM values) expression matrices from the datasets. This gives the user the opportunity to do any downstream analysis on the raw reads independently.

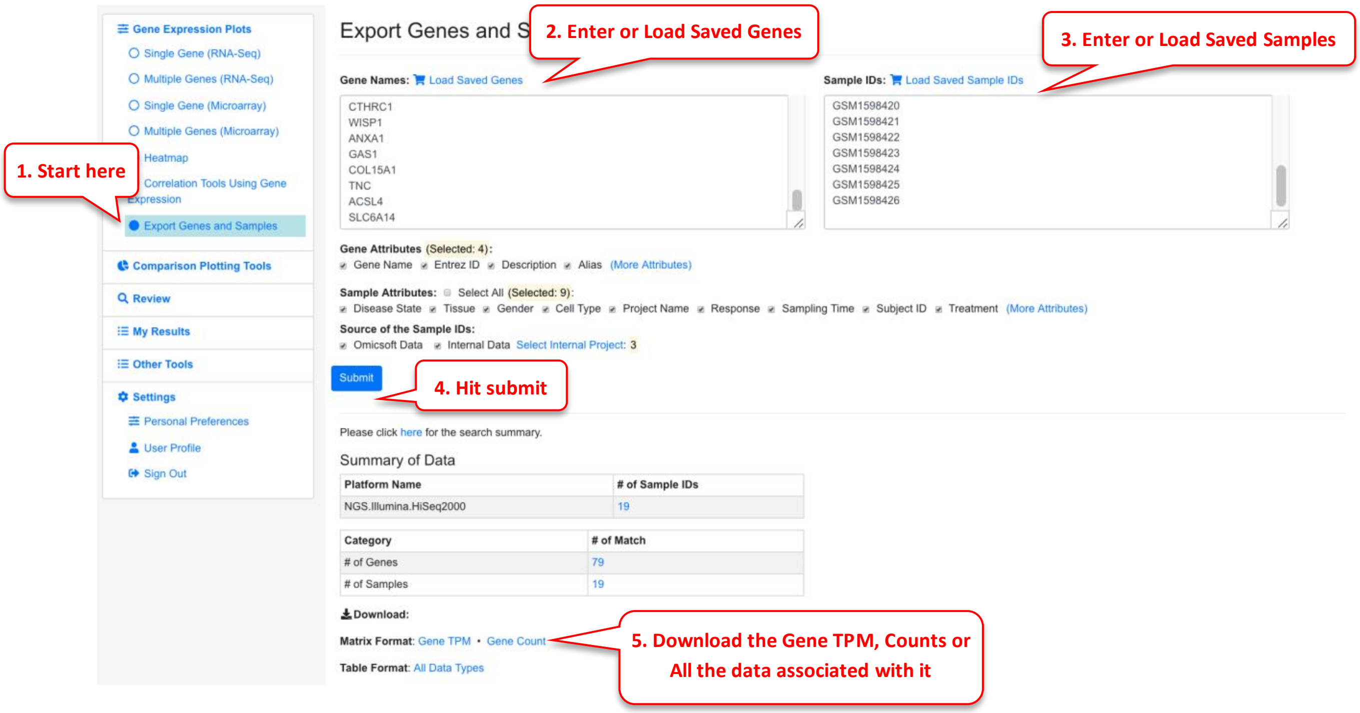

For this functionality, the samples need to belong to the same platform type and the requested limit is to 100 genes and 20 samples. The files are exported as csv and can be loaded and used by the user to perform their own downstream analysis.

#### 2.7 Similar comparisons

For a given comparison, this functionality helps identify other similar comparisons.

##### 2.7.1 Similar comparisons (GO)

To identify similar comparisons based on GO terms in up or downregulated genes.

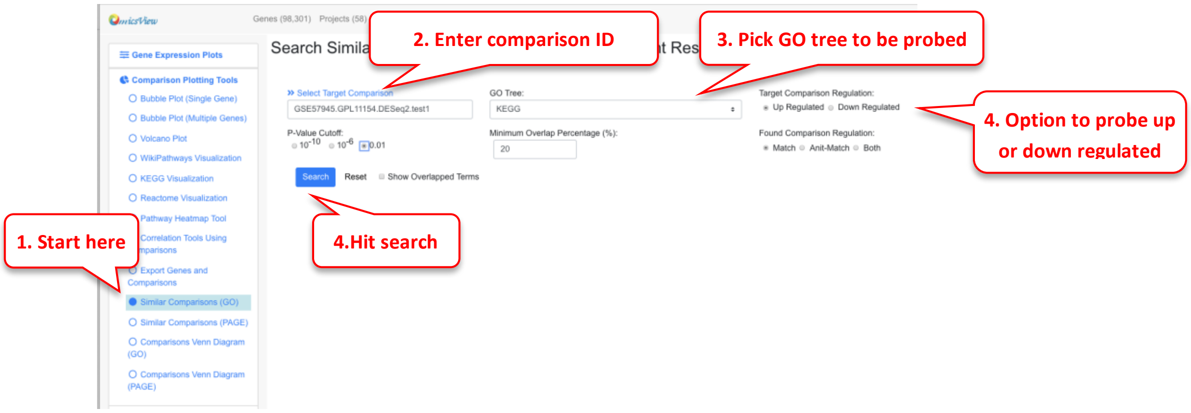

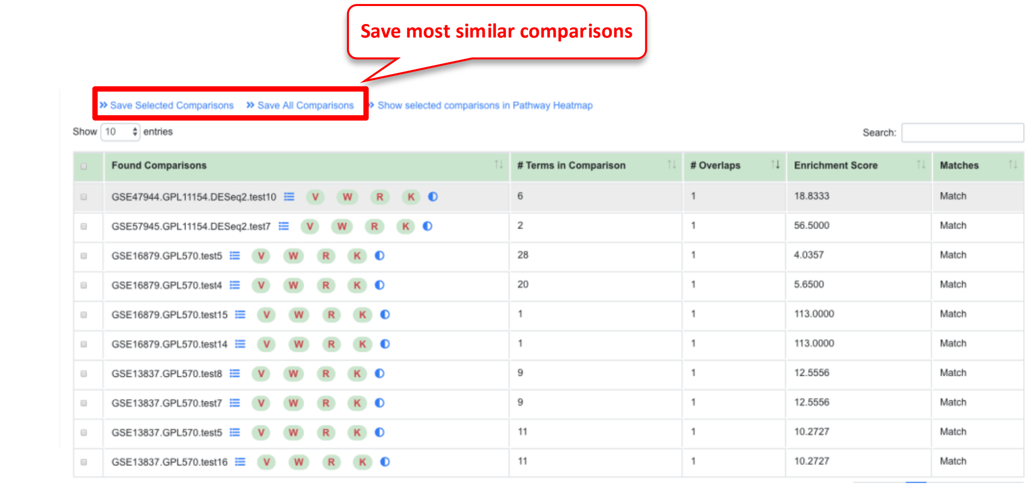

| Found Comparisons | # Terms in Comparison | # Overlaps | Enrichment Score | Matches |
|---|---|---|---|---|
| GSE47944.GPL11154.DESeq2.test10 | 6 | 1 | 18.8333 | Match |
| GSE57945.GPL11154.DESeq2.test7 | 2 | 1 | 56.5000 | Match |
| GSE16879.GPL570.test5 | 28 | 1 | 4.0357 | Match |
| GSE16879.GPL570.test4 | 20 | 1 | 5.6500 | Match |
| GSE16879.GPL570.test15 | 1 | 1 | 113.0000 | Match |
| GSE16879.GPL570.test14 | 1 | 1 | 113.0000 | Match |
| GSE13837.GPL570.test8 | 9 | 1 | 12.5556 | Match |
| GSE13837.GPL570.test7 | 9 | 1 | 12.5556 | Match |
| GSE13837.GPL570.test5 | 11 | 1 | 10.2727 | Match |
| GSE13837.GPL570.test16 | 11 | 1 | 10.2727 | Match |

#### 2.7.2 Similar comparisons (PAGE)

Similarly, the similar comparisons can be identified using PAGE results

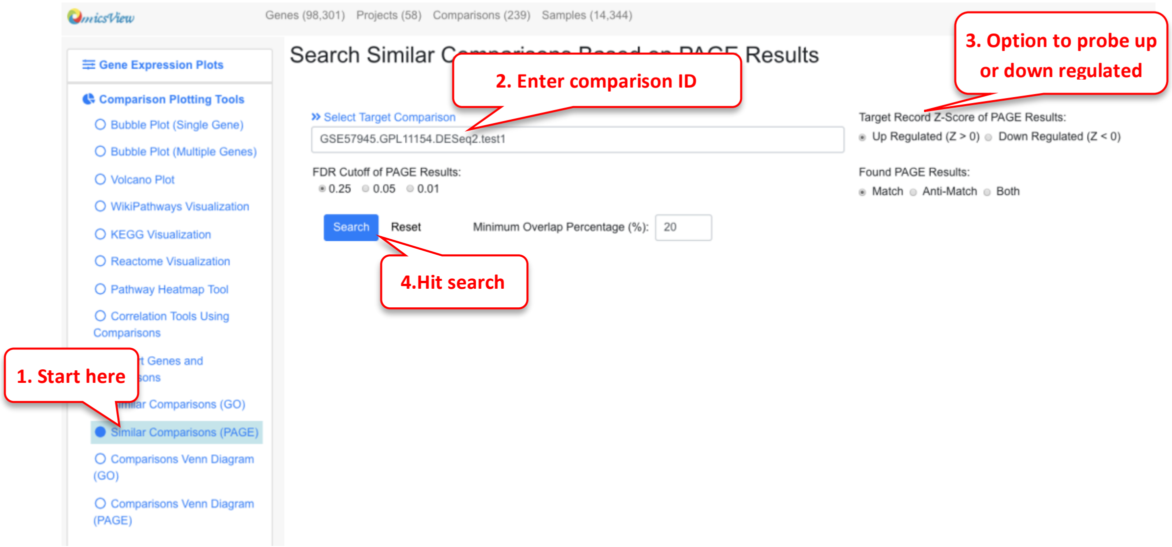

A list of comparisons that were found similar to the target comparison are displayed. They can be sorted on different attributed like # Terms in Comparison, # Overlaps and. Enrichment score

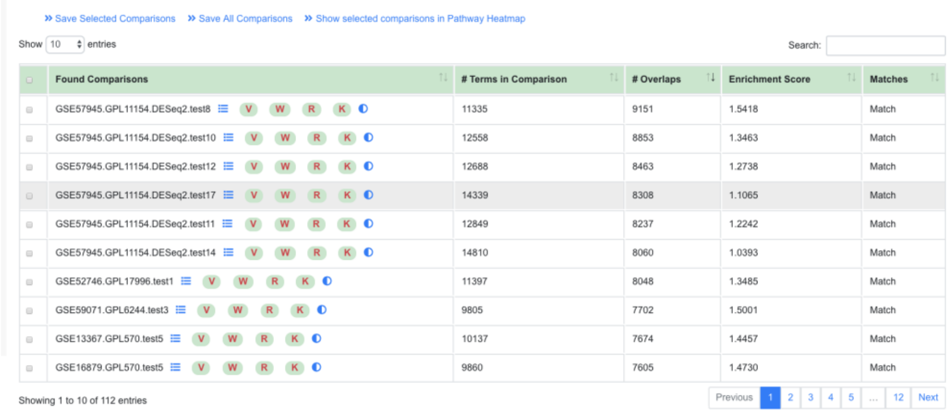

| Found Comparisons | # Terms in Comparison | # Overlaps | Enrichment Score | Matches |
|---|---|---|---|---|
| GSE57945.GPL11154.DESeq2.test8 | 11335 | 9151 | 1.5418 | Match |
| GSE57945.GPL11154.DESeq2.test10 | 12558 | 8853 | 1.3463 | Match |
| GSE57945.GPL11154.DESeq2.test12 | 12688 | 8463 | 1.2738 | Match |
| GSE57945.GPL11154.DESeq2.test17 | 14339 | 8308 | 1.1065 | Match |
| GSE57945.GPL11154.DESeq2.test11 | 12849 | 8237 | 1.2242 | Match |
| GSE57945.GPL11154.DESeq2.test14 | 14810 | 8060 | 1.0393 | Match |
| GSE52746.GPL17996.test1 | 11397 | 8048 | 1.3485 | Match |
| GSE59071.GPL6244.test3 | 9805 | 7702 | 1.5001 | Match |
| GSE13367.GPL570.test5 | 10137 | 7674 | 1.4457 | Match |
| GSE16879.GPL570.test5 | 9860 | 7605 | 1.4730 | Match |

#### 2.8 Comparisons Venn Diagrams

A good way to identify similarities and difference between different comparisons is by using the Comparisons Venn Diagram functionality. There are two ways to use this tool: By GO terms and PAGE results.

##### 2.8.1 Comparisons Venn Diagram (GO)

For identifying similar GO terms between comparisons either in the Up and Down regulated categories, this functionality can be used. In this example, 3 comparisons from Psoriasis are probed, the user also has the option of probing 2 comparisons by leaving the third field empty.

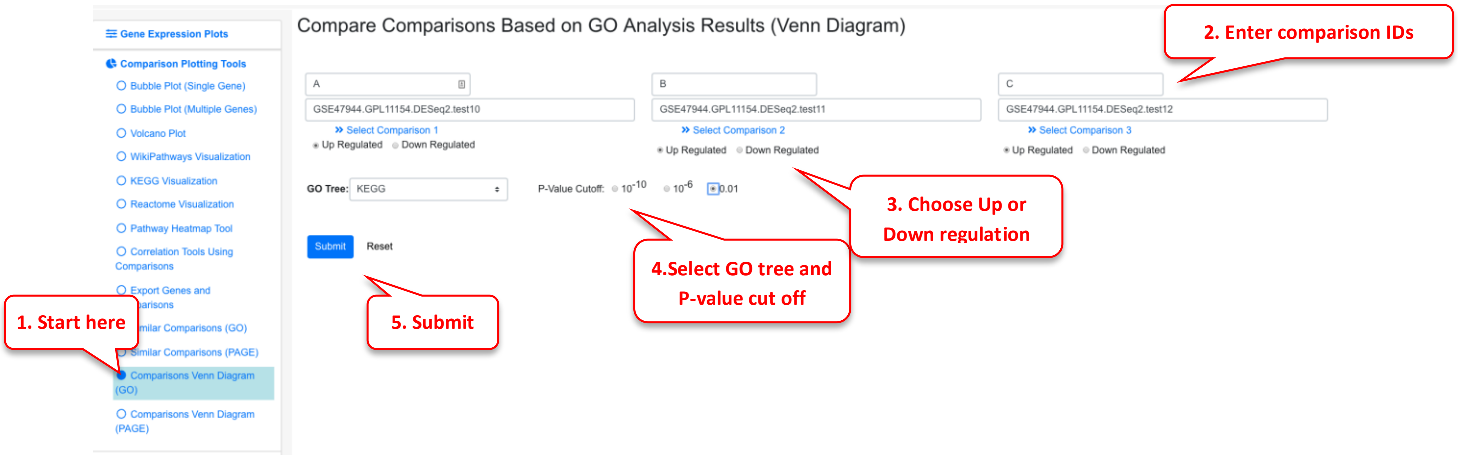

Upon finishing, an overlap summary is generated. This contains a Venn Diagram of the intersection of all the GO terms between the 3 comparisons. The count number can be clicked, as shown below, to view which terms are included.

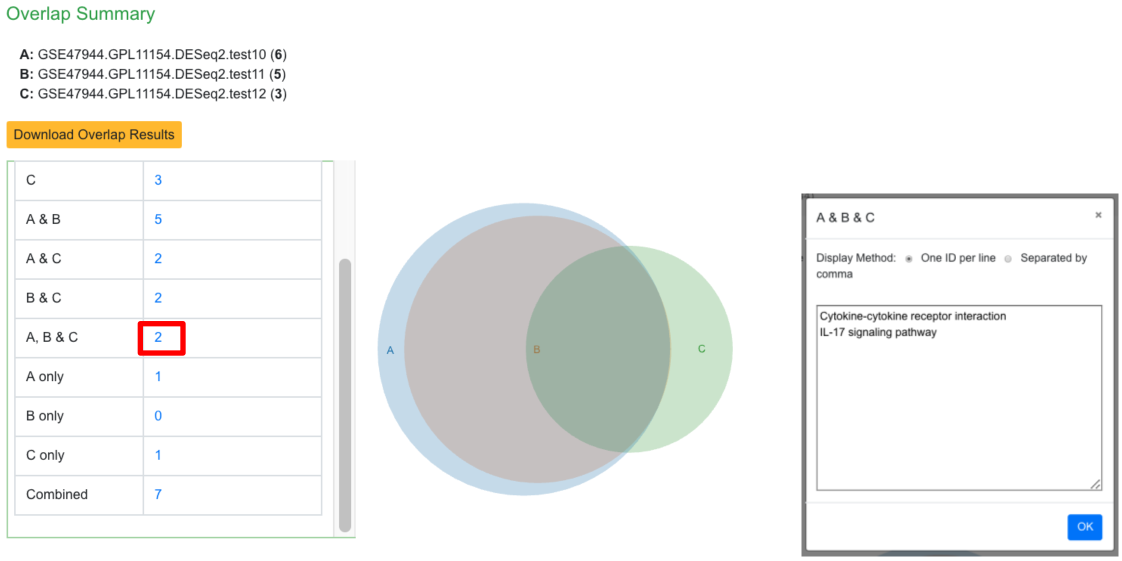

Furthermore, there is another section below this where pairs of the comparisons can be probed. Here, just A & B are displayed.

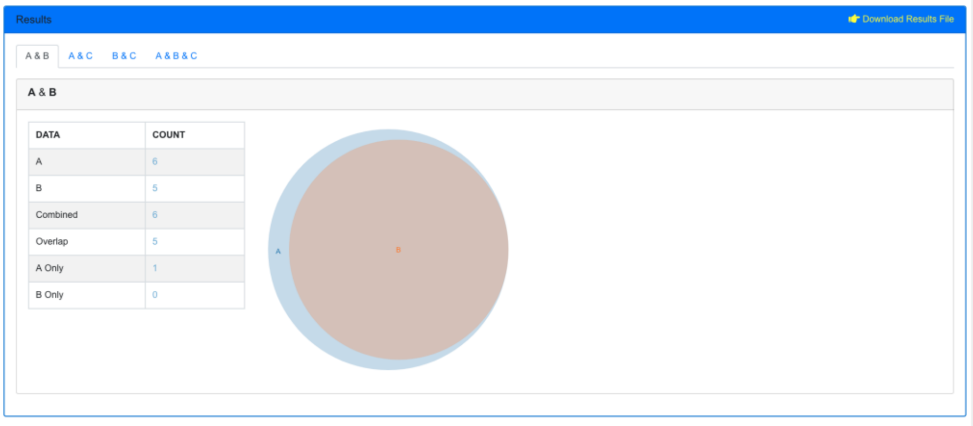

#### 2.8.2 Comparisons Venn Diagram (PAGE)

Similarly, the user can probe the similarities between PAGE results or 2 or 3 comparisons. In this example we probe comparisons from Psoriasis again.

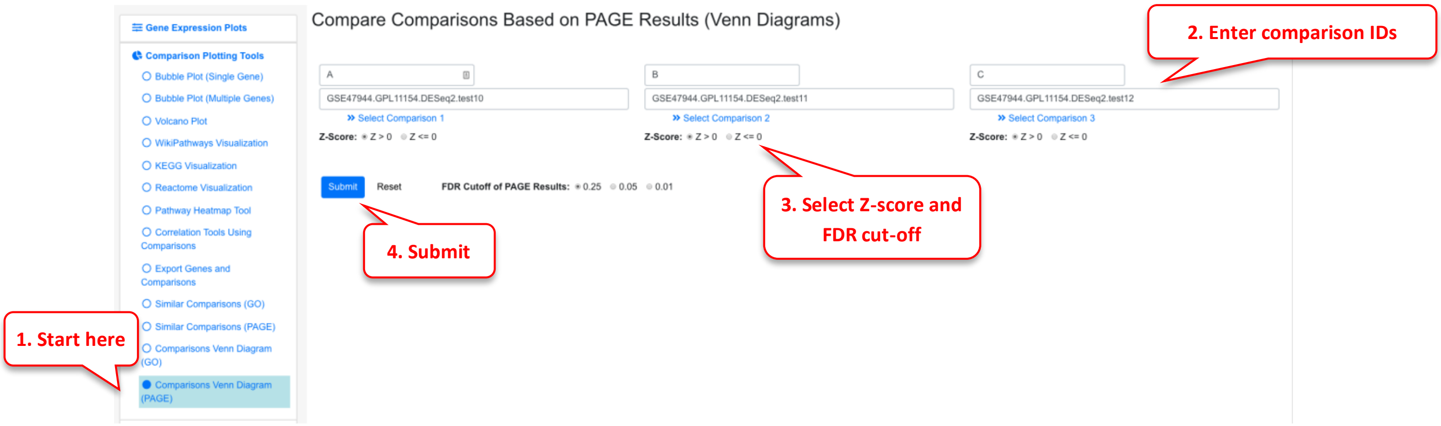

Upon finishing an overlap summary is generated which shows the intersection of the PAGE results between the comparisons.

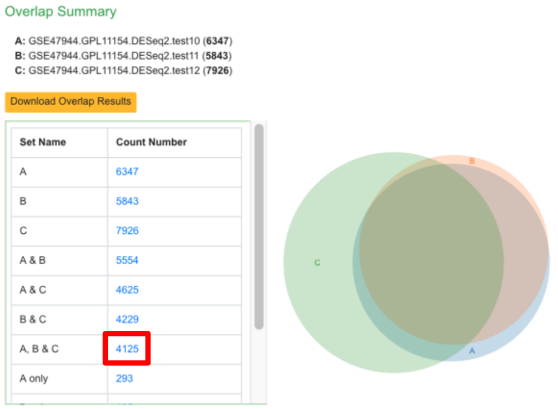

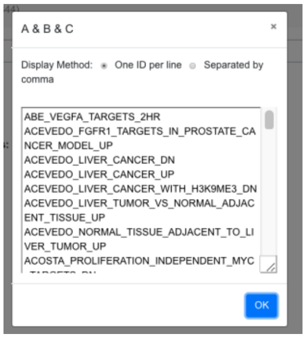

### 3 Visualize Comparison Data

There are 235 comparison in the demo system, covering different categories including Disease vs. Normal, Treatment vs. Control, Tissue 1 vs. Tissue 2 etc.  The system offers many ways to visualize the data and help users find the most interesting data related to their study.

#### 3.1 Dashboard View of Comparison

The dashboard shows a summary of all the comparisons.

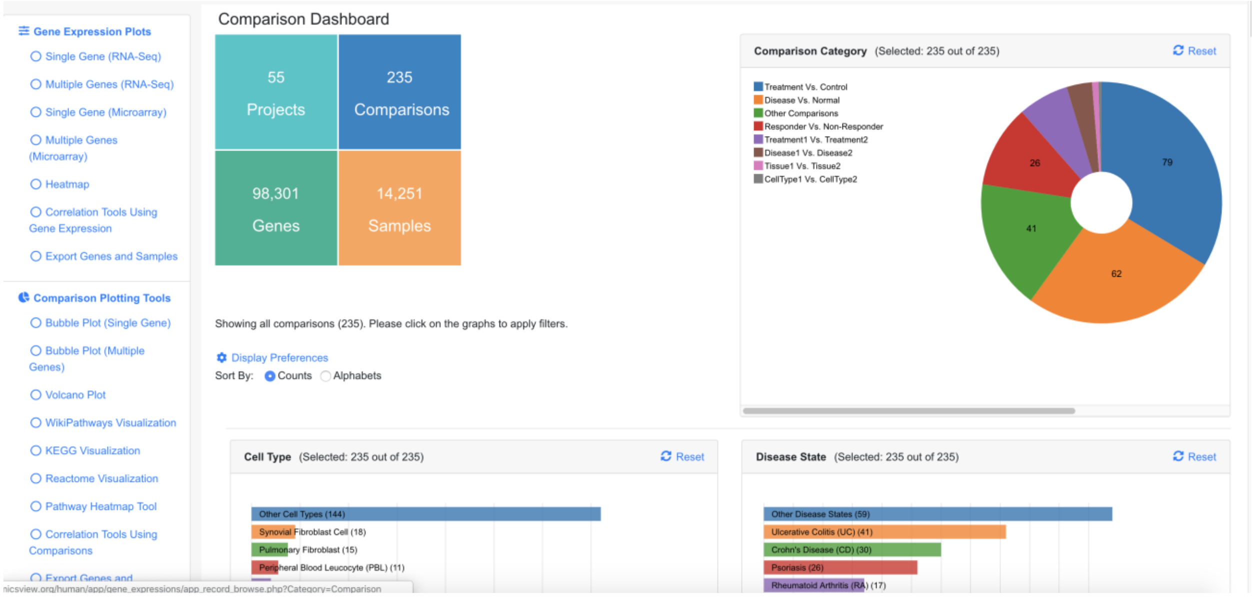

The dashboard shows Comparison Categories, Cell Type, Disease State, Treatment, Platform summaries. A table is also available below the dashboard that lists all the comparisons.

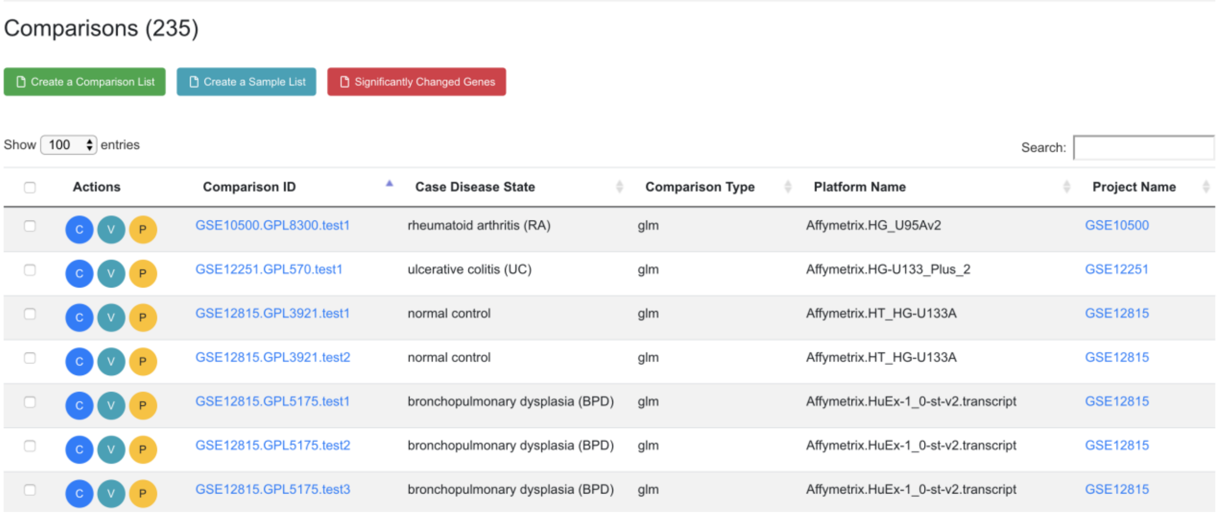

| Actions | Comparison ID | Case Disease State | Comparison Type | Platform Name | Project Name |
|---|---|---|---|---|---|
| C V P | GSE10500.GPL8300.test1 | rheumatoid arthritis (RA) | glm | Affymetrix.HG_U95Av2 | GSE10500 |
| C V P | GSE12251.GPL570.test1 | ulcerative colitis (UC) | glm | Affymetrix.HG-U133_Plus_2 | GSE12251 |
| C V P | GSE12815.GPL3921.test1 | normal control | glm | Affymetrix.HT_HG-U133A | GSE12815 |
| C V P | GSE12815.GPL3921.test2 | normal control | glm | Affymetrix.HT_HG-U133A | GSE12815 |
| C V P | GSE12815.GPL5175.test1 | bronchopulmonary dysplasia (BPD) | glm | Affymetrix.HuEx-1_0-st-v2.transcript | GSE12815 |
| C V P | GSE12815.GPL5175.test2 | bronchopulmonary dysplasia (BPD) | glm | Affymetrix.HuEx-1_0-st-v2.transcript | GSE12815 |
| C V P | GSE12815.GPL5175.test3 | bronchopulmonary dysplasia (BPD) | glm | Affymetrix.HuEx-1_0-st-v2.transcript | GSE12815 |

#### 3.2 Set Dashboard Preference

Users can change how the comparison summary is displayed by following the red numbers.

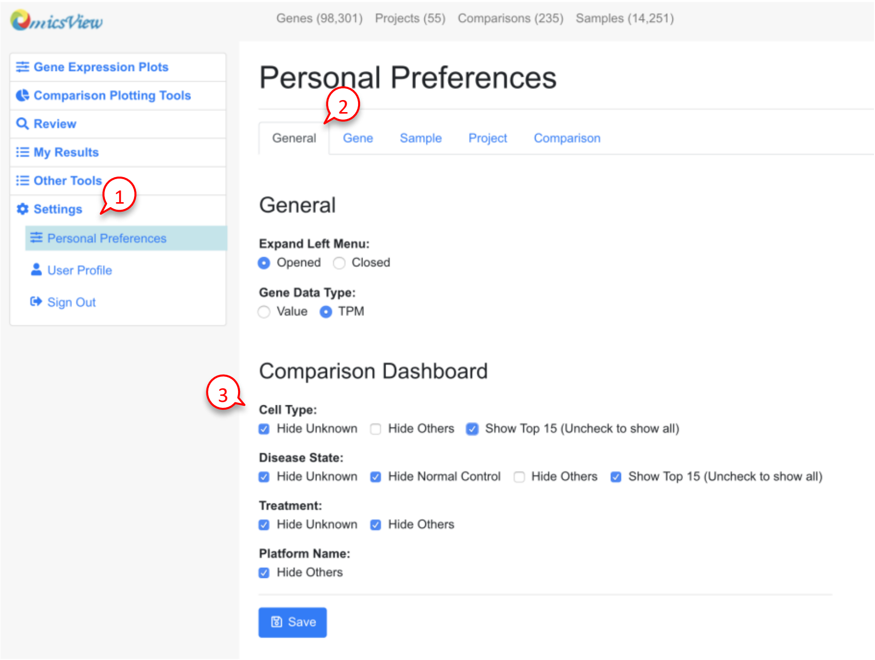

Since there are many Cell Types and Disease States, the user can choose to display only the top 15 categories, or all categories.  See example screenshots below.

| Show top 15 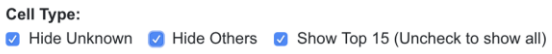  | Show all cell types 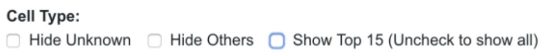  |
|---|---|
| 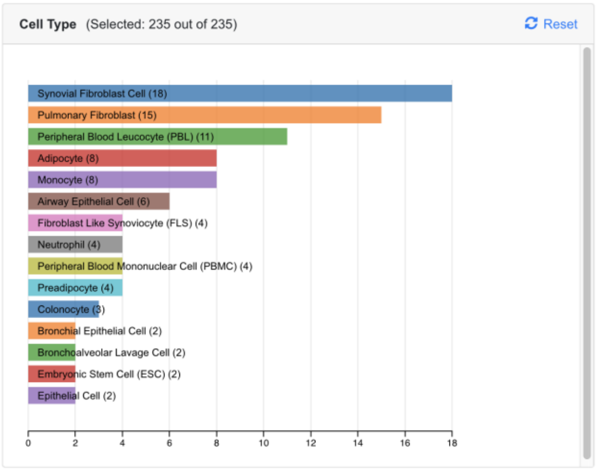  | 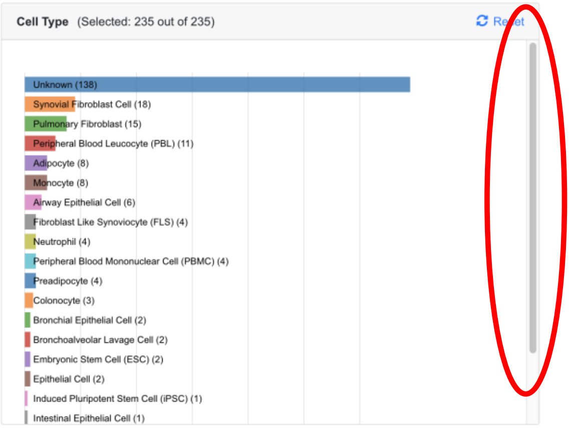  In this view, Users need to use the scroll bar at the right to move up and down to view all cell types. |

When the user chooses only the top 15 categories, all other categories are shown as Other Cell Type. There are other data points where the Cell Type is unknown from the study. Two categories, denoted as "Other" and "Unknown" can be hidden by the user in the preference.

#### 3.3 Dynamic Filtering of Comparisons on Dashboard

The user can click any chart in the dashboard to focus on one or more categories that are of interest to the study.

In the example below, we are only interested in Disease vs. Normal comparisons, and we further narrow down the data to three disease areas: ulcerative colitis, crohn's disease, and rheumatoid arthritis. Now we are looking at 36 out of 199 comparisons. The table at the bottom of the page shows details of these 36 comparisons. The "Reset All Charts" link above the statistics panel is used to show all comparisons again.

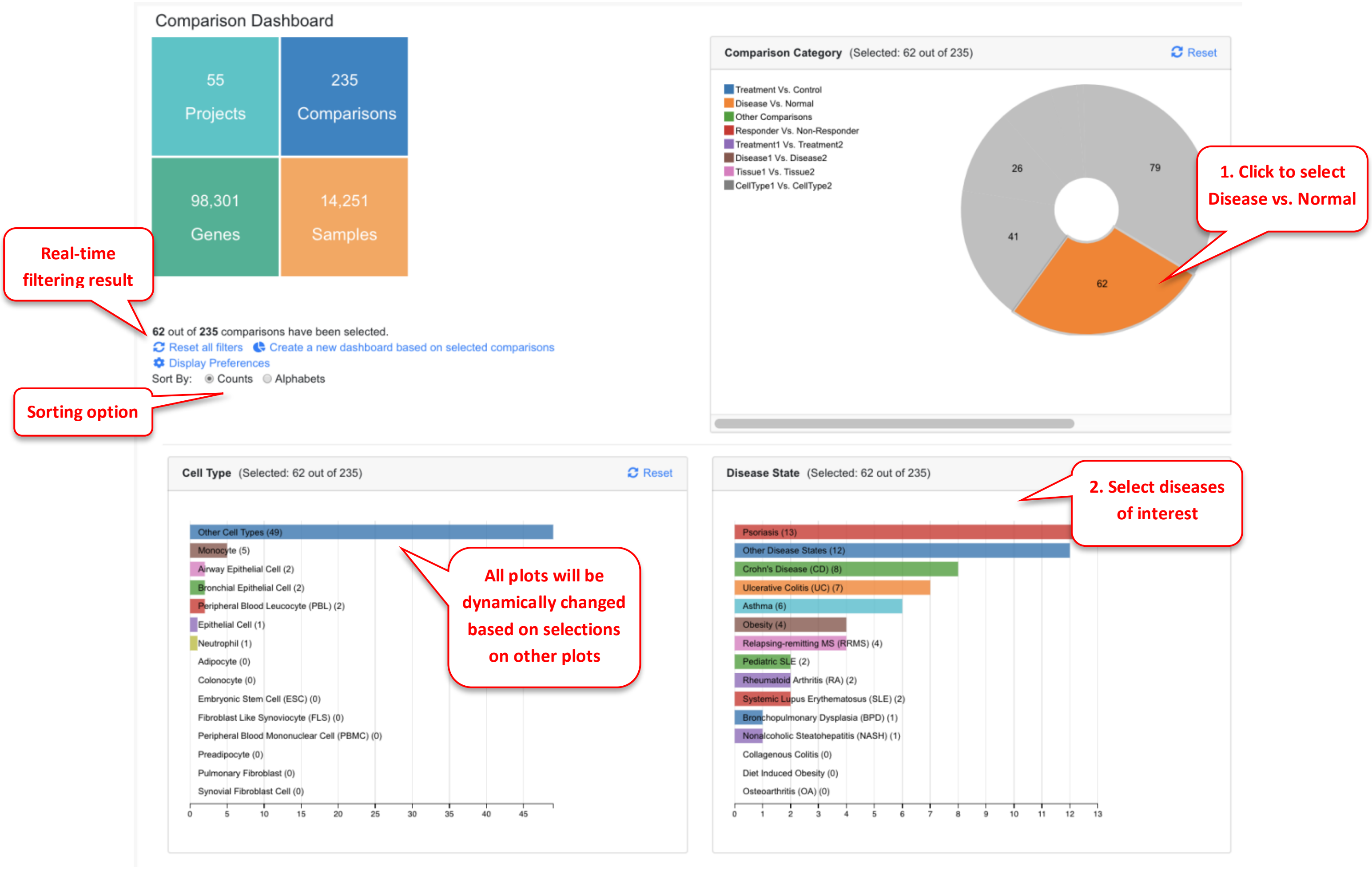

#### 3.4 Bubble Plot of Comparisons Associated with a Single Gene

For each gene, Users can view all the available comparisons in a bubble chart.

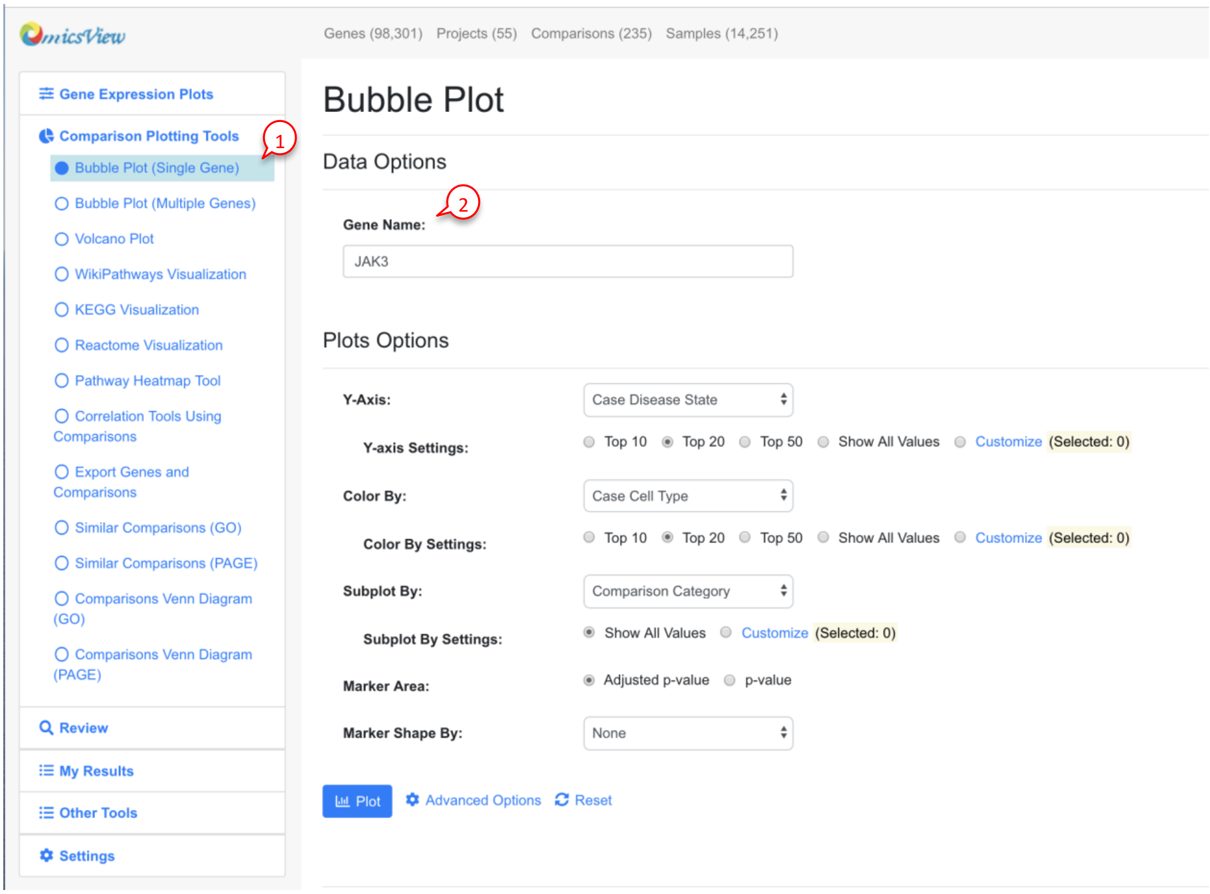

The default settings work for most users. After clicking the Plot button, you will see a plot like below:

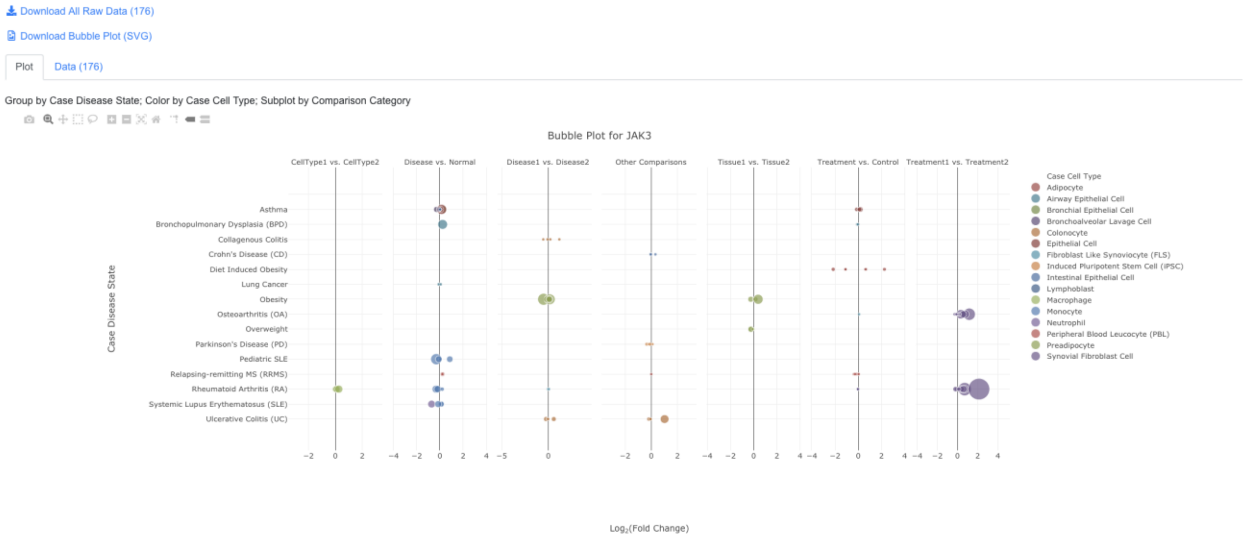

In the bubble plot, the X-axis shows log2 Fold Change of the comparison, the Y-axis shows diseases state. Each dot represents the comparison result of this gene from one comparison. The color of the dot represents cell type, and the size of the dot represents significance (-log10(FDR), larger is more significant).

The user can click and unclick the color legend at right to select or deselect cell types. When mouse over a dot, more details are shown. And the user can also click the dot to link to other graphs.

The tool bars at top right corner allows the user to zoom and pan the graph.

#### 3.5 Data Filter and Advanced Settings in Bubble Plot

In addition, advanced users can change settings by click "Modify Settings Button". For example, the user may want to show a selected list of diseases. After clicking Customize in Case_DiseaseState, user can select which diseases to display in the pop-up window.

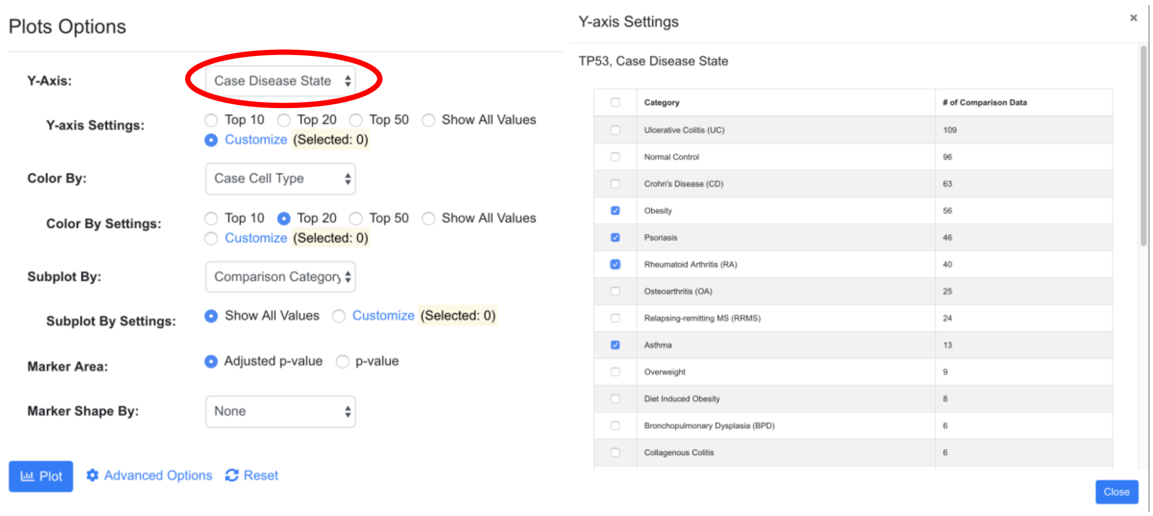

After modifying the setting, the user can click plot button to view the new chart. The system will display how many data points are chosen based on the filter.

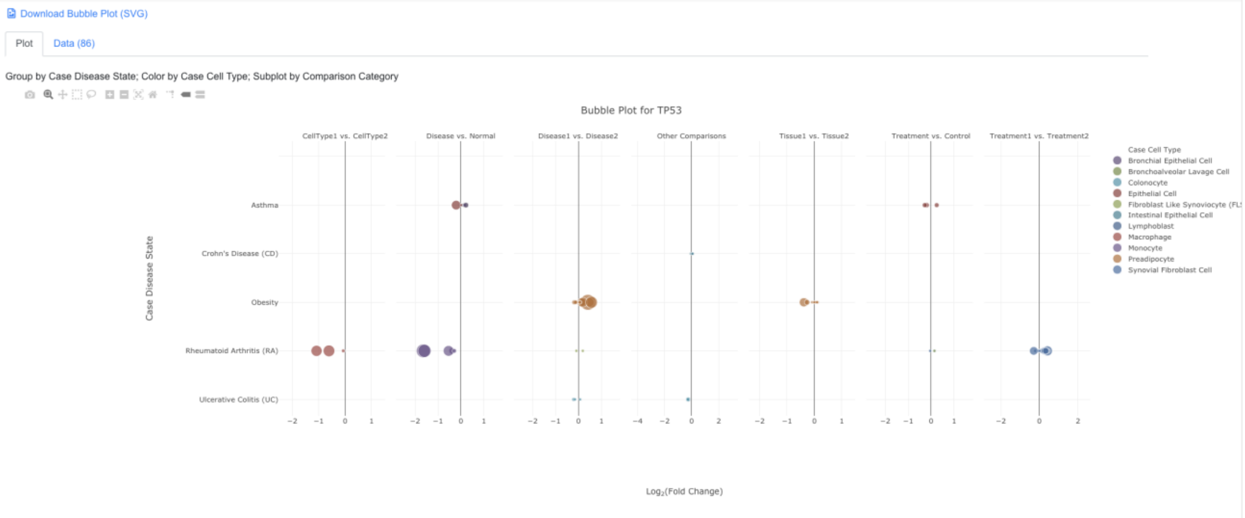

#### 3.6 Bubble Plot of Sets of Genes and Comparisons

It can be useful to look at a set of genes (e.g. all differentially expressed genes, or genes from a certain pathways) in a set of related comparisons (e.g. all from the same disease).

To view this type of bubble plot, select Bubble Plot (Multiple Genes).

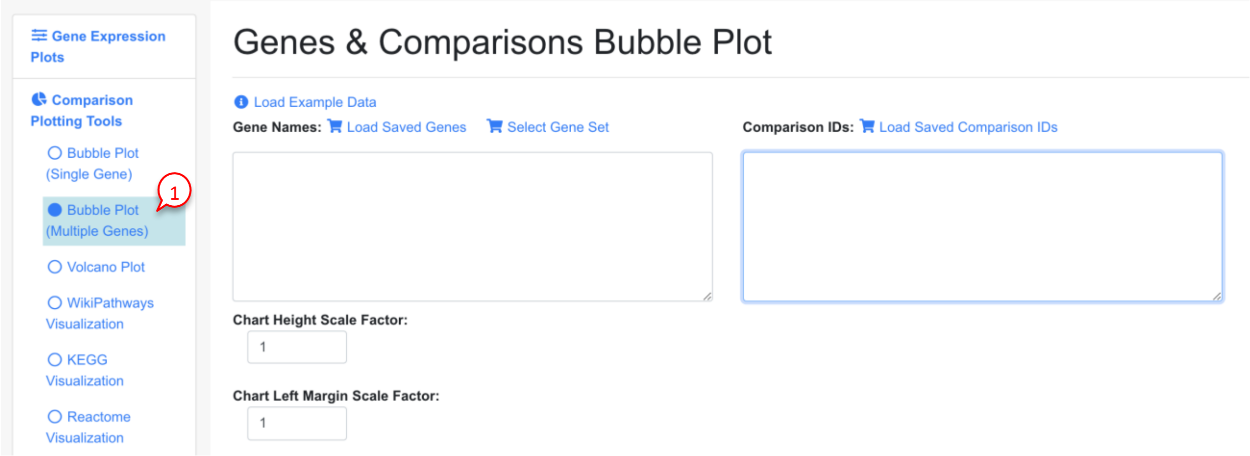

In the Genes and Comparisons Bubble plot window, Users can now enter the symbols of the genes, and the comparison names. However, it is much easier to use the saved genes and saved comparisons features, or other tools from the system to quickly get a get set.

In the example below, we use dashboard to select 8 comparisons that are for Disease vs. Normal in crohn's disease (CD). We save the comparisons and load in the bubble plot tool. For gene list, we get the up-regulated immune response genes from comparison GSE57945.GPL11154.DESeq2.test1 and paste into the gene names fields.

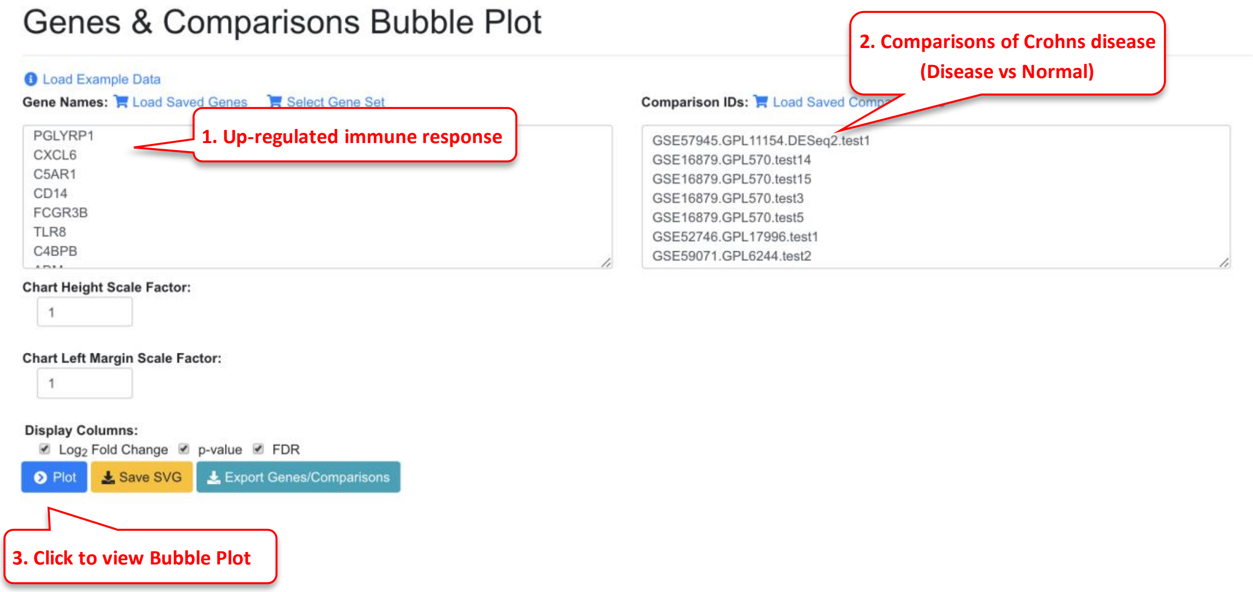

In the bubble plot, the gene symbols are listed in Y-axis. The X-axis represents logFC, and color of the bubble represents comparison; the size of the bubble represents the significance.

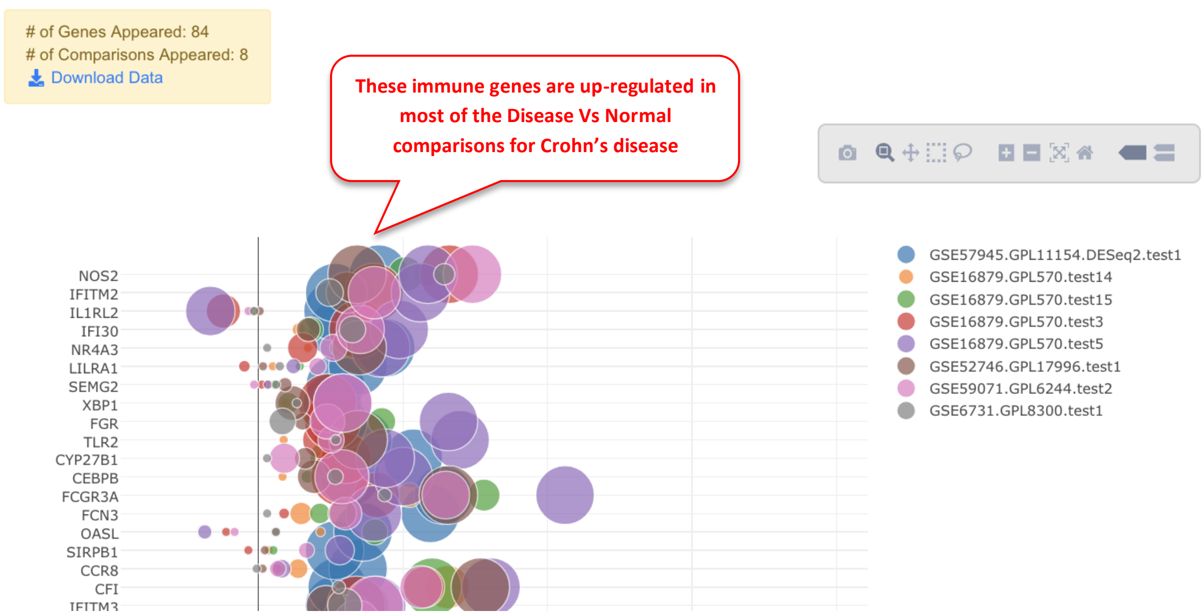

In the legend, the color keys for comparisons are shown. Users can click the color key in the legend to hide/show comparisons. The size of the color dot in the legend correlates to the largest bubble for that comparison, which is the most significant gene with the smallest FDR.

#### 3.7 Get significant genes from comparisons

Another way to get a gene set to visualize in the genes/comparisons bubble plot is to filter for significantly changed genes. To do this, first select a few comparisons from the dash board, and click the "View Significantly Changed Genes" button.

Dashboard filter:

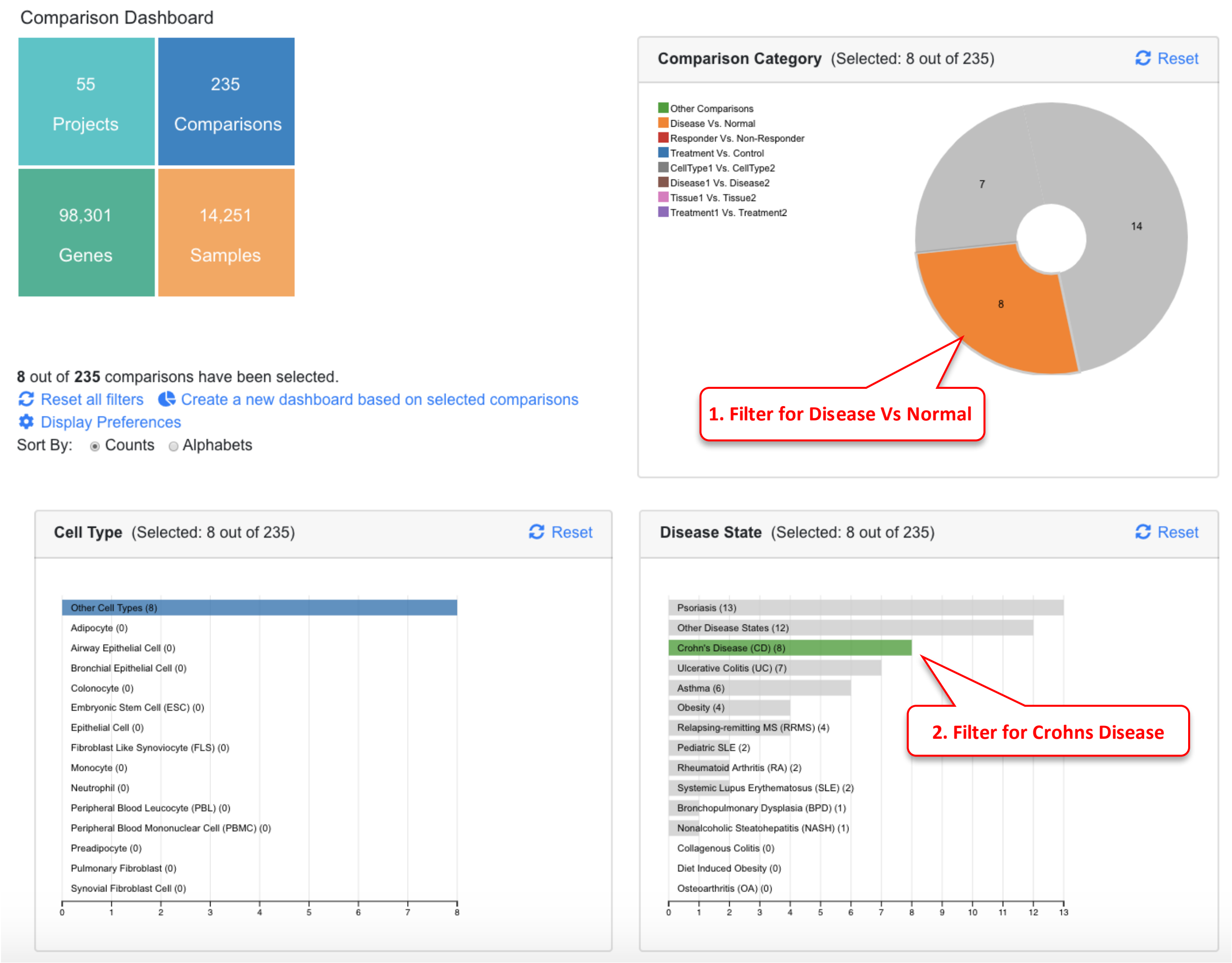

In table, select comparisons and view significantly changed genes.

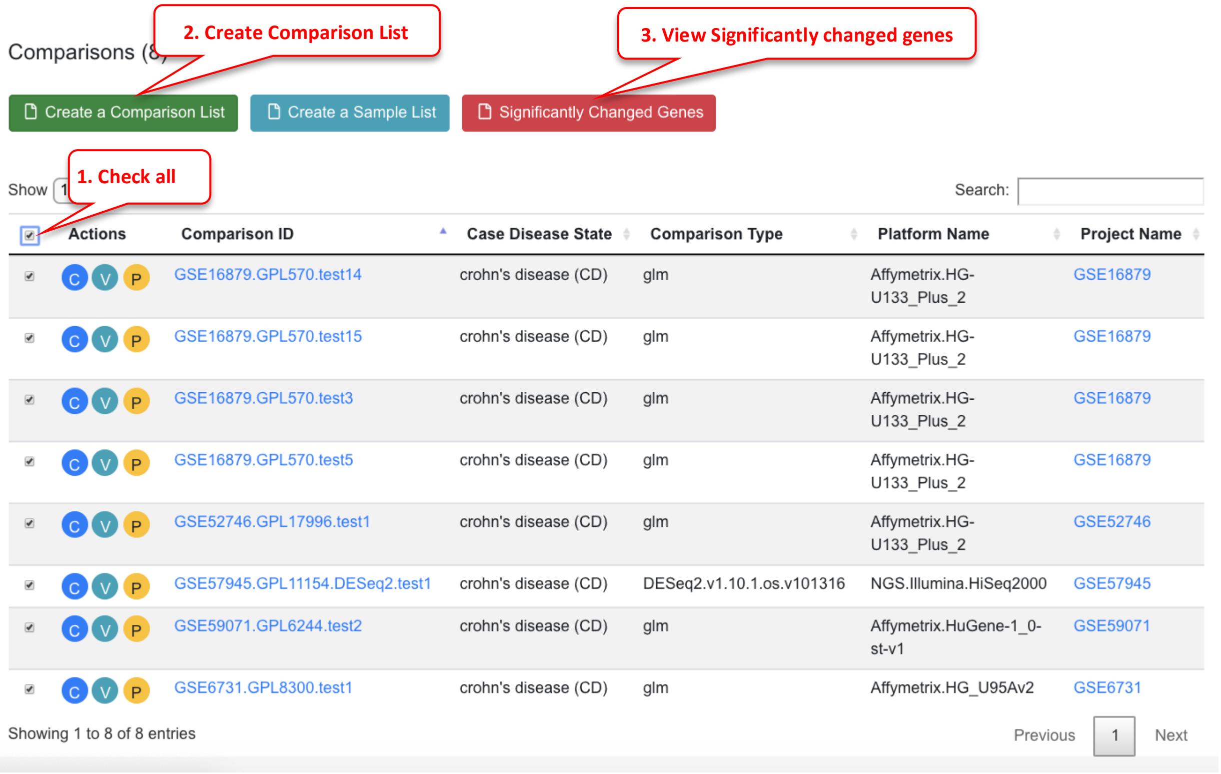

| | Actions | Comparison ID | Case Disease State | Comparison Type | Platform Name | Project Name |
|---|---|---|---|---|---|---|
| ☑ | C V P | GSE16879.GPL570.test14 | crohn's disease (CD) | glm | Affymetrix.HG-U133_Plus_2 | GSE16879 |
| ☑ | C V P | GSE16879.GPL570.test15 | crohn's disease (CD) | glm | Affymetrix.HG-U133_Plus_2 | GSE16879 |
| ☑ | C V P | GSE16879.GPL570.test3 | crohn's disease (CD) | glm | Affymetrix.HG-U133_Plus_2 | GSE16879 |
| ☑ | C V P | GSE16879.GPL570.test5 | crohn's disease (CD) | glm | Affymetrix.HG-U133_Plus_2 | GSE16879 |
| ☑ | C V P | GSE52746.GPL17996.test1 | crohn's disease (CD) | glm | Affymetrix.HG-U133_Plus_2 | GSE52746 |
| ☑ | C V P | GSE57945.GPL11154.DESeq2.test1 | crohn's disease (CD) | DESeq2.v1.10.1.os.v101316 | NGS.Illumina.HiSeq2000 | GSE57945 |
| ☑ | C V P | GSE59071.GPL6244.test2 | crohn's disease (CD) | glm | Affymetrix.HuGene-1_0-st-v1 | GSE59071 |
| ☑ | C V P | GSE6731.GPL8300.test1 | crohn's disease (CD) | glm | Affymetrix.HG_U95Av2 | GSE6731 |

In the significantly Changed Genes window, the comparisons from the previous page are already loaded. You can add or remove comparisons if needed.

Now select direction (up-, down-, or both), and use the logFC cutoff and FDR value to get a list of genes. Depending on the comparisons, sometimes you may need to adjust the logFC and FDR values to get a good list of genes. In general, for bubble plot, using <100 genes will make the graph easier to read.

Once you are happy with the gene list, you can save it. You can also export the list for later use.

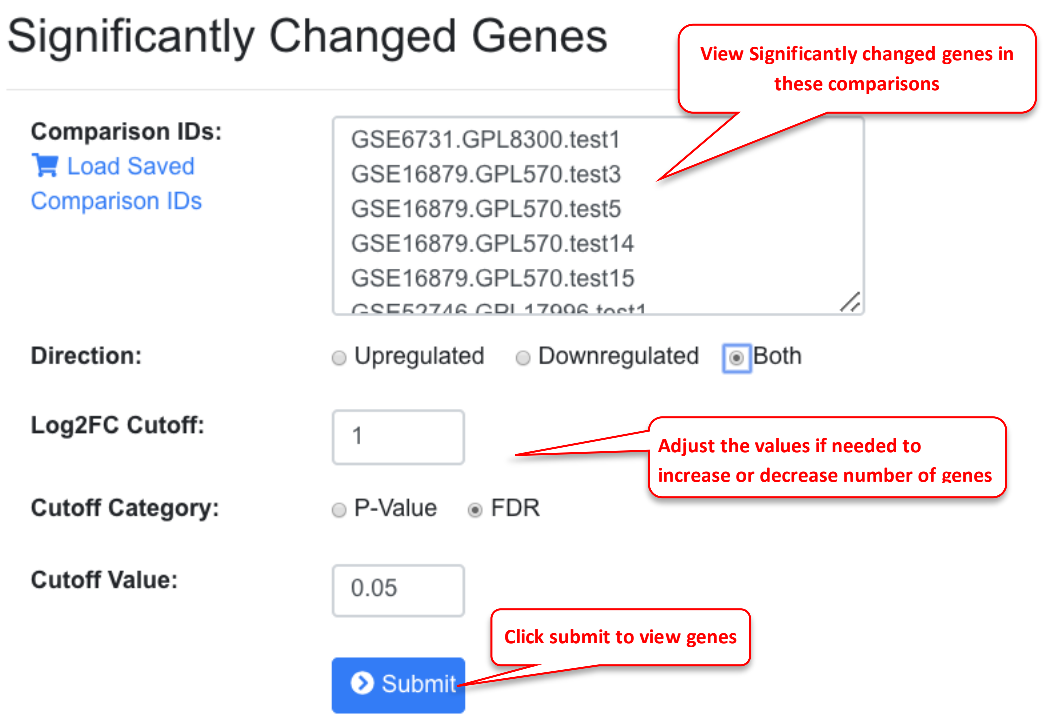

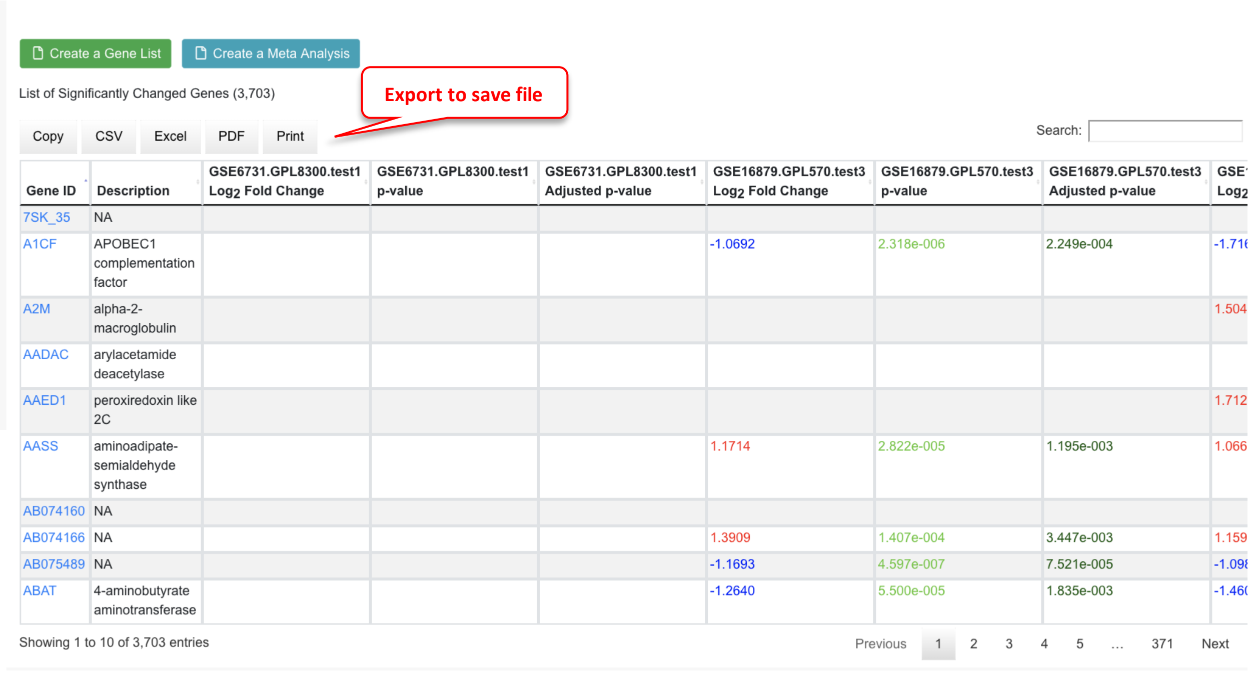

| Gene ID | Description | GSE6731.GPL8300.test1 Log2 Fold Change | GSE6731.GPL8300.test1 p-value | GSE6731.GPL8300.test1 Adjusted p-value | GSE16879.GPL570.test3 Log2 Fold Change | GSE16879.GPL570.test3 p-value | GSE16879.GPL570.test3 Adjusted p-value | GSE Log2 |
|---|---|---|---|---|---|---|---|---|
| 7SK_35 | NA | | | | | | | |
| A1CF | APOBEC1 complementation factor | | | | -1.0692 | 2.318e-006 | 2.249e-004 | -1.71 |
| A2M | alpha-2-macroglobulin | | | | | | | 1.504 |
| AADAC | arylacetamide deacetylase | | | | | | | |
| AAED1 | peroxiredoxin like 2C | | | | | | | 1.712 |
| AASS | aminoadipate-semialdehyde synthase | | | | 1.1714 | 2.822e-005 | 1.195e-003 | 1.066 |
| AB074160 | NA | | | | | | | |
| AB074166 | NA | | | | 1.3909 | 1.407e-004 | 3.447e-003 | 1.159 |
| AB075489 | NA | | | | -1.1693 | 4.597e-007 | 7.521e-005 | -1.09 |
| ABAT | 4-aminobutyrate aminotransferase | | | | -1.2640 | 5.500e-005 | 1.835e-003 | -1.46 |

#### 3.8 View Significantly Changed Genes in Bubble Plot

Back to the bubble plot, Users can load the saved comparisons and saved genes and view the plot.

In the example below, it can be seen that most significant genes come from up-regulated direction.

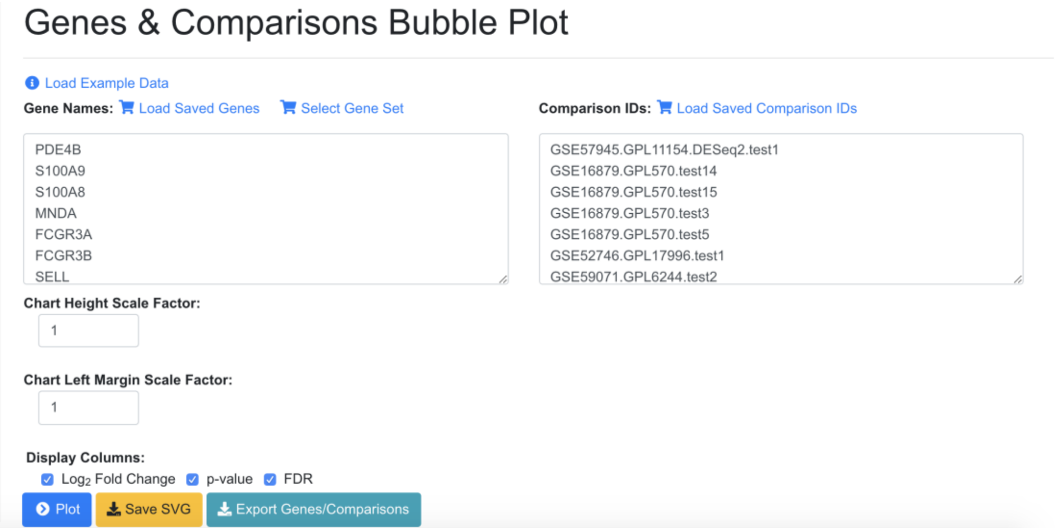

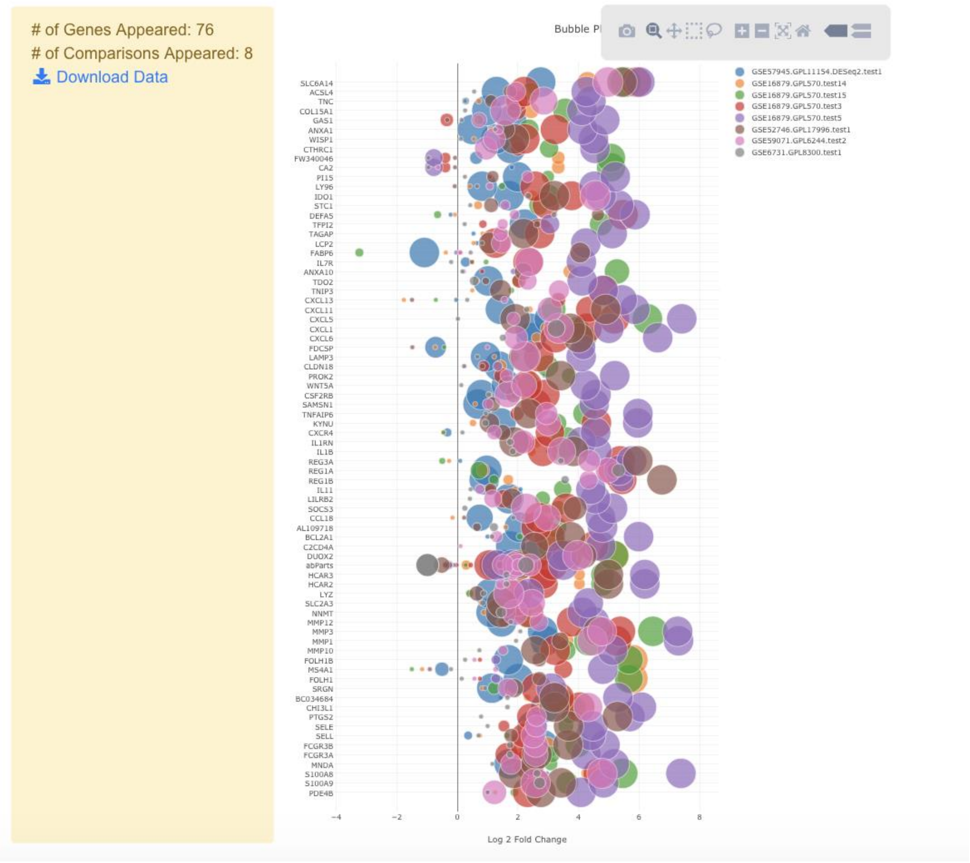

#### 3.9 Volcano Plot of a Comparison

Volcano plot is useful to view a top level summary of how many genes are significantly up- or down-regulated in a comparison.

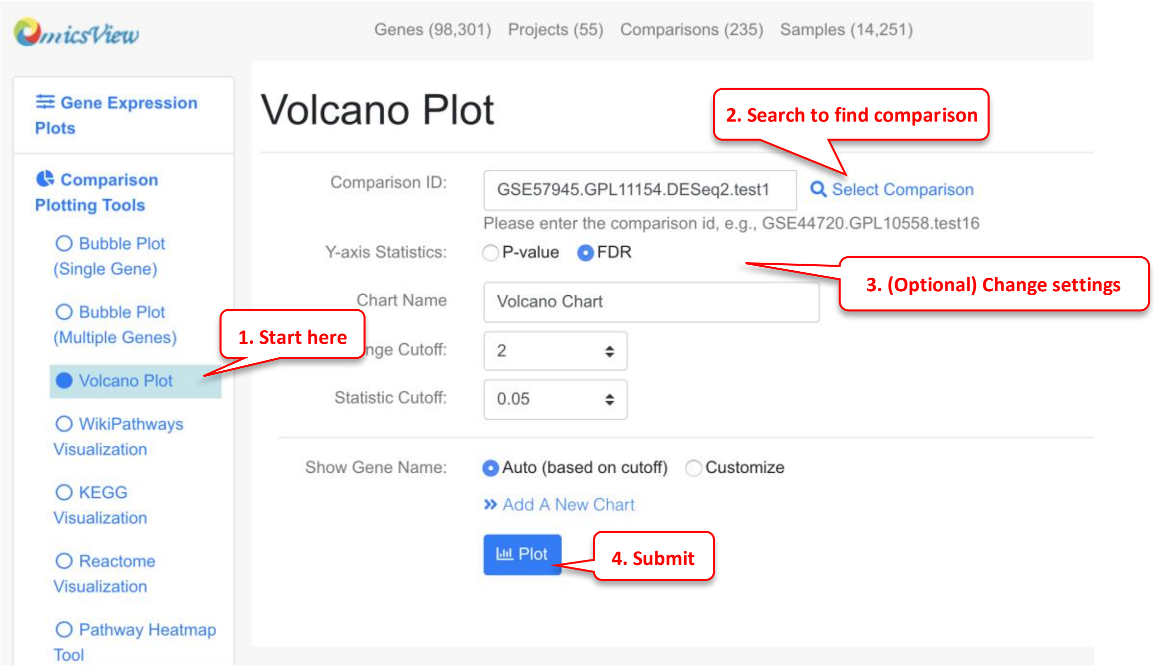

Summary

| | |
|---|---|
| Comparison: | GSE57945.GPL11154.DESeq2.test1 |
| Fold Change Cutoff: | 2 |
| $Log_2$(Fold Change Cutoff): | 1.000 |
| Stat Cutoff: | 0.05 |
| $-Log_{10}$(Stat Cutoff): | 1.301 |
| # of Upregulated Genes: | 452 |
| # of Downregulated Genes: | 156 |
| All Genes: | 30,379 |

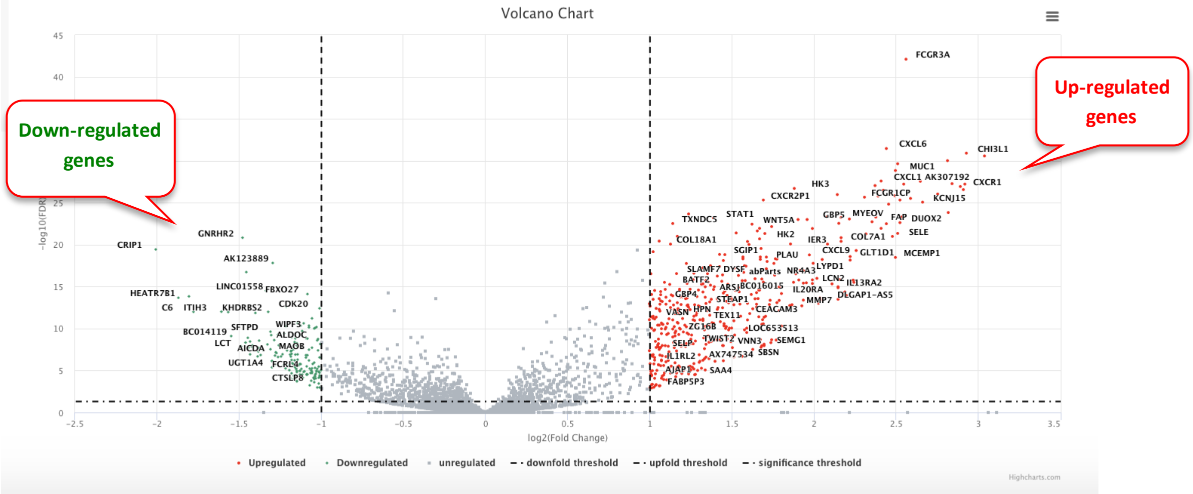

Users can use mouse to drag over an area to zoom in. Mouse over a point will show the gene details. Click the data point will show you links to other graphs.

#### 3.10 View Multiple Volcano Plots Together

Users can also show multiple comparisons side-by-side. If needed, the user can also highlight the same group of genes across the volcano plots.

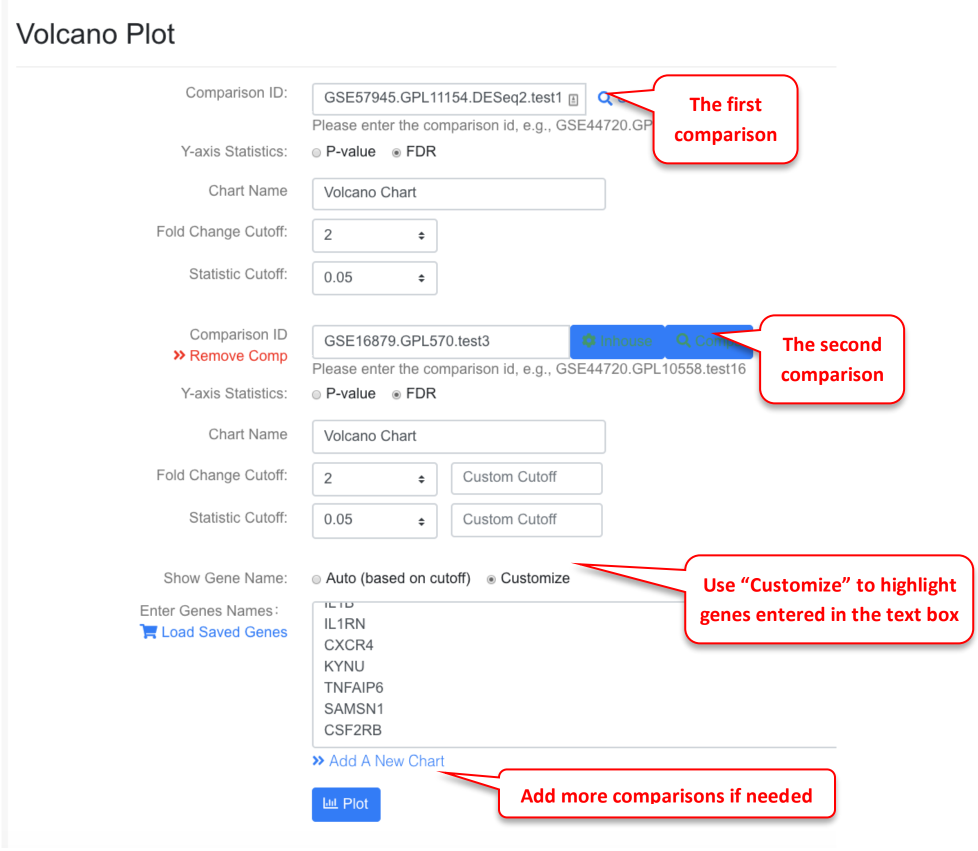

The resulting volcano plots are shown as below. Selected genes are shown as orange dots.

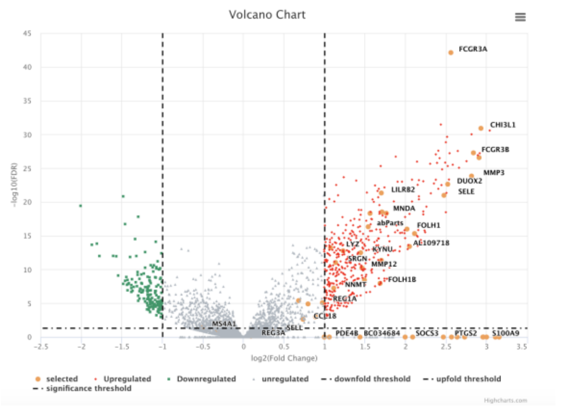

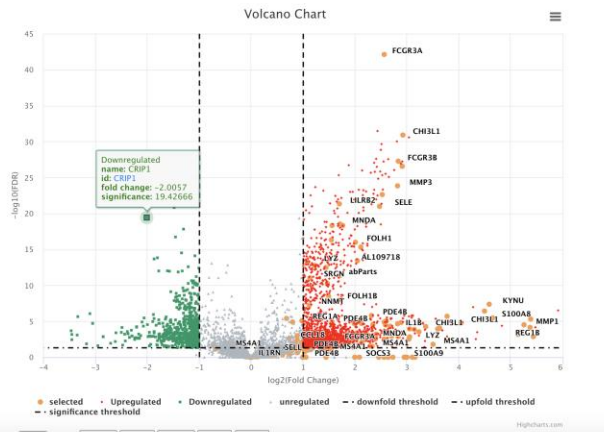

#### 3.11 Overlap Comparison Data to Pathway Graph

If Users are interested in a particular pathway, sometimes it is useful to map the RNA-Seq or microarray data to the pathway for visualization.

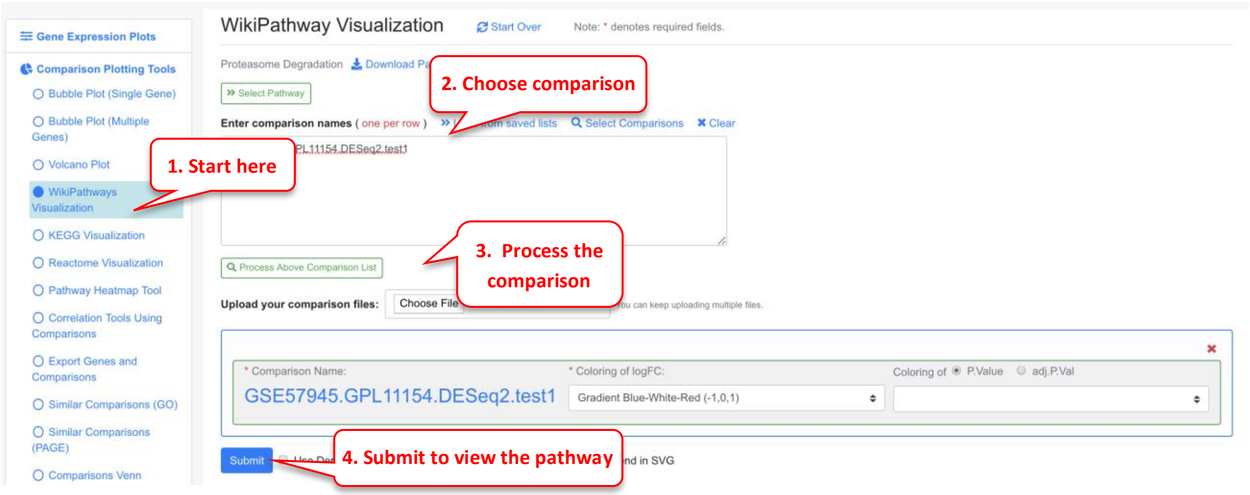

In the pathway plot, typically we use red-blue color scale to show the log2 Fold Change. Blue is down-regulated, red is up-regulated.

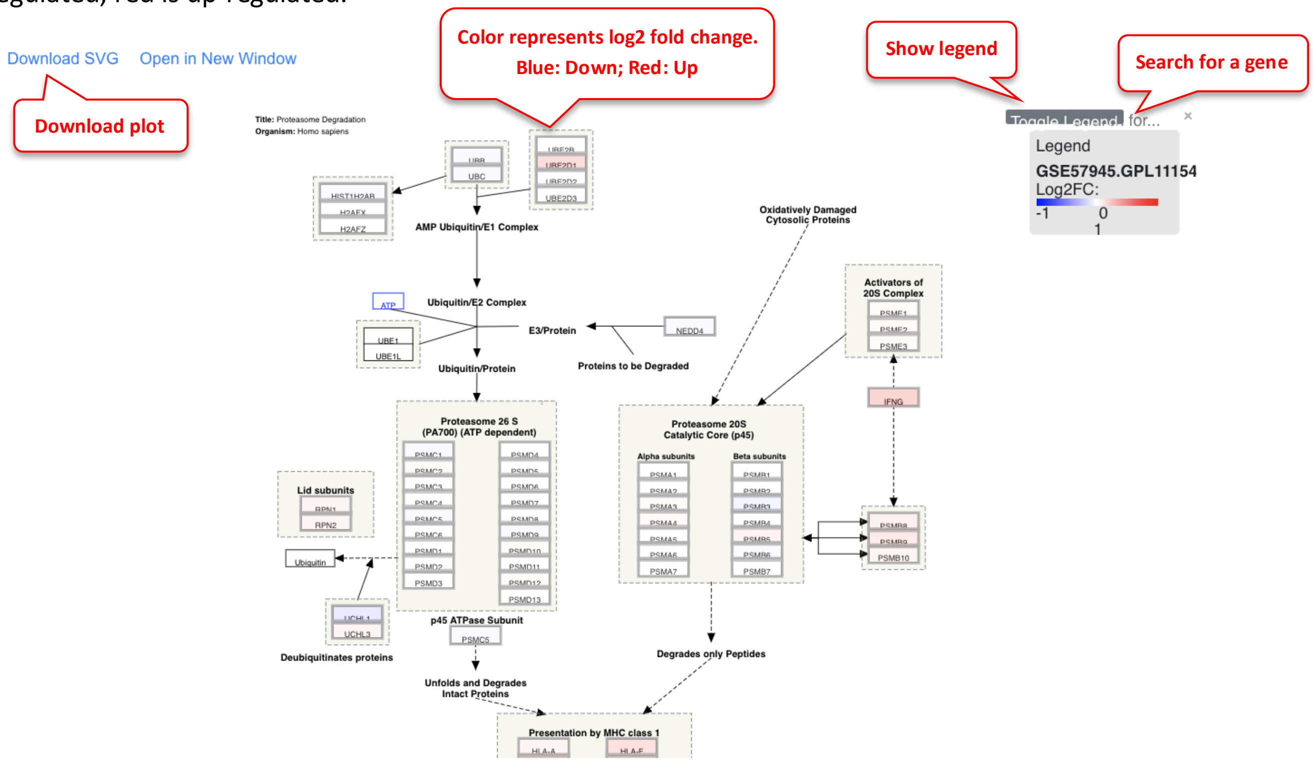

#### 3.12 Pathway Plot from Several Comparisons

The user can add multiple comparisons from the pathway plot tool by clicking Add Comparison link. This is a good functionality to overlay differential genes from multiple comparisons along a pathway.

In this example we use data from Connor-Robson et al., Neurobiology of Disease, 2019. In this paper, the authors perform proteomics (Mass spec) and RNAseq on diseased and normal iPSCs. They perform both the assays at 2 timepoints of differentiation: D35 and D56. This visualization will identify if the same genes are differential expressed in the proteomic and the RNAseq datasets, as well as in the two timepoints on a WikiPathway.

The data was uploaded onto OmicsView. For the proteomic dataset, the protein ID was converted to the gene ID. Here the 4 comparisons (2 Proteomics and 2 RNAseq) are loaded, and the P13k Akt signaling pathway is probed.

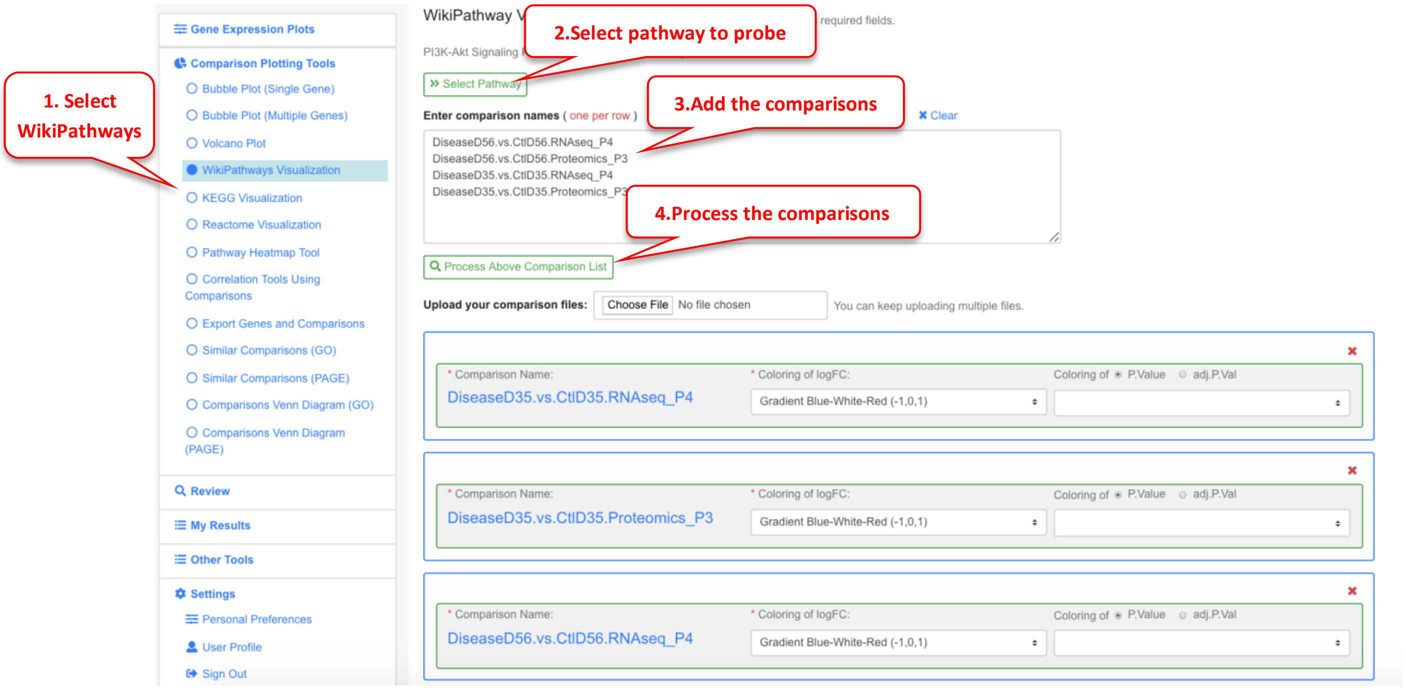

The pathway plot now has 4 color bars corresponding to the different comparisons.

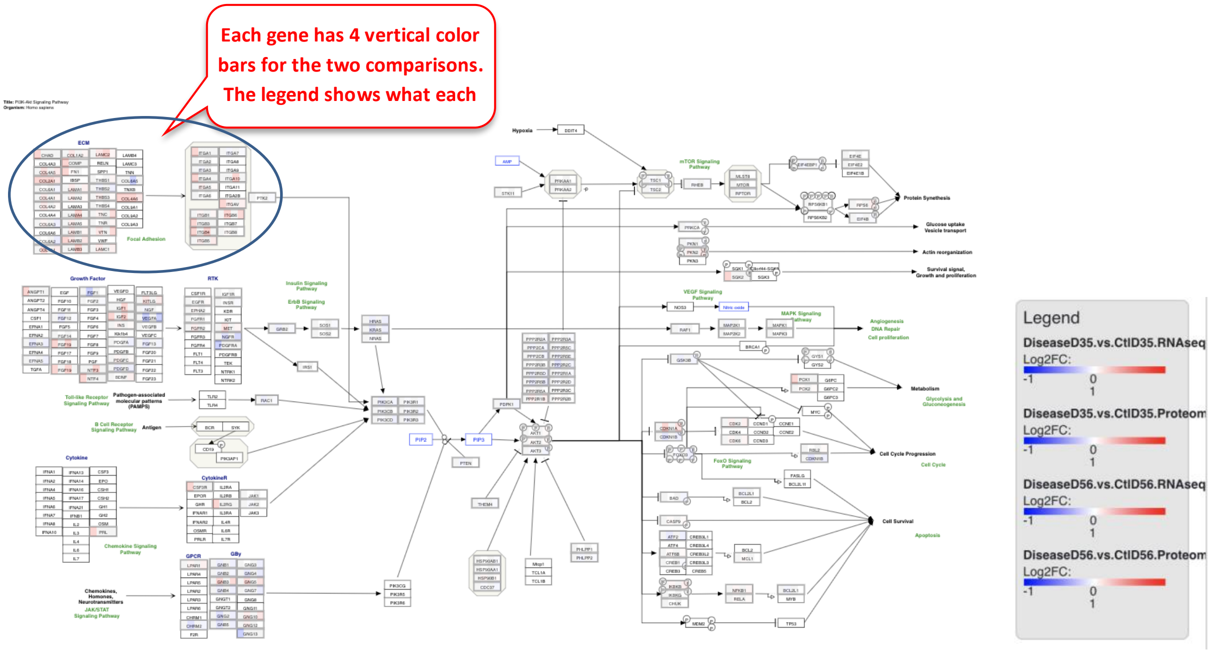

Users can also zoom into the plot for better visibility of the genes. The figure below is zoomed into the circular section from the plot above.

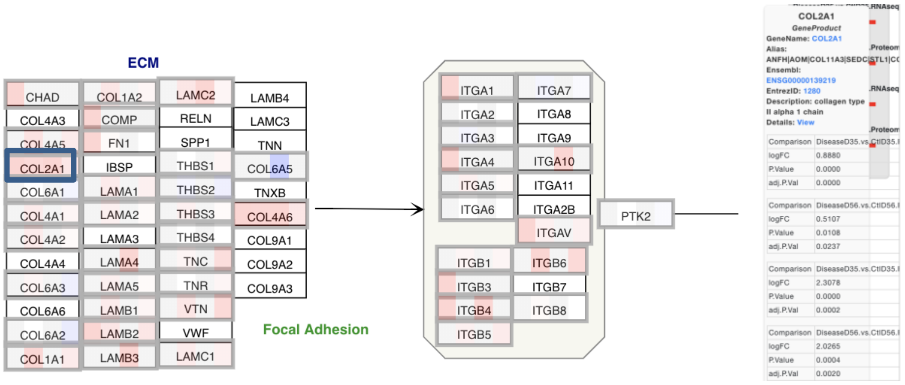

In this example, many of the ECM genes are getting differentially regulated in the same direction across all 4 comparisons. If a particular gene is clicked on, like COL2A1 here, the details of changes in the gene in each comparison can be seen.

### 4 Functional Enrichment from Comparisons

#### 4.1 Enrichment from Up and Down Regulated Genes

Let's first select a comparison to view.

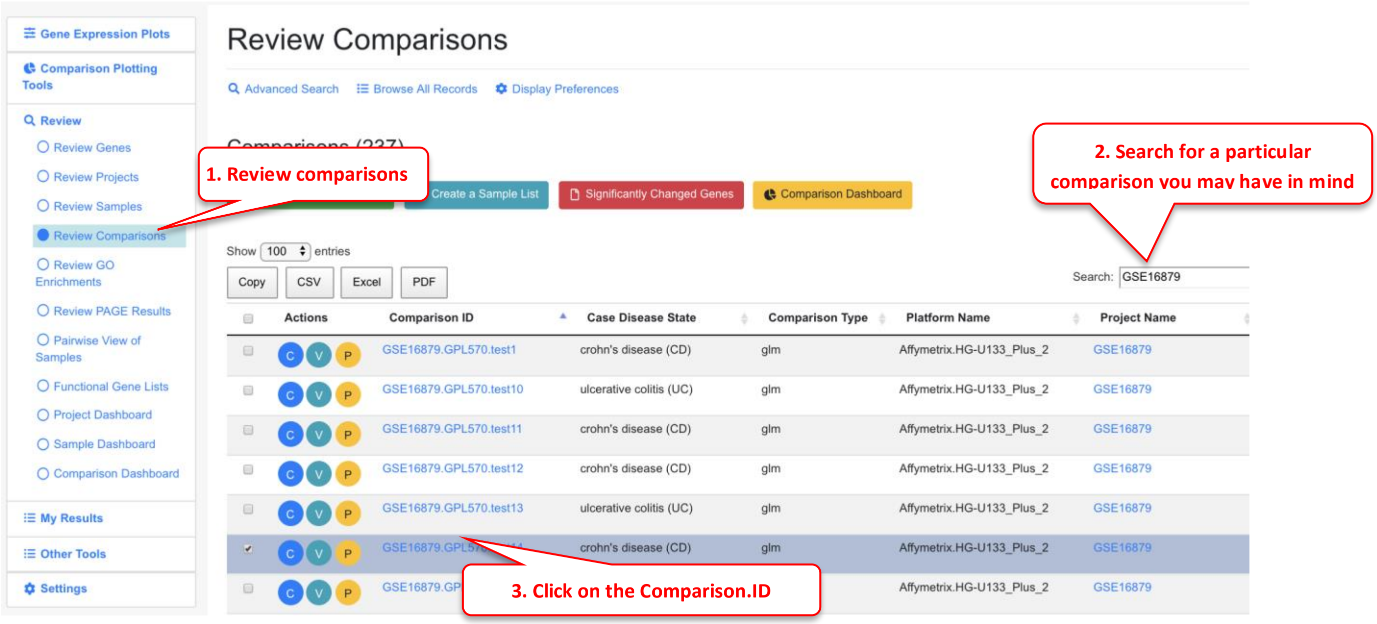

When Users view details of a comparison, the functional enrichment results are shown. Briefly, for each comparison, we generated the up- and down- regulate gene lists and use these lists to compare with all genes in the genome to identify functions that are significantly enriched.

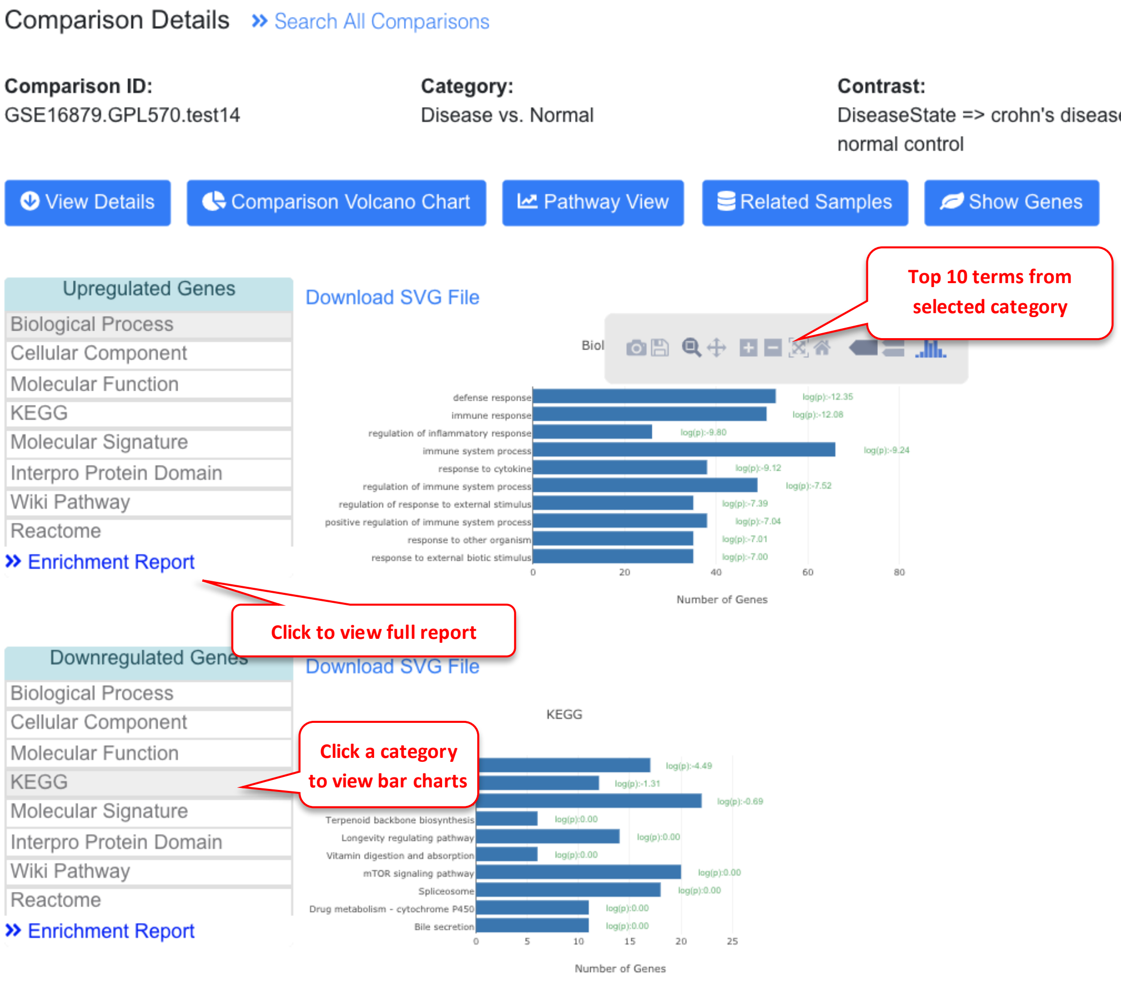

In the example above, this comparison is between Crohn's disease (CD) vs normal control, and the top up-regulated biological processes are defense and immune response.

Click the left menu will switch the bar charts for different categories (Gene Ontology, KEGG, Molecular signature, Protein domain etc).

The bar charts here show the top 10 categories. To view complete results, click the Enrichment Report.

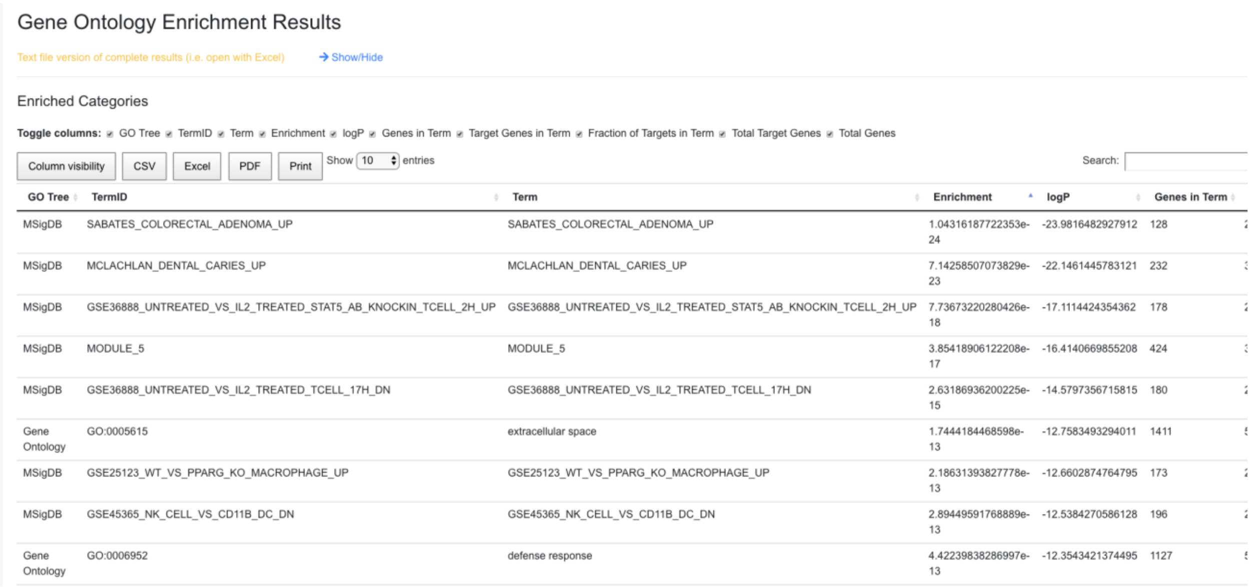

Gene Ontology Enrichment Results

Text file version of complete results (i.e. open with Excel) → Show/Hide

Enriched Categories

**Toggle columns:** GO Tree TermID Term Enrichment logP Genes in Term Target Genes in Term Fraction of Targets in Term Total Target Genes Total Genes

Column visibility | CSV | Excel | PDF | Print | Show 10 entries | Search:

| GO Tree | TermID | Term | Enrichment | logP | Genes in Term |
|---|---|---|---|---|---|
| MSigDB | SABATES_COLORECTAL_ADENOMA_UP | SABATES_COLORECTAL_ADENOMA_UP | 1.04316187722353e-24 | -23.9816482927912 | 128 |
| MSigDB | MCLACHLAN_DENTAL_CARIES_UP | MCLACHLAN_DENTAL_CARIES_UP | 7.14258507073829e-23 | -22.1461445783121 | 232 |
| MSigDB | GSE36888_UNTREATED_VS_IL2_TREATED_STAT5_AB_KNOCKIN_TCELL_2H_UP | GSE36888_UNTREATED_VS_IL2_TREATED_STAT5_AB_KNOCKIN_TCELL_2H_UP | 7.73673220280426e-18 | -17.1114424354362 | 178 |
| MSigDB | MODULE_5 | MODULE_5 | 3.85418906122208e-17 | -16.4140669855208 | 424 |
| MSigDB | GSE36888_UNTREATED_VS_IL2_TREATED_TCELL_17H_DN | GSE36888_UNTREATED_VS_IL2_TREATED_TCELL_17H_DN | 2.63186936200225e-15 | -14.5797356715815 | 180 |
| Gene Ontology | GO:0005615 | extracellular space | 1.7444184468598e-13 | -12.7583493294011 | 1411 |
| MSigDB | GSE25123_WT_VS_PPARG_KO_MACROPHAGE_UP | GSE25123_WT_VS_PPARG_KO_MACROPHAGE_UP | 2.18631393827778e-13 | -12.6602874764795 | 173 |
| MSigDB | GSE45365_NK_CELL_VS_CD11B_DC_DN | GSE45365_NK_CELL_VS_CD11B_DC_DN | 2.89449591768889e-13 | -12.5384270586128 | 196 |
| Gene Ontology | GO:0006952 | defense response | 4.42239838286997e-13 | -12.3543421374495 | 1127 |

In the enrichment report, the full list of functional terms are shown by order of Enrichment.

#### 4.2 View Changed Genes from a Functional Term in Volcano Plot

From the bar chat, click a functional term, and you have the option to view these genes in a volcano plot.

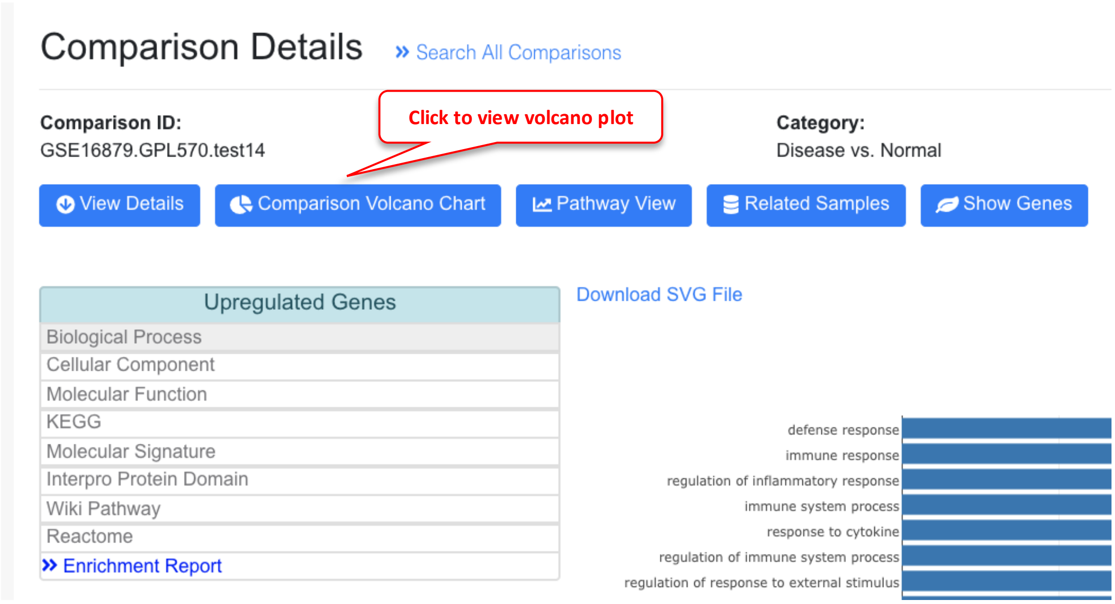

Once you click the link, volcano plot will be generated for the comparison with the changed genes from the selected term highlighted.

Comparison of interest

### Volcano Plot » View Genes

Comparison ID: GSE16879.GPL570.test14 Select Comparison

Please enter the comparison id, e.g., GSE44720.GPL10558.test16

Y-axis Statistics: ○ P-value ◉ FDR

Chart Name: Volcano Chart

Fold Change Cutoff: 2

Statistic Cutoff: 0.05

Show Gene Name: ○ Auto (based on cutoff) ◉ Customize

Changed genes from the functional term

Enter Genes Names: Load Saved Genes

S100A8
MNDA
FCGR3A
FCGR3B
SELL
SELE
PTGS2

» Add A New Chart

Plot

#### Summary

| | |
|---|---|
| Comparison: | GSE16879.GPL570.test14 |
| Fold Change Cutoff: | 2 |
| $Log_2$(Fold Change Cutoff): | 1.000 |
| Stat Cutoff: | 0.05 |
| $-Log_{10}$(Stat Cutoff): | 1.301 |
| # of Upregulated Genes: | 87 |
| # of Downregulated Genes: | 24 |
| All Genes: | 54,076 |

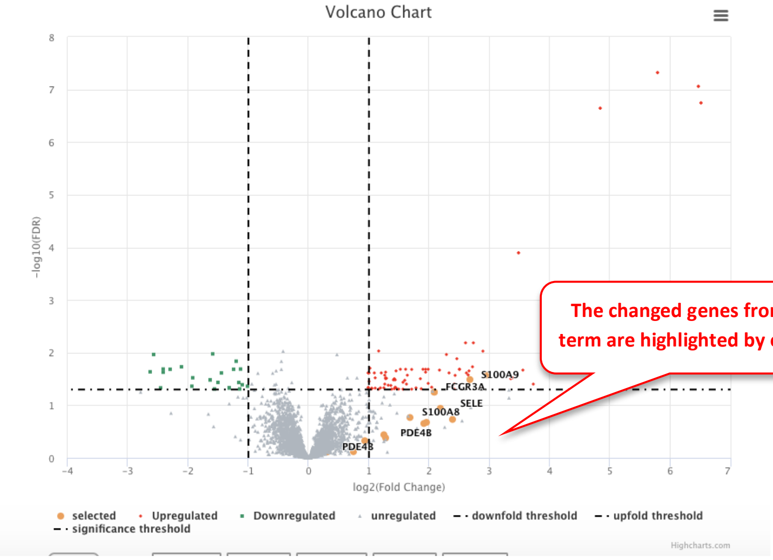

The changed genes from the term are highlighted by orange

Table of changed genes

Show 10 entries | Copy | CSV | Excel | PDF | Print | Search:

| Gene ID | Description | Log2FC | FDR | P-Value |
|---|---|---|---|---|
| FCGR3A | Fc fragment of IgG receptor IIIa | 2.0886 | 5.619e-002 | 5.234e-004 |
| FCGR3B | Fc fragment of IgG receptor IIIb | 2.0886 | 5.619e-002 | 5.234e-004 |
| FCGR3B | Fc fragment of IgG receptor IIIb | 2.971 | 2.677e-002 | 6.056e-005 |
| MNDA | myeloid cell nuclear differentiation antigen | 1.9234 | 2.219e-001 | 2.188e-002 |
| [illegible] | [illegible] | [illegible] | [illegible] | [illegible] |

#### 4.3 View Enriched Pathways Directly from Comparison Details

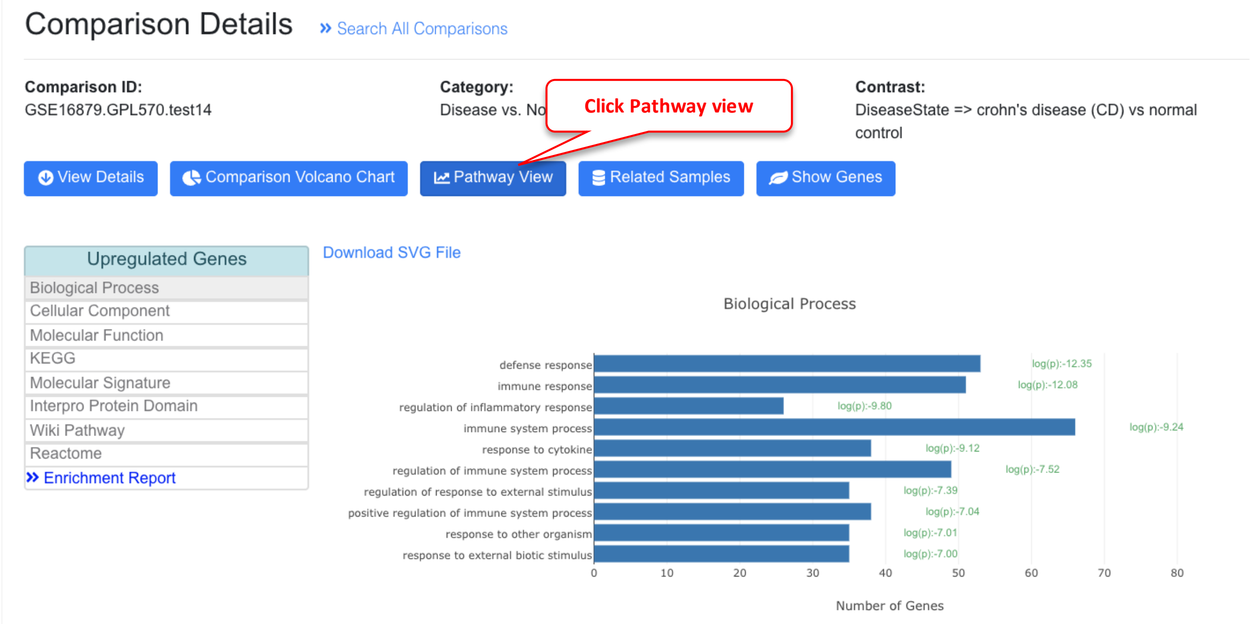

This will automatically open the pathway visualization page, and preload the pathway and comparison.

#### 4.4 Gene Set Enrichment from Ranked Genes

For each comparison, we produce a rank file for all genes using logFC. We use PAGE (Parametric Analysis of Gene Set Enrichment) to identify significant biological changes. PAGE can be more sensitive for comparisons where the logFC is relatively small, but most genes in a functional set show the same direction of change.

The predefined gene sets were from MSigDB.

For each comparison, the top up-regulated and down-regulated gene sets are plotted.

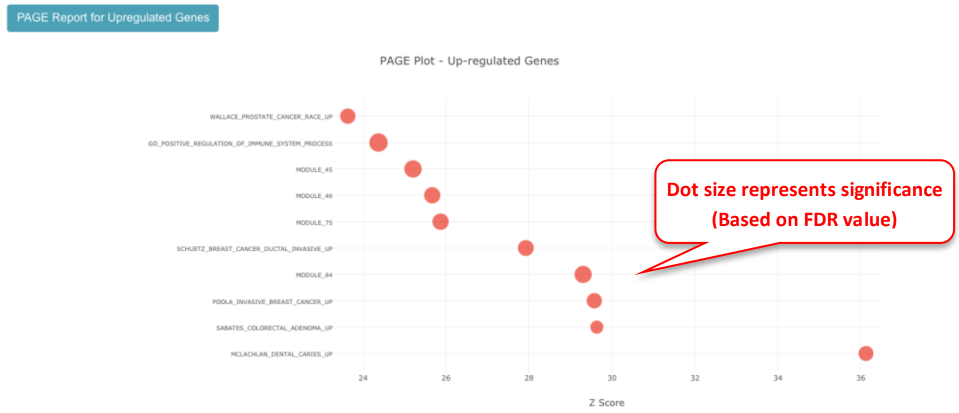

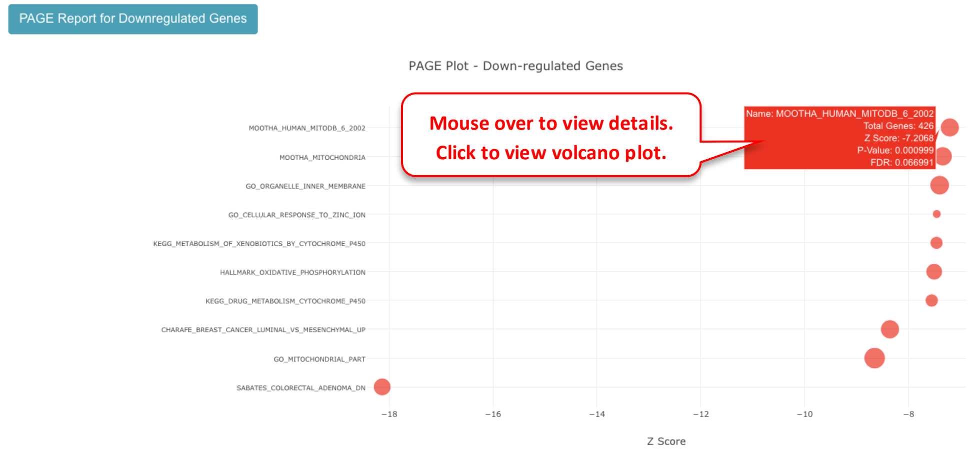

To view the genes within a gene set using volcano plot, you can click the dot, and use the link from the pop-up.

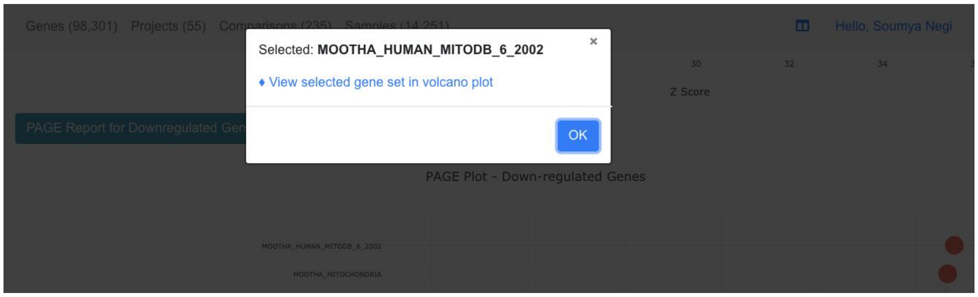

#### 4.5 Pathway Heatmap From Comparisons

Users can display the enriched pathways from several related comparisons, and visualize the top enriched pathways across comparisons. Users can mix public data and inhouse comparisons.

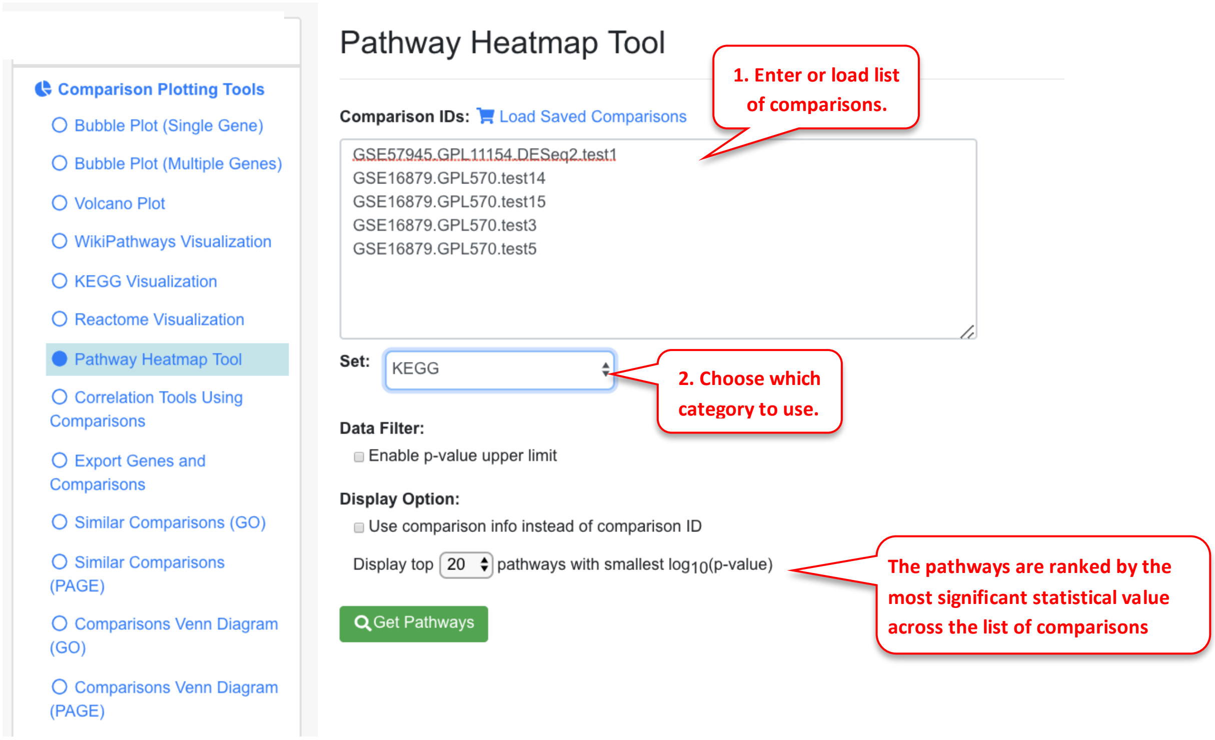

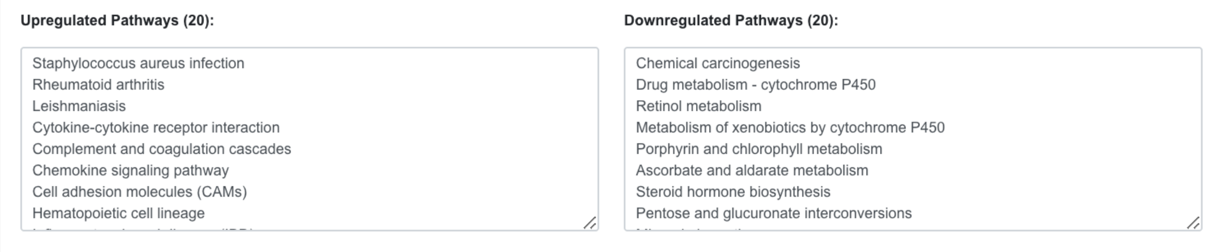

The heatmap shows pathways in rows, comparisons in columns. The statistical significance is color-coded (log P-value, or Z-score). Pathways are sorted by the negative logP values from the highest to the lowest.

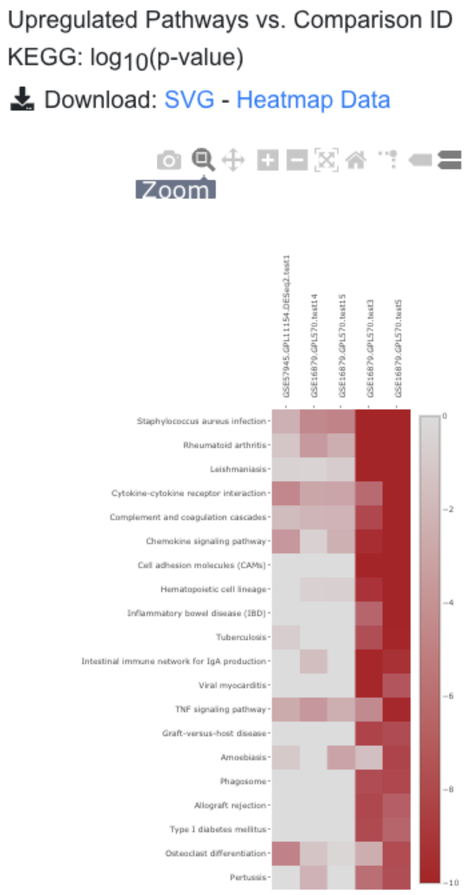

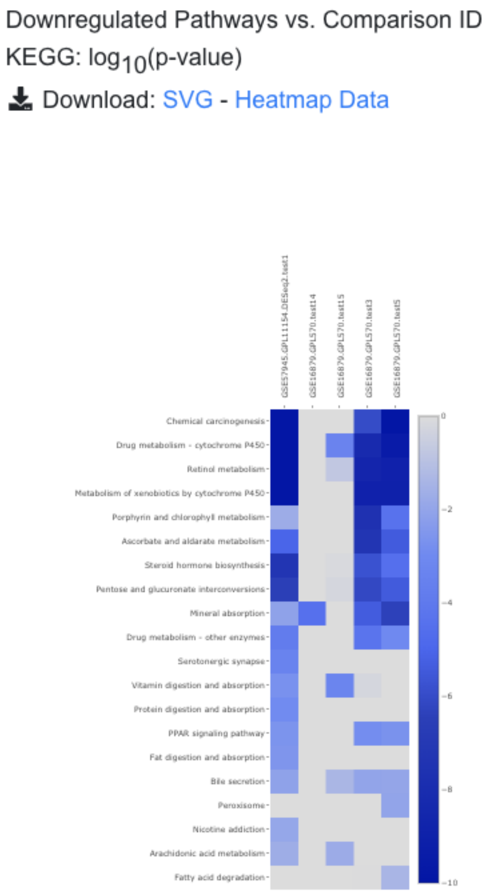

From the pathway heatmap, users can click any data point to view details.

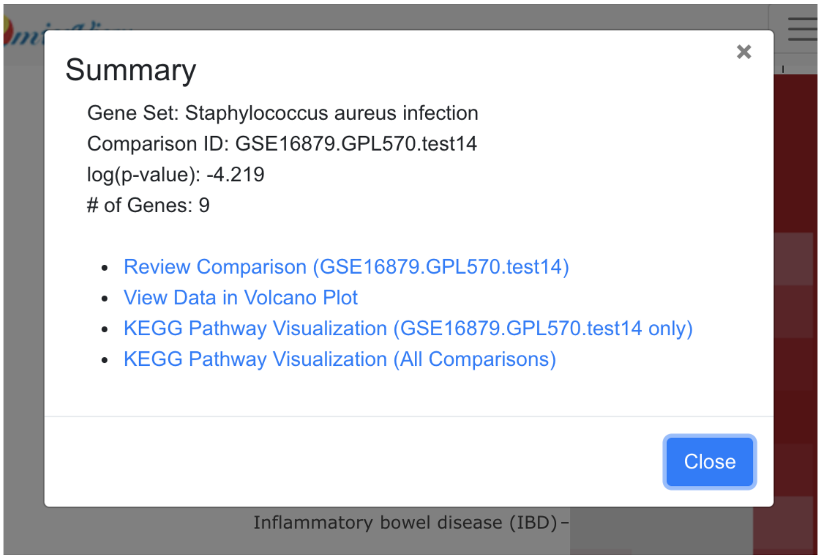

### 5 Data Upload

OmicsView gives users the option to add their own data, or published data that is not on the portal yet. This is achieved through the “Internal Data” option under “My Results”.

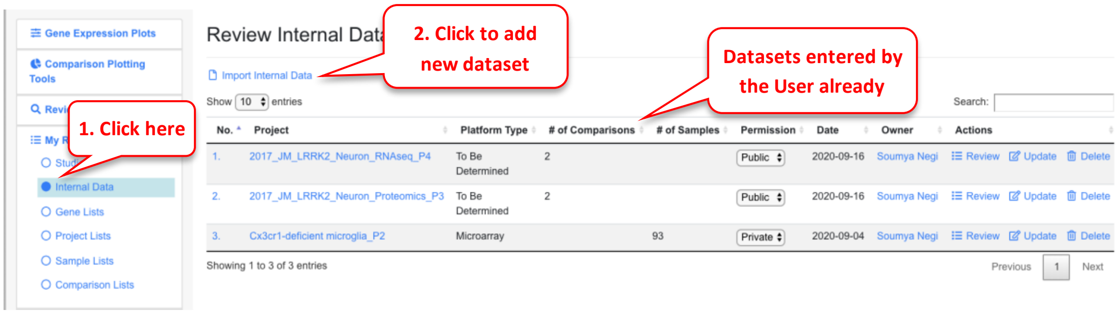

To add a new dataset, click on the link “Import Internal Data” that will lead you to a page like below.

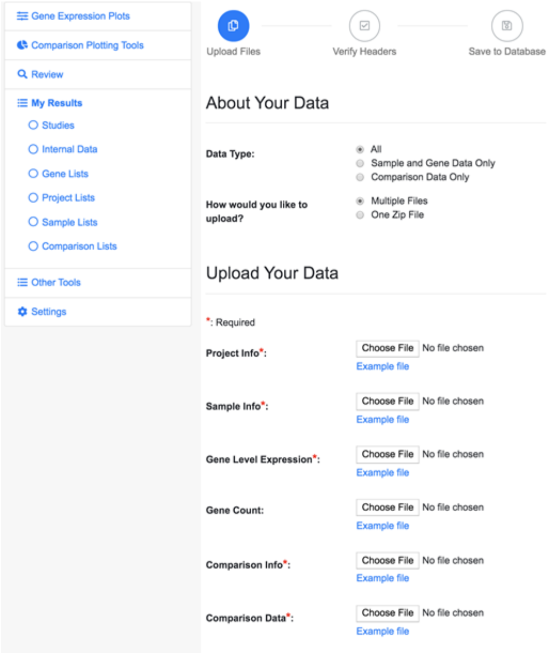

#### 5.1 Data Type

There are three types of project data. The file requirements for each data type are summarized in the table below.

| Data Type | All | Sample and Gene Data Only | Comparison Data Only |
|---|---|---|---|
| Files Required | Project Info file<br>Sample Info file<br>Gene Level Expression file<br>Comparison Info file<br>Comparison Data file<br>Gene Count file (Optional) | Project Info file<br>Sample Info file<br>Gene Level Expression file<br>Gene Count file (Optional) | Project Info file<br>Comparison Info file<br>Comparison Data file |

When you select different Data Types, the **Upload Your Data** session in the page changes accordingly.

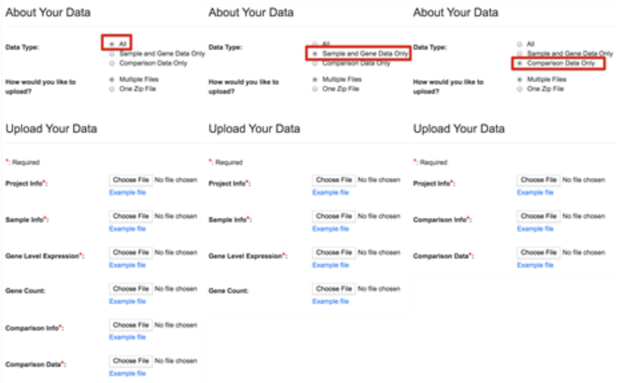

##### 5.1.1 Supported Data File Format

The system support two data formats:

- The csv format (comma separated)
- The tab delimited format

Be sure that your source data files are in the supported format. Usually **Auto Detect** is selected as default and doesn't need to change.

##### 5.1.2 Expression Data Format

The Gene Level Expression file can have two formats: Matrix format and Table format. Click on the **Help** link below the selection box to see examples.

The application supports two different formats:

Matrix Format

This format supports only one kind of data at a time.

| Gene | Sample_1 | Sample_2 | Sample_3 |
|---|---|---|---|
| CREB1 | 6.08 | 4.92 | 9.56 |
| TP53 | 6.27 | 7.55 | 6.68 |
| WASH7P | 3.08 | 3.17 | 8.76 |

Table Format:

This format supports only multiple kinds of data.

| Gene | Sample_ID | Expression | Count |
|---|---|---|---|
| CREB1 | Sample_1 | 4.55 | 89 |
| CREB1 | Sample_2 | 2.41 | 18 |
| CREB1 | Sample_3 | 3.53 | 76 |
| TP53 | Sample_1 | 2.87 | 71 |
| TP53 | Sample_2 | 8.72 | 64 |
| TP53 | Sample_3 | 7.29 | 37 |
| WASH7P | Sample_1 | 7.51 | 31 |
| WASH7P | Sample_2 | 9.79 | 30 |
| WASH7P | Sample_3 | 5.71 | 39 |

The **Study** option is useful if you want to include the uploaded project in a specific study. And the **Access** option can be checked if you want to make your project viewable by all the users.

#### 5.2 Data Upload options

Instead of uploading the source data file individually, you can compress your data files into a zip file and upload only this zip file. The application will determine the data type based on the file name. Make sure that your zip file contains the following files with exactly the same file names. You can check the requirement of the zipped file for each data type by clicking on the Requirement link under the Choose File box in the **Upload Your Data** session.

Upload Your Data

*: Required

Zip_File*: Choose File No file chosen
Requirement

This table below summarizes the file requirement for all data types.

| Data Type | File requirement |
|---|---|
| Data Type: ◉ All ○ Sample and Gene Data Only ○ Comparison Data Only<br>How would you like to upload? ○ Multiple Files ◉ One Zip File | *: Required<br>• **Project Info***: Project_Info.csv<br>• **Sample Info***: Sample_Info.csv<br>• **Gene Level Expression***: Gene_Expression_Data.csv<br>• Gene Count: Gene_Count.csv<br>• **Comparison Info***: Comparison_Info.csv<br>• **Comparison Data***: Comparisons_Data.csv |
| Data Type: ○ All ◉ Sample and Gene Data Only ○ Comparison Data Only<br>How would you like to upload? ○ Multiple Files ◉ One Zip File | *: Required<br>• **Project Info***: Project_Info.csv<br>• **Sample Info***: Sample_Info.csv<br>• **Gene Level Expression***: Gene_Expression_Data.csv<br>• Gene Count: Gene_Count.csv |
| Data Type: ○ All ○ Sample and Gene Data Only ◉ Comparison Data Only<br>How would you like to upload? ○ Multiple Files ◉ One Zip File | *: Required<br>• **Project Info***: Project_Info.csv<br>• **Comparison Info***: Comparison_Info.csv<br>• **Comparison Data***: Comparisons_Data.csv |

#### 5.3 Source File format

You can always check the source file format by looking at the example files. They are located below each file selection box.

Upload Your Data

*: Required

Project Info*: Choose File No file chosen
Example file

Sample Info*: Choose File No file chosen
Example file

Gene Level Expression*: Choose File No file chosen
Example file

Gene Count: Choose File No file chosen
Example file

Comparison Info*: Choose File No file chosen
Example file

Comparison Data*: Choose File No file chosen
Example file

##### 5.3.1 Project Info file

| ProjectID | Description | Disease | Platform | Accession |
|---|---|---|---|---|
| GSE76161 | Background: The bile acid-activ | nonalcoholic steatohepatitis (NASH) | GPL11532 | GSE76161 |
| GSE57227 | We report the genome-wide p | nonalcoholic steatohepatitis (NASH) | GPL11154 | GSE57227 |
| GSE67492 | Gene expression in the right ve | pulmonary arterial hypertension (PAH) | HuGene-1_0-st-v1 | GSE67492 |

- Only ProjectID is required and must be unique across the system.
- The other four fields are highly recommended.
- The user can add one or more projects.

Save the file in the supported format and upload to the system. The system has a tool to check all the available fields and existing values for the project info file. Click on **Distinct Values in Table** under the **Other Tools** in the left menu.

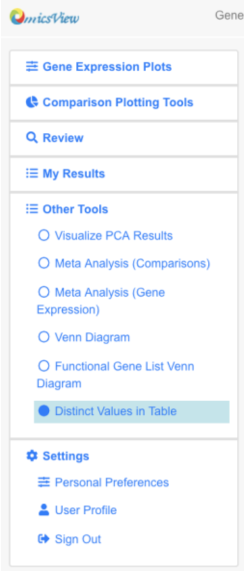

And you will see a searching tool.

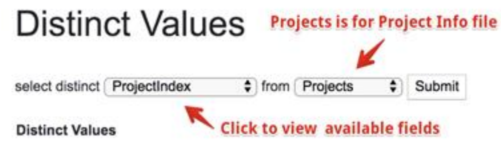

Select your interested fields to check the existing values for that field. Try your best to use the same value for the same info, which can ensure your imported projects are comparable when doing cross project analysis.

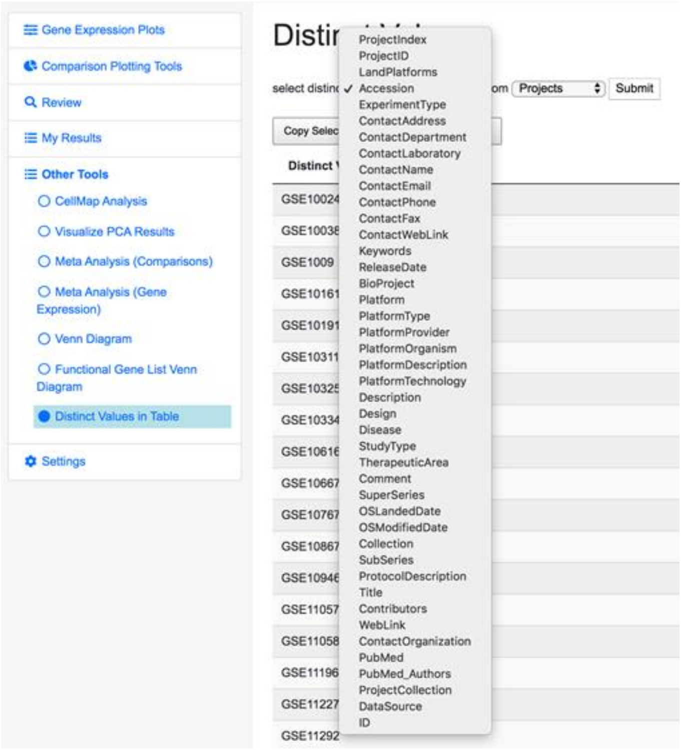

##### 5.3.2 Sample Info file

**Recommended fields**

| **SampleID** | **ProjectID** | **PlatformName** | Description | Tissue | DiseaseState | SampleSource | Gender |
|---|---|---|---|---|---|---|---|
| E-MTAB-2325_10 | E-MTAB-2325 | Agilent.Expression.Human4x44k | No Info | neocortex | normal control | neocortex | male |
| E-MTAB-2325_11 | E-MTAB-2325 | Agilent.Expression.Human4x44k | No Info | neocortex | amyotrophic lateral sclerosis | neocortex | male |
| E-MTAB-2325_12 | E-MTAB-2325 | Agilent.Expression.Human4x44k | No Info | neocortex | amyotrophic lateral sclerosis | neocortex | female |

- The first three fields (in bold) are required.
- SampleID should be unique among the same project.
- The projectID field should match with the projectID field in the Project Info file.
- The platformName should be one from the available values in the Distinct Values table.
- The other fields listed here are not required but highly recommended. These are used to capture information provided by researchers about the samples.
- The headers must match one of the distinct fields in the Distinct Values tool. If the column header provided by the user does not match any available field, the system will show the unmatched fields in the file and let you manually map it to a known field.

To check the available fields and existing values, you can also check the **Distinct Values** tool by selecting **Samples** in the box.

##### 5.3.3 Comparison Info file

**Recommended fields**

| **ComparisonID** | **PlatformName** | **ProjectID** | Case.SampleIDs | Control.SampleIDs | ComparisonCategory | ComparisonContrast | Case.DiseaseState | Case.Tissue |
|---|---|---|---|---|---|---|---|---|
| E-MTAB-4304.GPL11 | NGS.Illumina.HiSeq | E-MTAB-4304 | E-MTAB-4304_0;E-MTA | E-MTAB-4304_1;E-MTA | Tissue1 vs. Tissue2 | TissueRegion => distal medial condyl | knee osteoarthritis | knee cartilage |
| E-MTAB-4586.GPL11 | NGS.Illumina.HiSeq | E-MTAB-4586 | E-MTAB-4586_0;E-MTA | E-MTAB-4586_14;E-MT | Disease vs. Normal | DiseaseState => Parkinson's disease ( | Parkinson's disease | skin |
| E-MTAB-4586.GPL11 | NGS.Illumina.HiSeq | E-MTAB-4586 | E-MTAB-4586_0;E-MTA | E-MTAB-4586_14;E-MT | Other Comparisons | GeneticModification => LRRK2 G2019 | Parkinson's disease | skin |

- The first three fields (in bold) are required.
- ComparisonID should be unique among the same project.
- The projectID field should match with the projectID field in the Project Info file.

- The platformName should be one from the available values for the platformName in the Distinct Values table.
- The next two fields Case.SampleIDs and Control.SampleIDs should be used to list the sampleIDs used as cases or controls for the comparison. Separate sampleIDs by comma.
- The other fields listed here are not required but highly recommended. For example, ComparisonCategory is used in group comparisons in dashboard. Case.DiseaseState and Case.Tissue are used in coloring or grouping bubble plots.
- The headers must match one of the distinct fields in the Distinct Values tool. If the column header provided by the user does not match any available field, the system will show the unmatched fields in the file and let you manually map it to a known field.

#### 5.4 Data Formats

There are certain formats for gene expression data and comparison data files. The data format below applies to both RNA-Seq and Array data.

##### 5.4.1 Gene Expression Data

For gene expression data (RNA-Seq or microarray), two types of formats are supported.

**Matrix format**

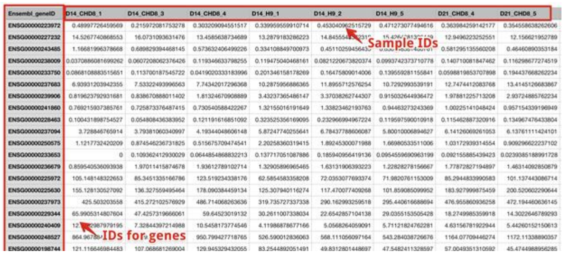

**Table format**

| Gene | SampleID | Expression |
|---|---|---|
| ARRB2 | lesional_trt_2 | 16.0249 |
| ARRB2 | non_lesional_none_2 | 10.1214 |
| ARRB2 | non_lesional_trt_2 | 8.4008 |
| ARRB2 | lesional_none_3 | 25.3016 |
| ARRB2 | lesional_trt_3 | 16.5416 |
| ARRB2 | non_lesional_none_3 | 9.1222 |
| ARRB2 | non_lesional_trt_3 | 9.501 |
| ARRB2 | lesional_none_4 | 20.2206 |
| ARRB2 | lesional_trt_4 | 21.5801 |
| ARRB2 | non_lesional_none_4 | 14.3564 |
| ARRB2 | non_lesional_trt_4 | 8.947 |
| ARRB2 | lesional_none_5 | 12.1129 |
| ARRB2 | lesional_trt_5 | 9.8666 |
| ARRB2 | non_lesional_none_5 | 6.7286 |

The **Gene** must be listed in the first column. The IDs can be official gene symbols, gene IDs (Entrez Id), or Ensmbl IDs. The system will automatically recognize the IDs and map the IDs to the gene annotation table. The SampleIDs must match the sampleIDs in the Sample Info file.

##### 5.4.2 Comparison Data

For comparison data, the following template should be used.

| Gene | ComparisonID | logFC | P.Value | adj.P.Val |
|---|---|---|---|---|
| ENSG00000223972 | D14_CHD8.vs.D14_H9 | -0.480181603380462 | 0.201172869981358 | 0.276823114956239 |
| ENSG00000227232 | D14_CHD8.vs.D14_H9 | -0.0260065294185647 | 0.782508711249897 | 0.834365881843091 |
| ENSG00000243485 | D14_CHD8.vs.D14_H9 | 0.614759995164825 | 0.12397256363312 | 0.183494049719199 |
| ENSG00000238009 | D14_CHD8.vs.D14_H9 | -0.489128093633942 | 0.325707592057561 | 0.414162918866014 |
| ENSG00000233750 | D14_CHD8.vs.D14_H9 | -1.10014257817111 | 0.0192044755497128 | 0.0369617315251983 |
| ENSG00000237683 | D14_CHD8.vs.D14_H9 | -0.605292542573319 | 3.41124756968239E-06 | 2.19219164320563E-05 |
| ENSG00000239906 | D14_CHD8.vs.D14_H9 | -1.0010125449076 | 0.0636863585580712 | 0.103931911218667 |
| ENSG00000241860 | D14_CHD8.vs.D14_H9 | -0.714487254252856 | 0.000262496694401327 | 0.000917136064345589 |
| ENSG00000228463 | D14_CHD8.vs.D14_H9 | -1.18951263566867 | 0.0140347276118906 | 0.0282154235613193 |
| ENSG00000237094 | D14_CHD8.vs.D14_H9 | -0.691939730736816 | 3.54212236297494E-07 | 3.17551702158323E-06 |
| ENSG00000250575 | D14_CHD8.vs.D14_H9 | -1.28179600460206 | 0.000183559753891093 | 0.000670021987202936 |
| ENSG00000233653 | D14_CHD8.vs.D14_H9 | -1.56406411915058 | 0.0289389868237879 | 0.0526241552771901 |
| ENSG00000236679 | D14_CHD8.vs.D14_H9 | 0.0814532190594377 | 0.835473327403157 | 0.877408636003058 |
| ENSG00000225972 | D14_CHD8.vs.D14_H9 | 0.545641008555407 | 0.000139660955234846 | 0.000526980960540253 |

List one or more comparisons in the table. The **Gene** must be listed in the first column. The IDs can be official gene symbols, gene IDs (Entrez Id), or Ensmbl gene IDs. The system will automatically recognize the IDs and map the IDs to the gene annotation table. The second column must be **ComparisonID**, which must match the comparisonIDs for the Comparison Info file. For each row, three values should be entered, **logFC** is the log2 Fold Change, **P.Value**, and **adj.P.Val** (FDR). Most statistical packages output these three values. If some values are missing, enter NAs.

### 6 Advanced Analyses

#### 6.1 Correlation Tools

Once the user has identified a gene of interest, the user can use correlation tools to find other genes that share similar (or opposite) profiles in terms of gene expression or fold change. First, enter the gene of interest, and samples to be used for correlation. In the example below, we entered OSM gene, and 69 samples involved in Crohn disease.

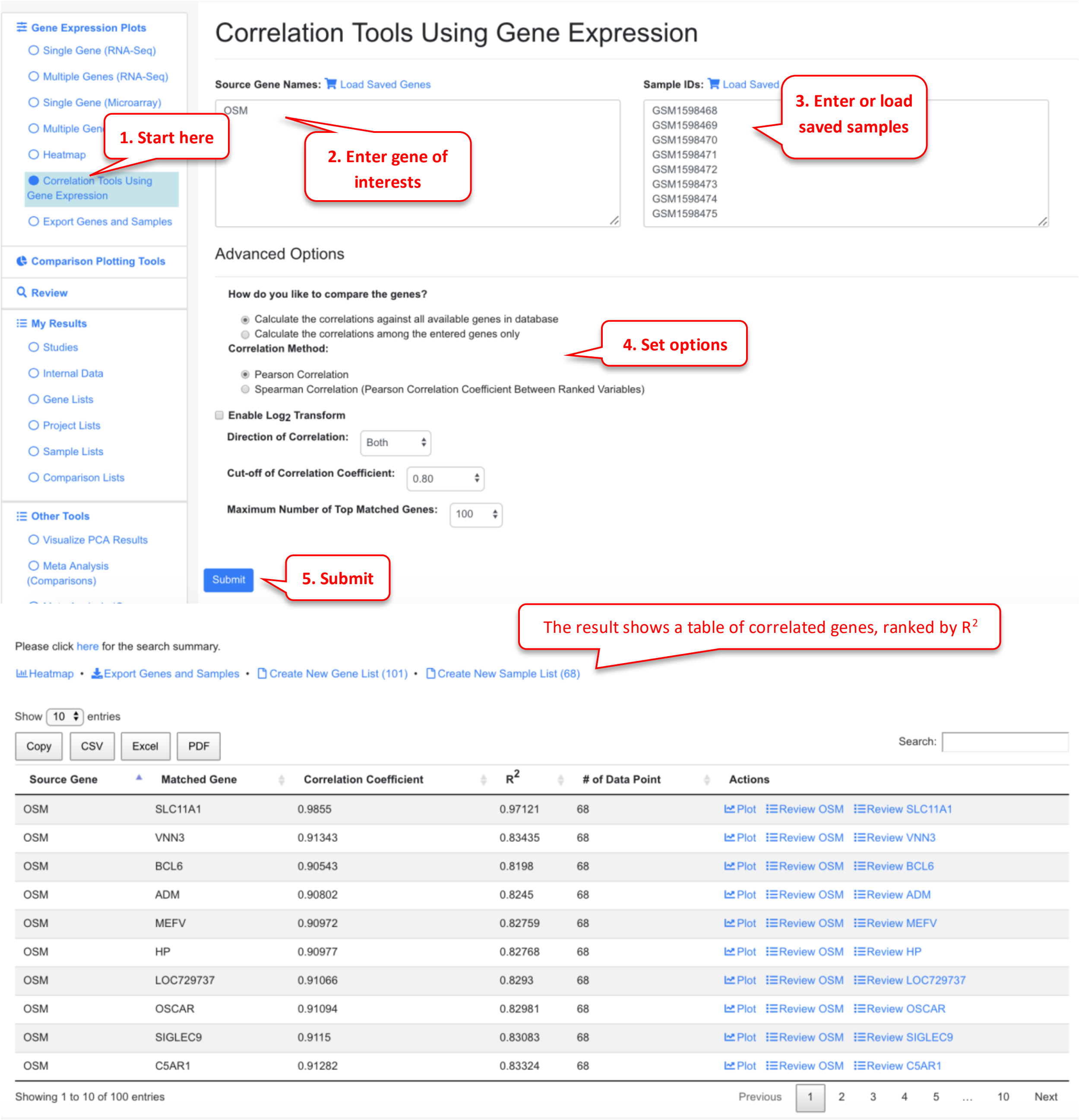

| Source Gene | Matched Gene | Correlation Coefficient | $R^2$ | # of Data Point | Actions |
|---|---|---|---|---|---|
| OSM | SLC11A1 | 0.9855 | 0.97121 | 68 | Plot Review OSM Review SLC11A1 |
| OSM | VNN3 | 0.91343 | 0.83435 | 68 | Plot Review OSM Review VNN3 |
| OSM | BCL6 | 0.90543 | 0.8198 | 68 | Plot Review OSM Review BCL6 |
| OSM | ADM | 0.90802 | 0.8245 | 68 | Plot Review OSM Review ADM |
| OSM | MEFV | 0.90972 | 0.82759 | 68 | Plot Review OSM Review MEFV |
| OSM | HP | 0.90977 | 0.82768 | 68 | Plot Review OSM Review HP |
| OSM | LOC729737 | 0.91066 | 0.8293 | 68 | Plot Review OSM Review LOC729737 |
| OSM | OSCAR | 0.91094 | 0.82981 | 68 | Plot Review OSM Review OSCAR |
| OSM | SIGLEC9 | 0.9115 | 0.83083 | 68 | Plot Review OSM Review SIGLEC9 |
| OSM | C5AR1 | 0.91282 | 0.83324 | 68 | Plot Review OSM Review C5AR1 |

Click the plot icon will show scatter plot of the target and the correlated gene.

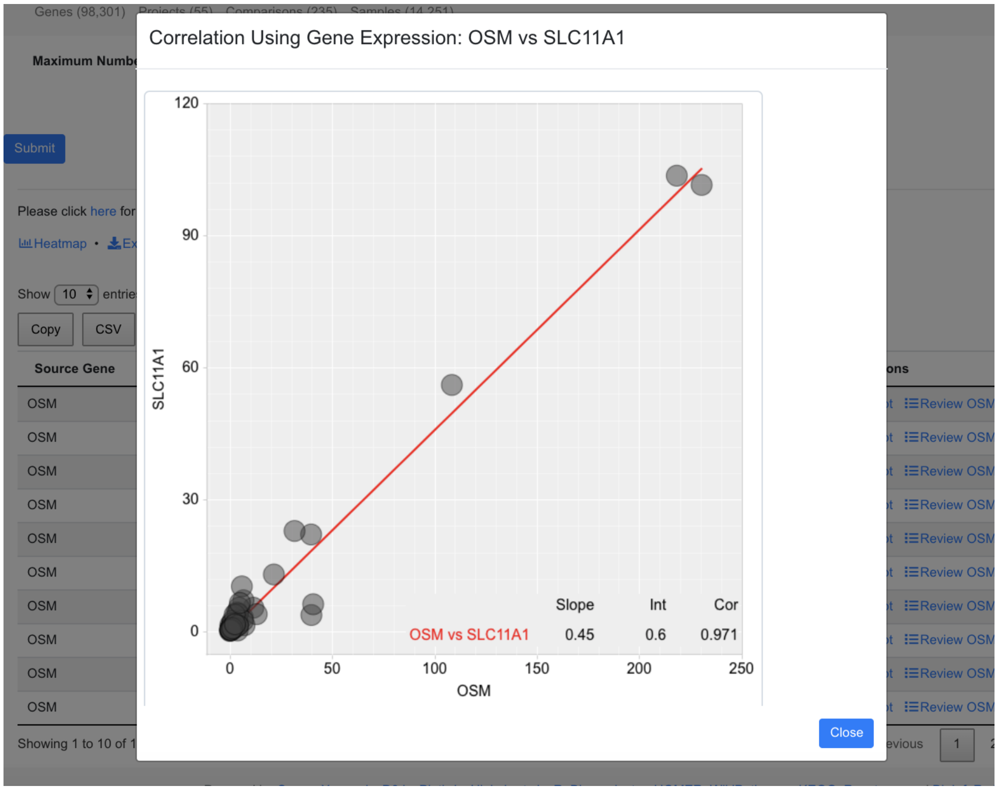

An additional way to plotting the correlation with other genes, is to enable the log2 transformation of the expression.

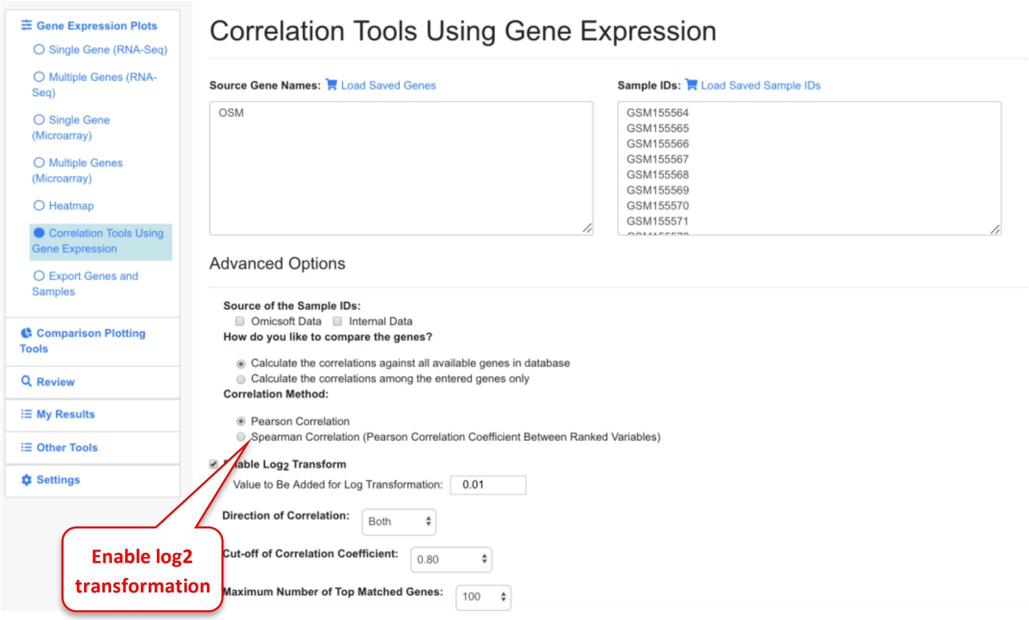

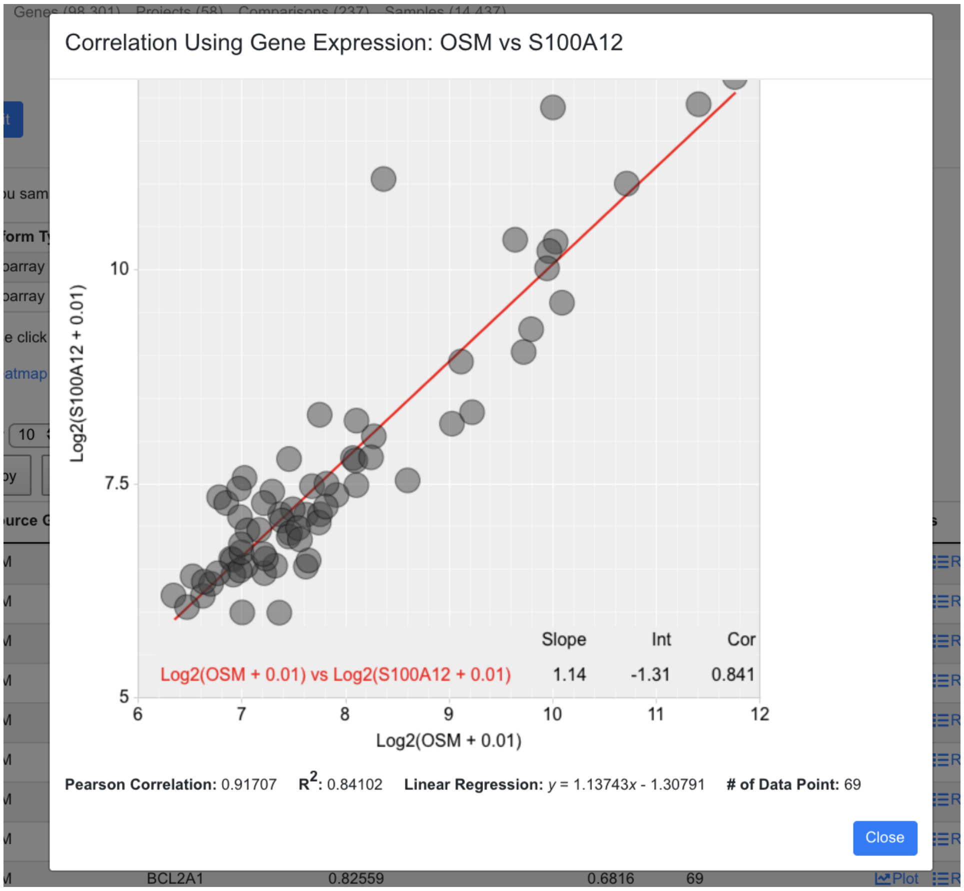

Now the expression is log transformed.

The examples above are from Correlation Tools using Expression. The interface and usage for Correlation Tools using Comparison is similar.

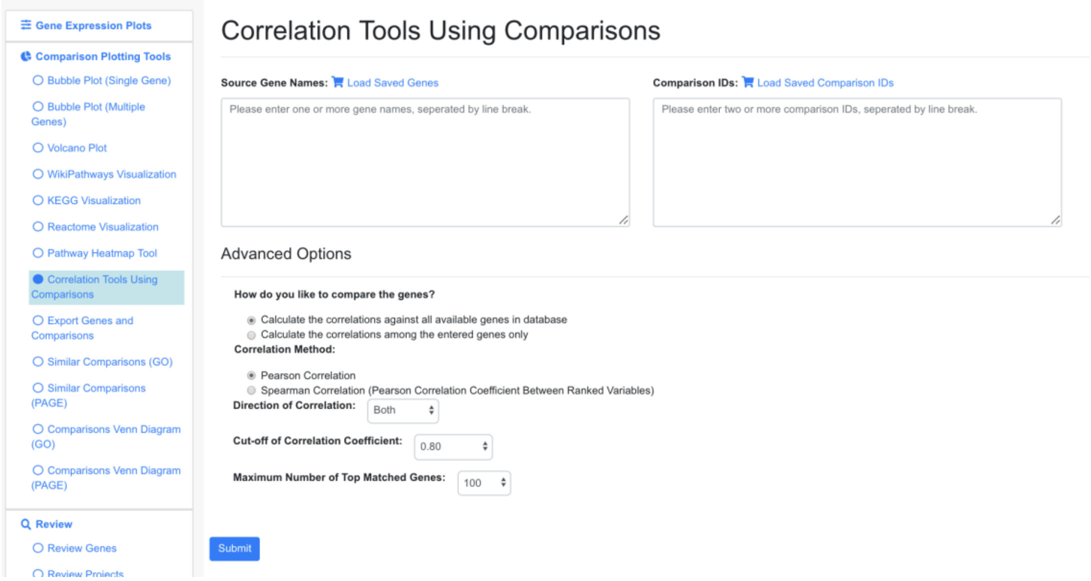

#### 6.2 PCA Analysis

Users can select a few samples and use PCA plot to visualize the sample relationships.

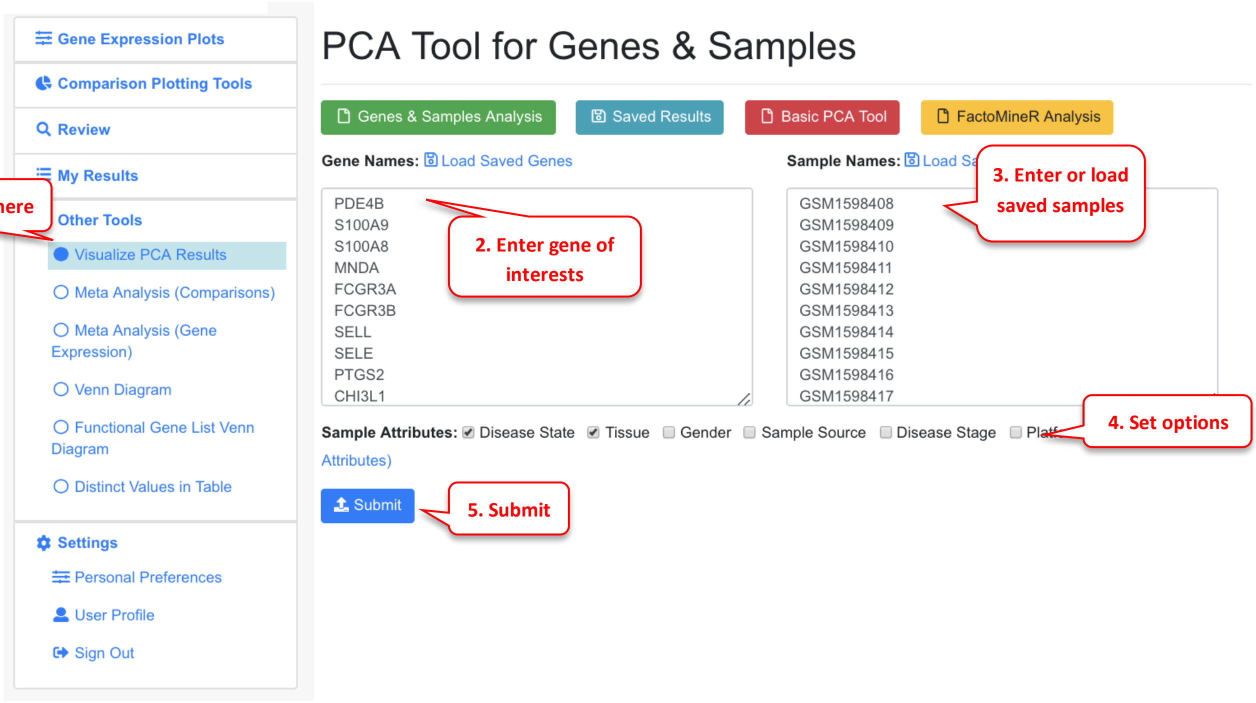

The system will use FactorMineR package to run PCA analysis and display the results.

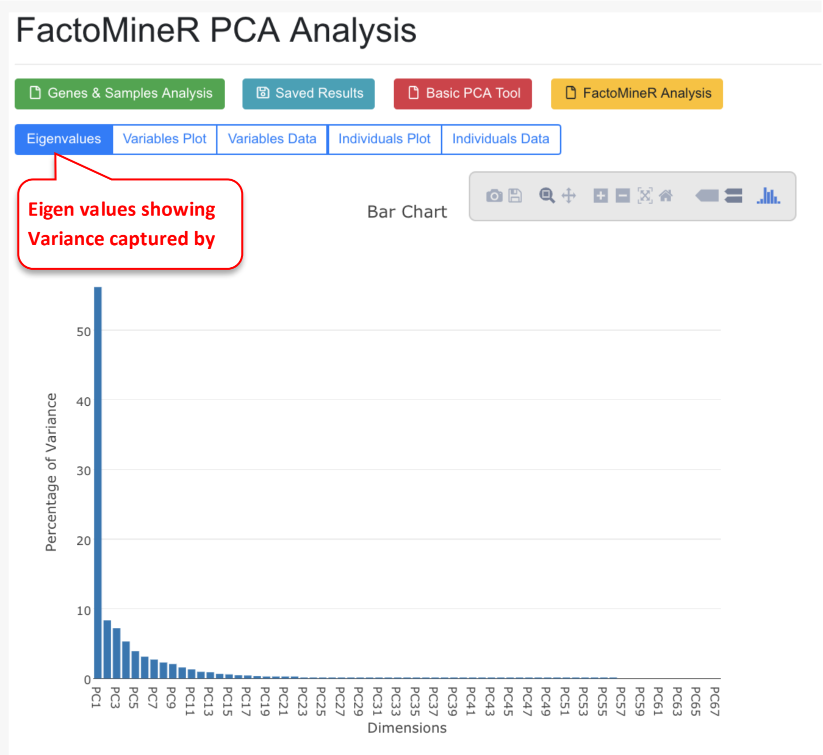

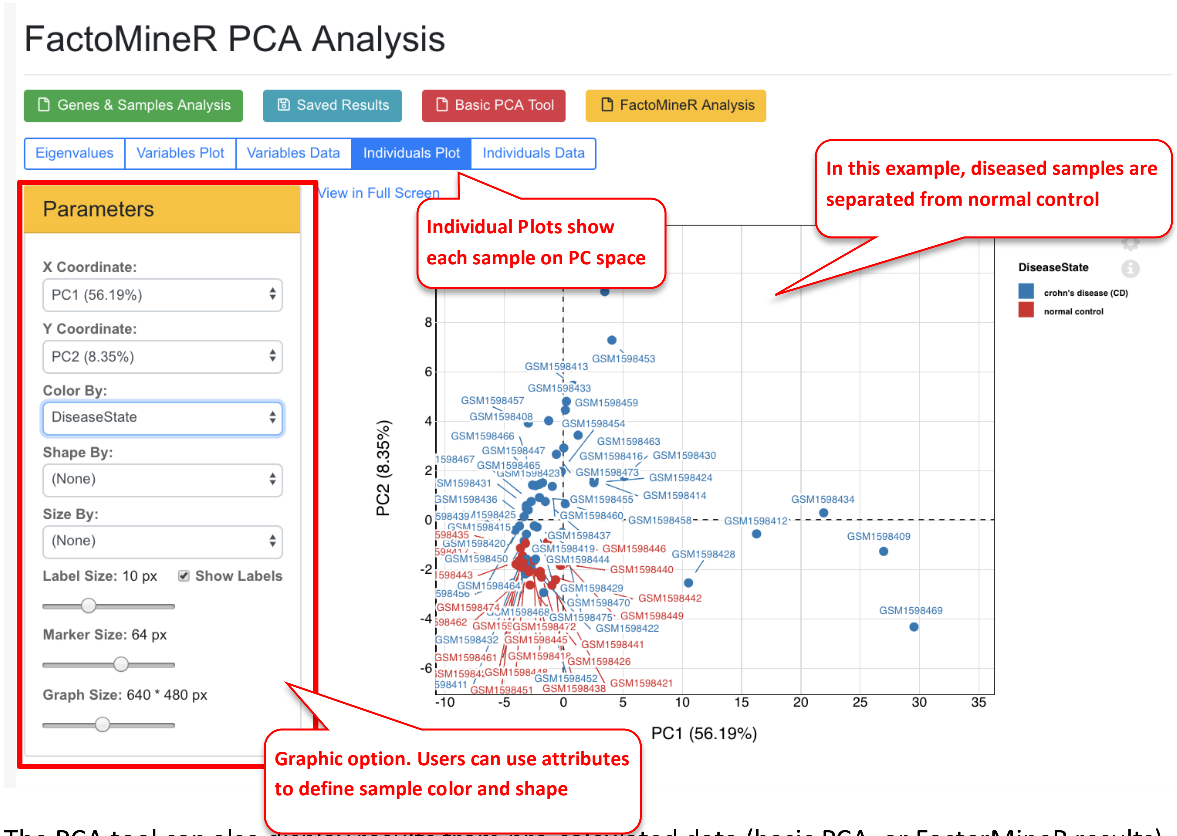

The PCA tool can also display results from pre-calculated data (basic PCA, or FactorMineR results).

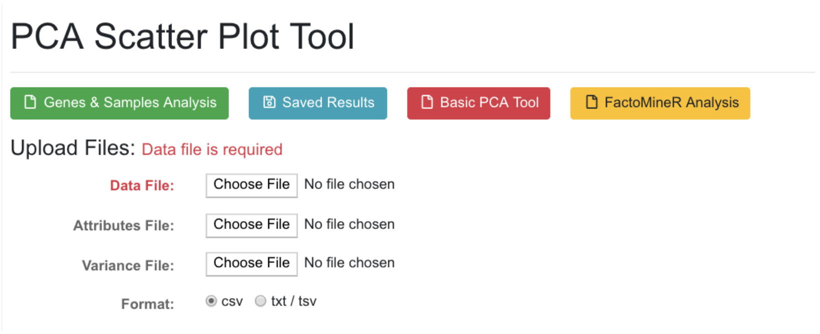

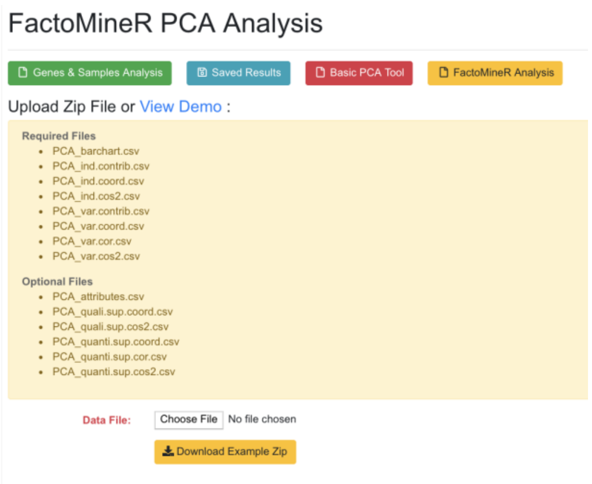

#### 6.3 Meta-Analysis

Meta-Analysis can be used to identify genes that are changed consistently across multiple projects. This can be either implemented by doing Meta-analysis using comparisons or meta-analysis using gene expression.

##### 6.3.1 Meta-Analysis (Comparisons)

In the example below, we are looking for the most significant DEGs in Disease vs. Normal comparisons of Crohns Disease.

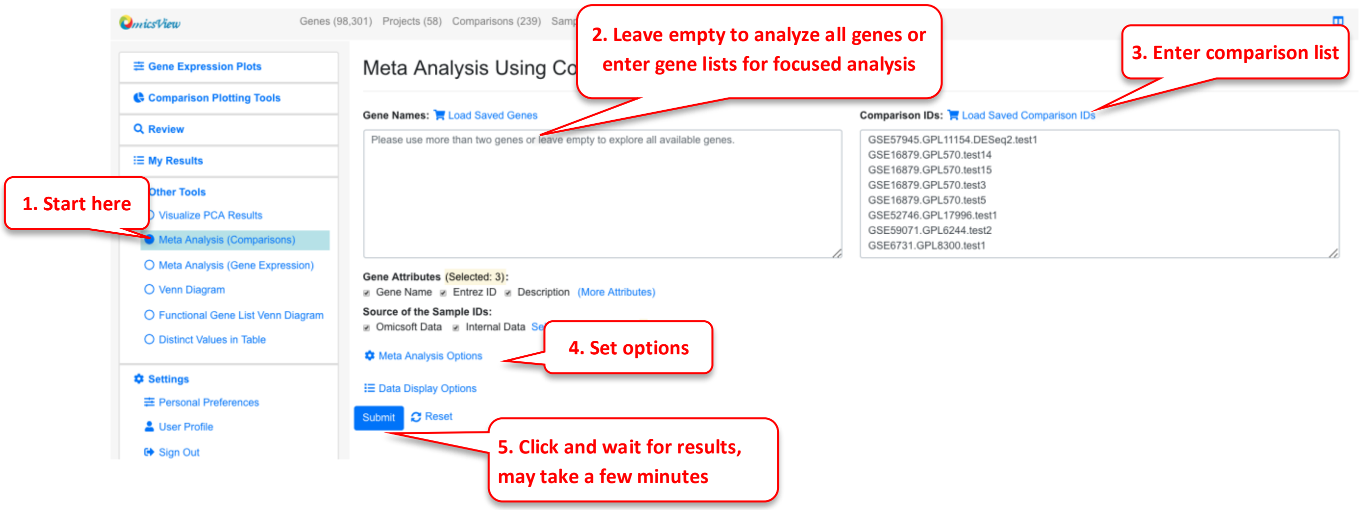

The system will use the comparison data (logFC, p-value) to compute combined p-value and rank product. It also uses a simple cutoff to get counts of up- and down- regulated genes. This method is fast and can be applied to any comparison data. However, it does not use the individual sample data or considers number of samples in each comparison.

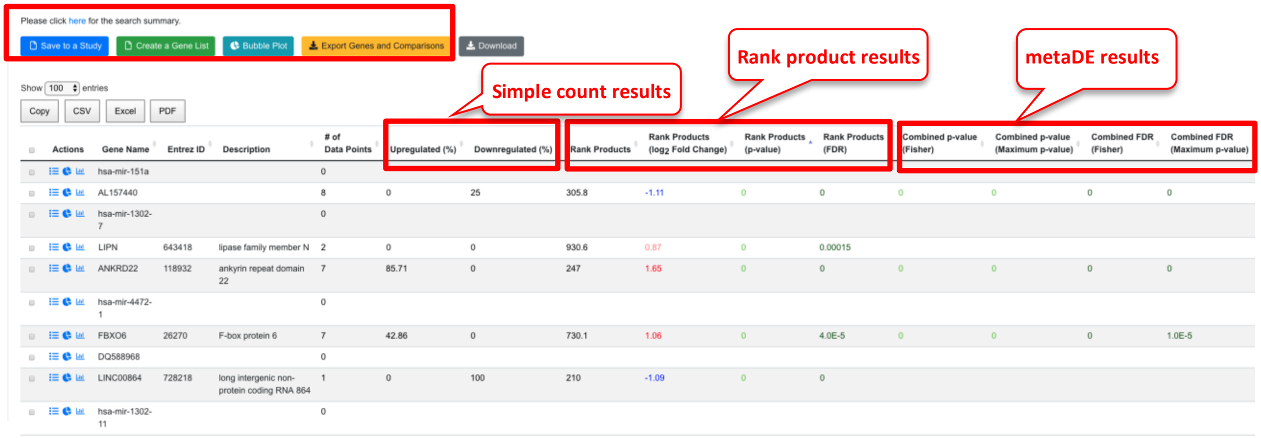

We can enable filter by percentage up-regulated genes to focus on genes that are consistently up-regulated.

From the output table, the user can select the genes most interesting to them. Some key columns are explained below.

- # of data points: this shows the number of comparisons with valid data for this gene. Although 8 comparisons were entered here, not all genes show up in all experiments. We used minimal 3 data points when we set up display options.
- Simple count results. Use the cutoff values we entered (logFC 1, FDR 0.05), the system checks if a gene pass the cutoff for each comparison, and computes percentage up- and down-regulation.
- Rank Products results (recommended results for meta-analysis). This is done with the RankProd package. The system rank genes in each comparison using logFC and computes a combined rank and statistical values. The Rank Products columns shows the rank (smaller is more consistent change), log2Fold Change and FDR.
- The Combined p-value/FDR (Fisher or Maximum) are computed using the individual p-values with metaDE package. Note unlike RankProd, these values didn't consider the direction of the change, so the data need to be interpreted carefully, and we recommend using these p-values together with the simple count values to have a better understanding.

The table can be sorted by any column. In the above view, we sorted the data by logFC. The table can also be sorted by # of data points to view the genes with most data.  In the example below, we chose 10 up-regulated genes with most data points form the table and used the bubble plot button to create bubble plot. It can be seen that indeed all these genes show consistent up-regulated in the selected comparisons.

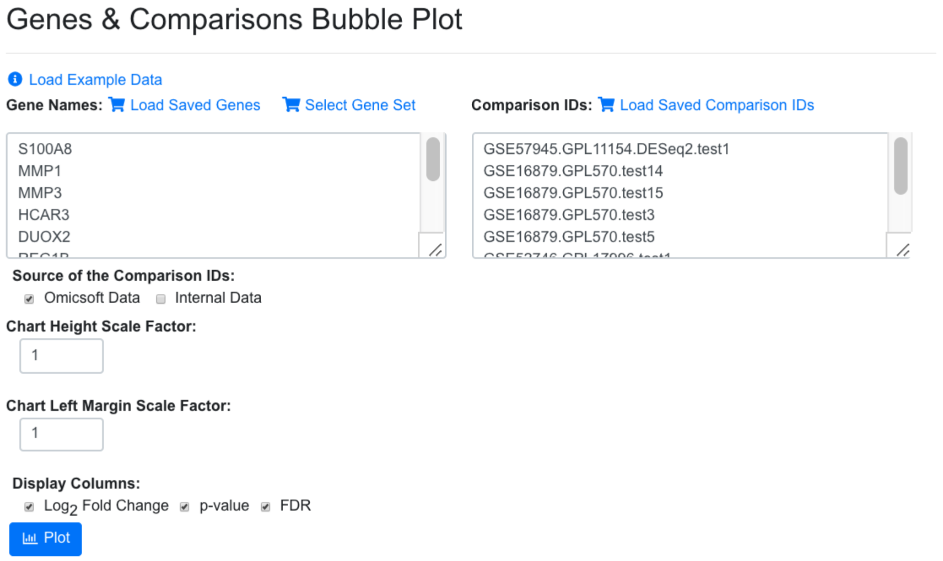

The resulting bubble plot will show all 8 comparisons for each gene.

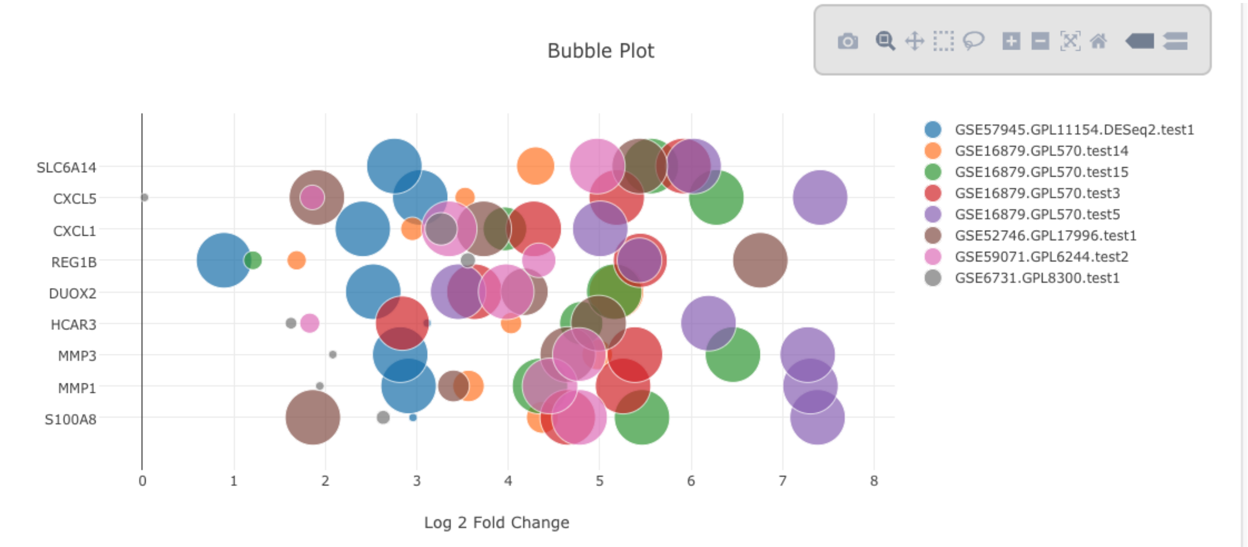

We can also create a gene list from the table for up or down gene lists from meta-analysis (using the green button at top of meta-analysis results) and use the list for subsequent study like functional enrichment or generating heatmap. Below is a heatmap using 100 up-regulated genes from meta-analysis, nd its very clear that the diseased samples have an upregulation of these genes.

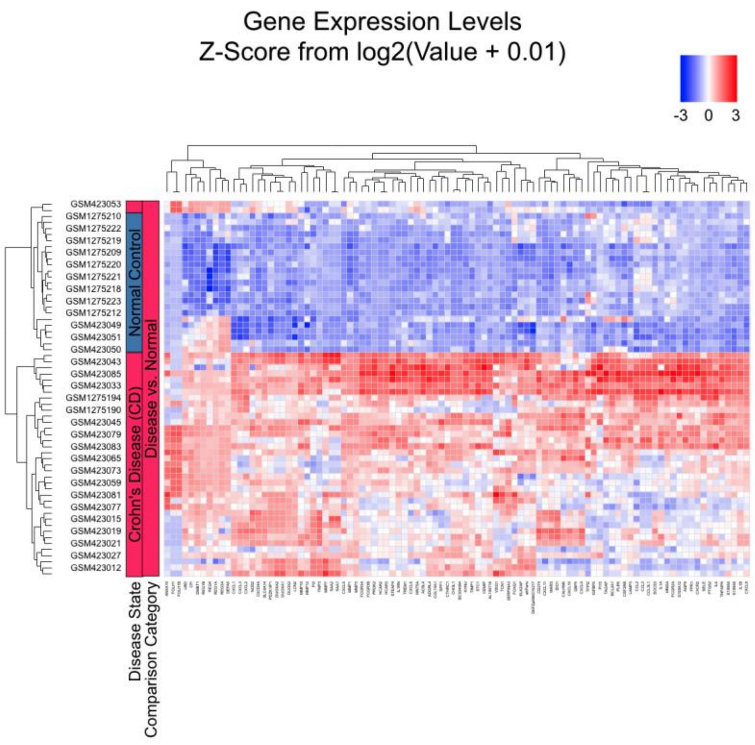

#### 6.3.2 Meta-Analysis (Gene Expression)

In the example below, we use the same comparisons from Crohns disease.

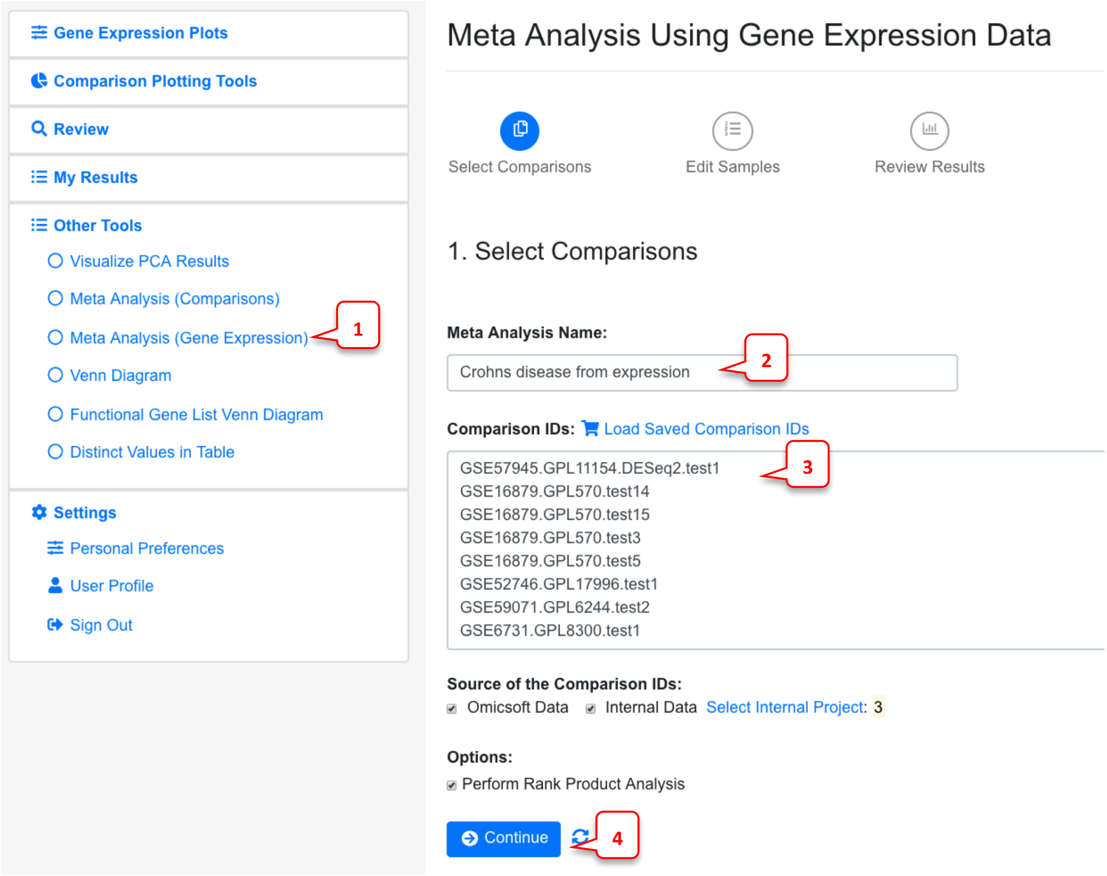

By default, the Rank Product Analysis is checked. Uncheck this if the data is too big for Rank product analysis, or you just need a quick run using limma for each individual comparison.

In the next page, system will display the Case and Control samples for each comparison. At this step, users can edit samples. Users can use this page to remove outliers or add other samples they want to include in the meta-analysis.

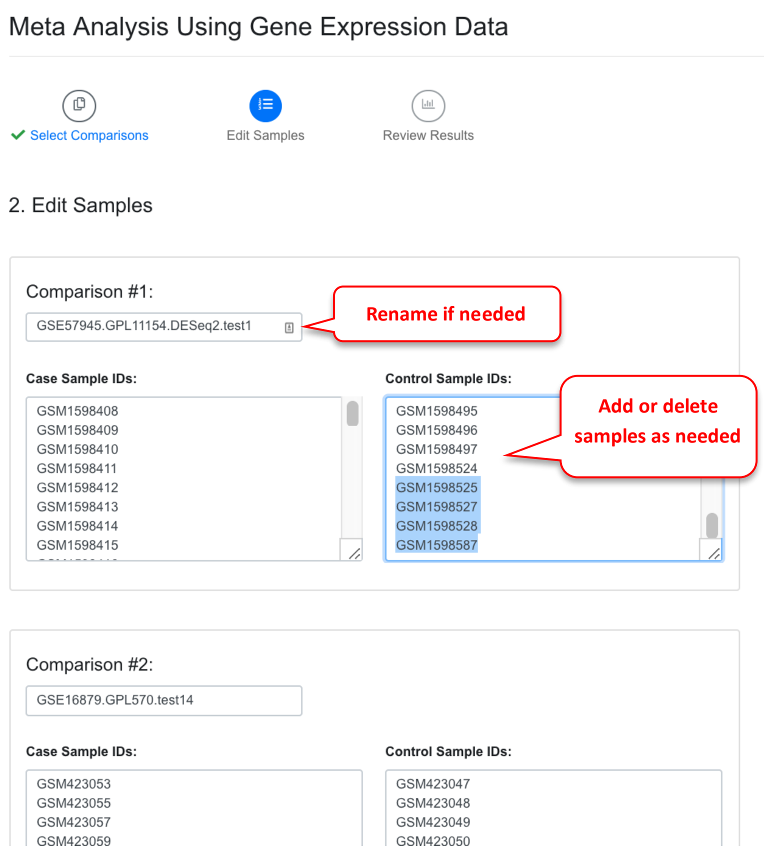

After users review the samples for each comparison, they can go to next step to run the meta-analysis. This step can take a while if the number of samples are large. For the meta-analysis, the system will do the following:

1) Run limma to get logFC, p-valua and confidence interval for each individual comparison
2) Run metaDE to get combined p-values
3) Run Rank Product package for all the expression data to get statistical significance and combined logFC. This step can be very slow is number of samples are large.

After the analysis is done, the system will show a summary and the result table.

#### Meta Analysis Using Gene Expression Data

Create New Meta Analysis

Summary | Meta Analysis Results

##### Meta Analysis Results for Crohns disease meta

Comparisons used in the analysis:

| Comparison Name | Comparison Number | Average Number of Genes for Meta Analysis | Average Number of Not Available (NA) Genes | # of Case Samples | # of Control Samples |
|---|---|---|---|---|---|
| GSE57945.GPL11154.DESeq2.test1 | 1 | 30,373 | 0 | 4 | 4 |
| GSE16879.GPL570.test14 | 2 | 21,893 | 8,480 | 4 | 4 |
| GSE52746.GPL17996.test1 | 3 | 21,893 | 8,480 | 3 | 4 |
| GSE59071.GPL6244.test2 | 4 | 21,562 | 8,811 | 4 | 4 |

View samples in each comparison (Comparison_List.csv) | View number of genes in each comparison (Sample_geneCount.csv)

From **30,373** genes listed in the comparisons, **19,214** genes are present in all comparisons and thus produce statistical results from meta-analysis.

Note: The statistical values from meta analysis are only available for genes that are present in all comparisons. If the number of genes in one comparison is much lower than others, consider running another meta-analysis without this comparison to get statistical values for more genes.

Meta Analysis Results

Download

Show 100 entries

Copy | CSV | Excel | PDF

Search:

| Actions | Symbol | Description | logFC_RP | P.Val_RP | FDR_RP | RankProd | ES_pval | ES_FDR | logFC_Ave | logFC_1 | SE_1 | CI.L_1 | CI.R_1 | P.Value_1 | adj.P.Val_1 | logFC_2 | SE_2 | CI.L_2 | CI.R_2 | P.Value_2 | adj.P.Va |
|---|---|---|---|---|---|---|---|---|---|---|---|---|---|---|---|---|---|---|---|---|---|
| | 5S_rRNA | NA | | | | | | | -0.00345 | 0 | 0.00377 | -0.00908 | 0.00908 | 1 | 1 | | | | | | |
| | 5S_rRNA | NA | | | | | | | -0.00408 | 0 | 0.00377 | -0.00908 | 0.00908 | 1 | 1 | | | | | | |
| | 5S_rRNA | NA | | | | | | | -0.00052 | 0 | 0.00377 | -0.00908 | 0.00908 | 1 | 1 | | | | | | |
| | 5S_rRNA | NA | | | | | | | 0 | 0 | 0.00377 | -0.00908 | 0.00908 | 1 | 1 | | | | | | |
| | 5S_rRNA | NA | | | | | | | 0.00314 | 0 | 0.00377 | -0.00908 | 0.00908 | 1 | 1 | | | | | | |
| | 5S_rRNA | NA | | | | | | | -0.46437 | -1.24584 | 0.40456 | -2.22073 | -0.27095 | 0.01993 | 0.30861 | | | | | | |
| | 5S_rRNA | NA | | | | | | | 0 | 0 | 0.00377 | -0.00908 | 0.00908 | 1 | 1 | | | | | | |
| | 5S_rRNA | NA | | | | | | | 0.00445 | 0 | 0.00377 | -0.00908 | 0.00908 | 1 | 1 | | | | | | |
| | 5S_rRNA | NA | | | | | | | 0.00029 | 0 | 0.00377 | -0.00908 | 0.00908 | 1 | 1 | | | | | | |
| | 5S_rRNA | NA | | | | | | | -0.00108 | 0 | 0.00377 | -0.00908 | 0.00908 | 1 | 1 | | | | | | |
| | 5S_rRNA | NA | | | | | | | 0 | 0 | 0.00377 | -0.00908 | 0.00908 | 1 | 1 | | | | | | |

In the summary, the four comparisons are shown, along with number of genes used for meta-analysis, and number of case and control samples. Because different platforms (especially arrays) have different number of genes present, and some genes may have too low signals to be detected in certain projects, not all genes are present in all the comparisons. In this example, 19,214 genes are present in all comparisons. In some cases, it may be beneficial to remove some comparisons with small number of genes.

In the meta-analysis result table, results from Rank Product are shown (from RankProd package, preferred results to use), followed by effective size method (MetaDE.ES from metaDE package). The logFC, SE, confidence internal, p-value and FDR for each comparison is shown next. The comparison numbers are the same as those listed in the summary table above. The gene list can be sorted by RankProd (most significant changes) or by logFC_RP (up or down-regulated).

From this analysis, we can create Forest plot for each gene by click the icon near the gene symbol.

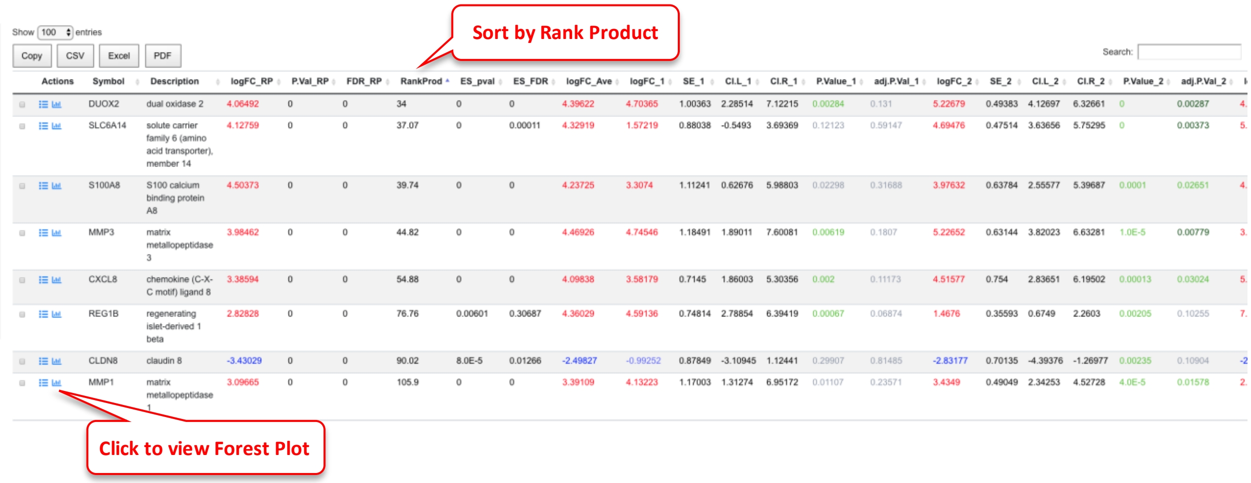

Show 100 entries

Copy CSV Excel PDF

Search:

| Actions | Symbol | Description | logFC_RP | P.Val_RP | FDR_RP | RankProd | ES_pval | ES_FDR | logFC_Ave | logFC_1 | SE_1 | CI.L_1 | CI.R_1 | P.Value_1 | adj.P.Val_1 | logFC_2 | SE_2 | CI.L_2 | CI.R_2 | P.Value_2 | adj.P.Val_2 | l |
|---|---|---|---|---|---|---|---|---|---|---|---|---|---|---|---|---|---|---|---|---|---|---|
| | DUOX2 | dual oxidase 2 | 4.06492 | 0 | 0 | 34 | 0 | 0 | 4.39622 | 4.70365 | 1.00363 | 2.28514 | 7.12215 | 0.00284 | 0.131 | 5.22679 | 0.49383 | 4.12697 | 6.32661 | 0 | 0.00287 | 4. |
| | SLC6A14 | solute carrier family 6 (amino acid transporter), member 14 | 4.12759 | 0 | 0 | 37.07 | 0 | 0.00011 | 4.32919 | 1.57219 | 0.88038 | -0.5493 | 3.69369 | 0.12123 | 0.59147 | 4.69476 | 0.47514 | 3.63656 | 5.75295 | 0 | 0.00373 | 5. |
| | S100A8 | S100 calcium binding protein A8 | 4.50373 | 0 | 0 | 39.74 | 0 | 0 | 4.23725 | 3.3074 | 1.11241 | 0.62676 | 5.98803 | 0.02298 | 0.31688 | 3.97632 | 0.63784 | 2.55577 | 5.39687 | 0.0001 | 0.02651 | 4. |
| | MMP3 | matrix metallopeptidase 3 | 3.98462 | 0 | 0 | 44.82 | 0 | 0 | 4.46926 | 4.74546 | 1.18491 | 1.89011 | 7.60081 | 0.00619 | 0.1807 | 5.22652 | 0.63144 | 3.82023 | 6.63281 | 1.0E-5 | 0.00779 | 3. |
| | CXCL8 | chemokine (C-X-C motif) ligand 8 | 3.38594 | 0 | 0 | 54.88 | 0 | 0 | 4.09838 | 3.58179 | 0.7145 | 1.86003 | 5.30356 | 0.002 | 0.11173 | 4.51577 | 0.754 | 2.83651 | 6.19502 | 0.00013 | 0.03024 | 5. |
| | REG1B | regenerating islet-derived 1 beta | 2.82828 | 0 | 0 | 76.76 | 0.00601 | 0.30687 | 4.36029 | 4.59136 | 0.74814 | 2.78854 | 6.39419 | 0.00067 | 0.06874 | 1.4676 | 0.35593 | 0.6749 | 2.2603 | 0.00205 | 0.10255 | 7. |
| | CLDN8 | claudin 8 | -3.43029 | 0 | 0 | 90.02 | 8.0E-5 | 0.01266 | -2.49827 | -0.99252 | 0.87849 | -3.10945 | 1.12441 | 0.29907 | 0.81485 | -2.83177 | 0.70135 | -4.39376 | -1.26977 | 0.00235 | 0.10904 | -2 |
| | MMP1 | matrix metallopeptidase 1 | 3.09665 | 0 | 0 | 105.9 | 0 | 0 | 3.39109 | 4.13223 | 1.17003 | 1.31274 | 6.95172 | 0.01107 | 0.23571 | 3.4349 | 0.49049 | 2.34253 | 4.52728 | 4.0E-5 | 0.01578 | 2. |

In the Forest plot, results from individual comparison and combined data are shown. The confidence internal for individual comparison is based on limma output. The confidence internal for combined results is based on p-value from RankProd (or ES method when RankProd is not run).

#### Forest Plot for Gene DUOX2

Return to Meta Analysis Results   Review Gene Details

**Meta Analysis:** Crohns disease meta
**Gene Name:** DUOX2
**EntrezID:** 50506
**Description:** dual oxidase 2

**Download Plot**: SVG • PNG • PDF

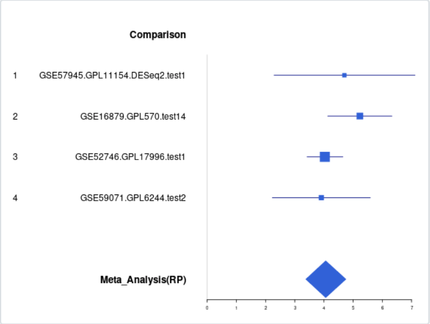

| Comparison Number | Comparison Name | # of Case Samples | # of Control Samples | logFC | CI.L | CI.R | SE | P.Value | FDR |
|---|---|---|---|---|---|---|---|---|---|
| 1 | GSE57945.GPL11154.DESeq2.test1 | 4 | 4 | 4.7 | 2.29 | 7.12 | 1.004 | 2.84e-03 | 1.31e-01 |
| 2 | GSE16879.GPL570.test14 | 4 | 4 | 5.23 | 4.13 | 6.33 | 0.494 | 9.16e-07 | 2.87e-03 |
| 3 | GSE52746.GPL17996.test1 | 3 | 4 | 4.03 | 3.42 | 4.65 | 0.271 | 1.38e-07 | 2.75e-04 |
| 4 | GSE59071.GPL6244.test2 | 4 | 4 | 3.91 | 2.23 | 5.59 | 0.749 | 4.49e-04 | 4.69e-02 |
| NA | Meta_Analysis(RP) | 15 | 16 | 4.06 | 3.38 | 4.75 | 0.349 | 8.53e-29 | 1.64e-24 |

The plot can be saved as SVG or PDF. The summary table is shown below the plot.

### 7 Methods

#### 7.1 Data processing

Public datasets

The public datasets that are accessible through OmicsView are a subset of the DiseaseLand https://www.qiagenbioinformatics.com/diseaseland/ database. The DiseaseLand data service uses common analysis pipelines to quantify and normalize publicly available microarray and RNA-seq expression data from raw files. For each project, and each sample, metadata are curated to apply controlled vocabularies and ensure consistent formatting of terms. Information on the data processing pipeline and DiseaseLand product is available at the following links ( http://www.arrayserver.com/wiki/index.php?title=DiseaseLand_Curation_Pipeline, http://www.arrayserver.com/wiki/index.php?title=Omicsoft_Affymetrix_Microarray_Preprocessing, http://www.arrayserver.com/wiki/index.php?title=RNA-Seq_Normalized_FPKM_Values_in_Land ). The data in OmicsView was exported from DiseaseLand prior to 8/27/2019. No OmicsView data can be exported from the portal, distributed with a software package, or otherwise redistributed without express permission from QIAGEN.

#### 7.2 Functional and Pathway Enrichment

The function enrichment algorithms implemented in OmicsView come in two flavors – one is based on hypergeometric enrichment of differentially expressed genes (DEGs) against a number of knowledge databases, based on the HOMER package (http://homer.salk.edu/homer/microarray/go.html). This requires a significance cutoff to be applied to the two group comparison expression data such that we can select a set of genes that is upregulated and downregulated, i.e. differentially expressed genes. The HOMER package comes with some collections of gene knowledge databases that we use for enrichment.

The second approach is based on a variant of the Gene Set Enrichment Analysis (GSEA) approach called PAGE (Parametric Analysis of Gene Set Enrichment) which is faster than the original GSEA approach and more sensitive[1]. We use an implementation of PAGE that is in the R piano package (https://www.bioconductor.org/packages/release/bioc/html/piano.html). GSEA based approaches just require a ranked list of genes (genes with a numeric quantity for ranking them that shows how different they are between groups in the comparison, usually logFC values) and are typically more sensitive in picking up pathway enrichment. Both packages come pre-built with a set of known functions/pathways in the form of gene sets, against which we look for enrichment in the comparison of interest.

##### 7.2.1 HOMER workflow

###### *7.2.1.1 Selection of Differentially Expressed Genes (DEGs) for enrichment analysis*

For each comparison, we generated the up- and down-regulated gene lists as input for functional enrichment analysis. We used a dynamic cutoff of logFC and AdjustedPValue / PValue to aim for 200-

2000 genes in each list. To achieve this we start with a stringent cutoff of adjusted p-value of 0.05, and 2-fold change up or down, if we get more than 200 genes we use these, otherwise we drop to nominal p-value of 0.01 (with 2-fold up or down), and if we do not reach 200 we drop again to 1.2 fold up or down with adjusted p-value of 0.1. If the most lenient cutoff does not generate a list of 50 up or down genes, we take the top 50 logFC (negative and positive) to generate the lists for enrichment analysis.

###### *7.2.1.2 Functional Enrichment of DEG list against genome*

The findGo.pl program from Homer package ( http://homer.salk.edu/homer/microarray/go.html ) is used to analyze the functional enrichment for each DEG list.

There are several different "ontologies", or libraries of gene groupings that came with the Homer package. We used Homer version v4.8.3, human-o v5.8 and mouse-o v5.8 libraries (current as of Dec 6 2016). These include

Gene Ontology – Biological Process, Molecular Function, Cellular Component

Chromosome Location: Genes with similar chromosome localization (NCBI Entrez Gene)

KEGG Pathways: Groups of proteins in the same pathways (From KEGG)

Protein-Protein Interactions: Groups of proteins interacting with the same protein (From NCBI Entrez Gene)

Interpro: Proteins with similar domains and features (Interpro)

Pfam: Proteins with similar domains and features (Pfam)

SMART: Proteins with similar domains and features (SMART)

Gene3D: Proteins with similar domains and features (Gene3D Database)

Prosite: Proteins with similar domains and features (Prosite Database)

PRINTS: Proteins with similar domains and features (PRINTS Database)

MSigDB: Lists of genes maintained by the Molecular Signature Database (includes many different categories of genes (MSigDB)

BIOCYC: Groups of proteins in the same pathway (NCBI Biosystems/BIOCYC)

COSMIC: Human proteins that are mutated in the same cancers (COSMIC)

GWAS Catalog: Human genes with risk SNPs identified in their vicinity for the same disease (GWAS Catalog)

Lipid Maps: Mouse proteins found in the same lipid processing pathways (NCBI Biosystems/LIPID MAPS)

Pathway Interaction Database: Proteins in the same pathway (NCBI Biosystems/PID)

REACTOME: Proteins in the same biochemical pathways (NCBI Biosystems/REACTOME)

SMPDB: Proteins in the same pathway (SMPDB)

Wikipathways: Protein in the same pathway (Wikipathways)

The HTML output file from HOMER is further enhanced by a custom php script to make it more user friendly.

##### 7.2.2 Gene Set Enrichment Analysis (GSEA) workflow (PAGE)

For our second enrichment tool, we used a variation of the GSEA method, PAGE (Parametric Analysis of Gene Set Enrichment)[1] to process the comparison data as PAGE is much faster than GSEA and also more sensitive.

For each comparison, we produce a rank file with gene symbol and logFC values. If a gene symbol appears multiple times in the same comparison, the average logFC is used.

Human gene sets are downloaded from MsigDB (http://software.broadinstitute.org/gsea/msigdb). Version 5.2 (msigdb.v5.2.symbols.gmt).

Mouse gene sets are downloaded from Bader Lab from Univ. of Toronto (http://baderlab.org/GeneSets). The version is December_01_2016 (Mouse_GO_AllPathways_with_GO_iea_December_01_2016_symbol.gmt) . This gmt file contains special characters that cannot be used by R piano package, therefore we manually replace special characters to / or _ . In addition, some mouse gene sets have the same name, so we added suffix _altSetX to make all the names unique.

###### *7.2.2.1 PAGE Analysis using R piano package*

The R package piano (https://bioconductor.org/packages/release/bioc/html/piano.html) is used to run the PAGE analysis for all the rank files. To simplify the piano output, we combined down-regulated and up-regulated gene sets into a single table for each gene set. From the piano results we extract out the p-value, FDR and Z-score.

#### 7.3 Meta-Analysis

The meta-analysis functions allow a user to combine expression data across multiple studies to find changes that are robust. DiseaseAtlas offers two ways to perform meta-analysis:

The first method works on comparison data. The system will use the comparison data (logFC, p-value) to compute combined p-value and rank product. This method is fast and can be applied to any type of comparison data. However, it does not use the individual sample data, nor does it consider the number of samples in each comparison.

The second method uses per sample Gene Expression data. The user will have input a list of factors that indicate comparisons across or within studies and then gene level significant changes are recomputed by extracting expression data from all samples for each comparison, and then applying the RankProd[2] and/or metaDE[3] packages to perform meta-analysis. Limma is also applied to get statistics for each individual comparison. This analysis takes much longer (10 minutes to an hour for a typical analysis, even longer if number of samples are very large), and it has more strict sample requirements (no samples can occur in two different comparisons).

Statistically, the second method is more robust. For consistently changed genes, both methods should detect them.

##### 7.3.1 Meta-analysis statistics

For comparison data the combined p-value is computed using Fisher's method, that is -2*(sum of ln(p-value)) is compared against a Chi-squared distribution with N degrees of freedom, where N is the number of p-values being combined. This is carried out for every gene and yields a combined p-value that is reported. Another simpler approach is to report the maximum p-value for the gene across all the comparisons – a much more stringent measure of overall significance. This approach to combining p-values is implemented in the MetaDE R package (https://github.com/metaOmics/MetaDE)[3]. Note MetaDE will not produce results if > 30% of the comparisons for a gene have missing values for the p-value. Note that the p-value combination does *not* take into account the direction of the fold change (up or down) so the Up regulated & Down regulated percentage summaries need to be also referred to for clarification.

The RankProd method (https://bioconductor.org/packages/release/bioc/html/RankProd.html) converts log2 fold changes across all genes in a comparison to ranks and then computes a meta-statistic per gene which is the geometric mean of the ranks across comparisons. It is a non-parametric approach, and computes statistical significance based on a permutation approach which also addresses the multiple testing aspect of looking for significance within the set of all genes in the transcriptome.
